# Extending the Martini coarse-grained forcefield to N-glycans

**DOI:** 10.1101/2020.05.08.085399

**Authors:** Aishwary T. Shivgan, Jan K. Marzinek, Roland G. Huber, Alexander Krah, Richard H. Henchman, Paul Matsudaira, Chandra S. Verma, Peter J. Bond

## Abstract

Glycans play a vital role in a large number of cellular processes. Their complex and flexible nature hampers structure-function studies using experimental techniques. Molecular dynamics (MD) simulations can help in understanding dynamic aspects of glycans if the forcefield (FF) parameters used can reproduce key experimentally observed properties. Here, we present optimized coarse-grained (CG) Martini FF parameters for N-glycans, calibrated against experimentally derived binding affinities for lectins. The CG bonded parameters were obtained from atomistic (ATM) simulations for different glycan topologies including high mannose and complex glycans with various branching patterns. In the CG model, additional elastic networks are shown to improve maintenance of the overall conformational distribution. Solvation free energies and octanol-water partition coefficients were also calculated for various n-glycan disaccharide combinations. When using standard Martini non-bonded parameters, we observed that glycans spontaneously aggregated in the solution and required down-scaling of their interactions for reproduction of ATM model radial distribution functions. We also optimised the non-bonded interactions for glycans interacting with seven lectin candidates and show that scaling down the glycan-protein interactions can reproduce free energies obtained from experimental studies. These parameters should be of use in studying the role of glycans in various glycoproteins, carbohydrate binding proteins (CBPs) as well as their complexes, while benefiting from the efficiency of CG sampling.

## Introduction

Glycosylation is a key post-translational modification involved in a wide range of cellular processes including host-pathogen interactions^1,2^, cell trafficking^3^, fertilization^4^, immune system function^5^, energy storage^6–8^, and are associated with disease states such as congenital disorders including cellular transport defects^9^, muscular dystrophies^10–12^ as well as cancer progression^13^. Glycans play a major role in folding and stability of glycoproteins^14^. They are one of the key parameters to be considered while developing therapeutic antibodies^15–17^. Glycans bind to carbohydrate recognition domains (CRDs) of lectins^18^ with low affinity^19^, giving cells a versatile system for carbohydrate-protein recognition. They are made up of monosaccharides which can form a variety of anomeric ring linkages resulting in different structures and associated specificities for diverse receptors. These structures range from e.g. cellulose, which is a linear polymer, to cyclodextrins^20^, a cyclic polymer. Branching of glycans gives them an overall tertiary structure and in turn contributes to the quaternary structures of glycoproteins^21^.

Glycans are classified according to their protein attachment sites. N-linked glycans are covalently bound to asparagine (N) at the NxS/T motif, where x can be any amino acid apart from proline (P). The other major type of glycans are called O-linked glycans due to their attachment to the hydroxyl oxygen atom of serine (S) or threonine (T) residues^21,22^. Depending upon the sugars (mannose (Man), n-Acetylglucosamine (GlcNAc) and galactose (Gal)) that extend the common core sequence, Man-α(1,6)-(Man-α(1,3)-Man-β(1,4)-GlcNAc-β(1,4)-GlcNAc-β1-N-X-S/T, N-glycans are classified into three classes. In the first class, the high mannose (oligomannose) type, the core is extended only by mannose sugars. The second is the complex class, in which branches are extended by N-acetylglucosaminyltransferases (GlcNAcTs). The third class includes the hybrid glycans, in which the Man-α(1,6) arm is attached only to mannose sugars while either one or two complex branches are attached to the Man-α(1,3) arm^21,22^. Even though there are only three classes of these glycans, the number of glycans found in each class is numerous, hampering structure-function studies. The complexity of the multistep glycosylation pathways very often results in multiple glycoforms for each glycoprotein^23^. Also, the inherently flexible nature of glycosidic linkages typically makes it difficult to define their precise structure by either X-ray crystallography or NMR spectroscopy beyond a few monosaccharide units^24^. The requirement of highly purified samples further complicates NMR studies^23^. Very often glycoproteins are deglycosylated in order to reduce the microheterogeneity and associated surface entropy in an attempt to obtain higher resolution crystal structures^25^. Mass spectrometry can provide structural data for small glycans but is harder for larger complex glycans due to difficulties in determining the glycosidic linkages^26,27^. Hence, even though glycans are biologically very important, rapid experimental characterization of their structure and delineation of their functional roles remains a major challenge.

The gaps in understanding the role of glycans at the molecular level and their potential impact on biological processes can potentially be filled by computational modelling, and in particular, by the use of molecular dynamics (MD) simulations. The precise dynamic, biophysical, and thermodynamic *in silico* properties of polysaccharides that can mimic experimental observations depend upon their representation and parameters defined within the FF. Several carbohydrate-specific FFs have been developed in recent years^28–31^, the choice of which depends upon the application and simulation conditions desired. The MM3^32^ FF is useful for reproducing gas phase potential energy curves while the SPASIBA^33^ FF is designed to reproduce infrared and Raman spectroscopy data. Other FFs such as AMBER^34^, CHARMM^35^, GROMOS^36^, and GLYCAM^28^ are good choices for simulating solvated systems but might not be able to reproduce crystal-phase infrared data (see e.g.^33,37^). Excellent web based tools such as CHARMM-GUI^38,39^ and GLYCAM-Web^40^ make it easier to generate input data for glycan simulations and have been employed in studies that seek to understand, for example, their interactions with other biomolecules as well as their dynamics in different environments^41,42^. Many of these studies were carried out using ATM representations, which can limit the accessible time scales that may be reached. Biologically relevant complexes containing glycans, such as antibodies, carbohydrate recognition domains like DC-SIGN, or mannose receptors, can encompass millions of atoms, thus making these calculations very expensive and limiting time scales to the sub-microsecond regime, i.e. not equivalent to those sampled in key biological processes or biophysical experiments that reach microseconds to milliseconds or beyond^43^. Alternatively, a CG representation, in which groups of atoms are represented as larger beads, can be helpful in overcoming the associated limitations, by reducing the number of degrees of freedom and simplifying the energy landscape.

The Martini FF^44^ is one of the most widely used CG models for biomolecular systems. Martini was originally developed for lipids, and was later extended to proteins^45^, carbohydrates^46^, and nucleic acids^47^. In the Martini representation, approximately four heavy atoms are grouped into a single bead. This represents a relatively lighter coarse-graining approach which allows maintenance of the key structural details of biomolecules. In Martini, non-bonded parameters for different particles have been calibrated against partitioning free energies of small compounds in polar and apolar solvents^44^. The bonded parameters are typically derived empirically by comparing the distributions with ATM simulations. An increasing number of studies have shown that there is an imbalance in the non-bonded interactions, making Martini (v2.2) too “sticky” which has necessitated fine-tuning of the parameters^48–50^. Recently, an open beta version of Martini v3.0b was released for phospholipids and proteins^51^. This version of Martini adds more bead types with various interaction types that aims to solve the shortcomings in the Martini v2.2 FF^48–50^.

The Martini FF has been extended to carbohydrates^46^ and includes the parameters for monosaccharides such as glucose and fructose and disaccharides like sucrose, trehalose, maltose, and cellobiose, whose particles have been calibrated to reproduce water-octanol solvation and partitioning energies. The application of these parameters to oligosaccharides such as amylose and curdlan reproduced their key structural properties^46^. Nevertheless, the FF still lacks bonded parameters for different types of glycosidic linkages as well as branching patterns specific to N-glycans such as trimannose (Man-α(1,6)-[Man-α(1,3)-]Man) and bisected N-glycans^14^. In addition, parameters are not presently available for N-Acetylglucosamine (GlcNAc), Fucose (Fuc), and Sialic acid (Neu5Ac), which are very common building blocks in many of the N-glycans found in glycoproteins. Hence, there is a gap in the availability of parameters covering the variety of linkages and topologies needed to model biologically relevant glycans, as well as in reproducing experimentally observed glycan-protein binding affinities and aggregation properties.

In this work, we have extended the Martini FF to N-glycans. As there are many possible glycans, we have restricted our parameterization to the most commonly found N-glycans at present. The bonded parameters for glycans with different glycosidic linkages and branching patterns were obtained by comparing them to ATM simulation data. Elastic networks were shown to be useful in maintaining the conformations of longer glycans. We also observed that the CG glycans tend to aggregate in solution, as in previous studies^48^. Solvation and partitioning coefficients were calculated and compared against prediction methods such as ClogP and KOWWIN^52^. Binding free energies of glycans to lectins, obtained from umbrella sampling (US) calculations were overestimated for all the glycans, confirming a requirement for the fine tuning of non-bonded parameters. This was achieved by scaling non-bonded interactions and comparing the data to binding free energies, radial distribution functions, as well as second virial coefficients (B_22_)^53^. We found that relatively modest scaling helped to reproduce solution behaviour of glycans only systems and most of the experimental binding affinities in the case of seven candidate lectins.

## Methods

### All atom simulations

The GROMOS54a7^54^ united atom (UA) FF was used for various α and β glycosidic linkages with different monosaccharides including D-glucose (Glc), D-mannose (Man), and D-galactose (Gal). D-Glucose parameters were used for the glucose unit of the n-acetyl-D-glucosamine (GlcNAc). The bonded parameters for GlcNAc-β1-asparagine (N-) connections were derived from the extended GROMOS53a6_GLYC_^31^ FF for glycoproteins. D-fucose (Fuc) and D-sialic acid (Neu5Ac) were manually prepared based on the corresponding galactose and mannose structures, respectively. Different types of ATM N-Glycan structures including disaccharides, trisaccharides (Figure 1) and full length glycans (Figure 2) were constructed using the GLYCAM carbohydrate builder^40^. Each structure was placed in a cubic box such that any atom was at least 1 nm away from any wall of the simulation box to avoid self-interaction. The molecules were then energy minimized for ≤ 2000 steps using the steepest descent (SD) algorithm with a 0.01 nm step size^55^. The simulation box was solvated with explicit SPC water molecules^56^ and then again energy minimized using SD. Ions were added to neutralize the overall system charge. The systems were equilibrated for 5 ns in the NPT (constant number of atoms, pressure and temperature) ensemble. The production runs were performed for 1000 ns and convergence was assessed by performing block analysis on the bond, angle, and dihedral distributions. The simulations were run using the velocity rescale thermostat with an additional stochastic term^57^ at a temperature of 310 K with a relaxation time of 0.1 ps. The Berendsen barostat^58^ was used to maintain the pressure at 1.0 bar with weak coupling using a relaxation time of 1 ps. All bonds to hydrogen atoms were constrained using the LINCS^59^ algorithm with a relative geometric tolerance of 10^−4^ enabling a time step of 2 fs to integrate Newton’s equations of motions with the leap-frog algorithm. A short-range cut-off of 1.2 nm was used for electrostatics and van der Waals interactions. The Particle Mesh Ewald (PME)^60^ method was used for long-range electrostatics, with a 1.2 nm real space cutoff. In addition to GROMOS54a7, comparative simulations using the CHARMM36m^61^ FF with the TIP3P water model were used to assess the solution behaviour of glycans, in which case similar conditions were applied along with an additional force switch smoothing function from 1.0 to 1.2 nm for van der Waals interactions. Atomic coordinates were saved every 0.1 ns. All AA simulations were run using the GROMACS 5.1.2^62^ package on an in-house Linux cluster as well as on the ASPIRE 1 supercomputer of the National Supercomputing Centre Singapore (NSCC).

**Figure 1:**
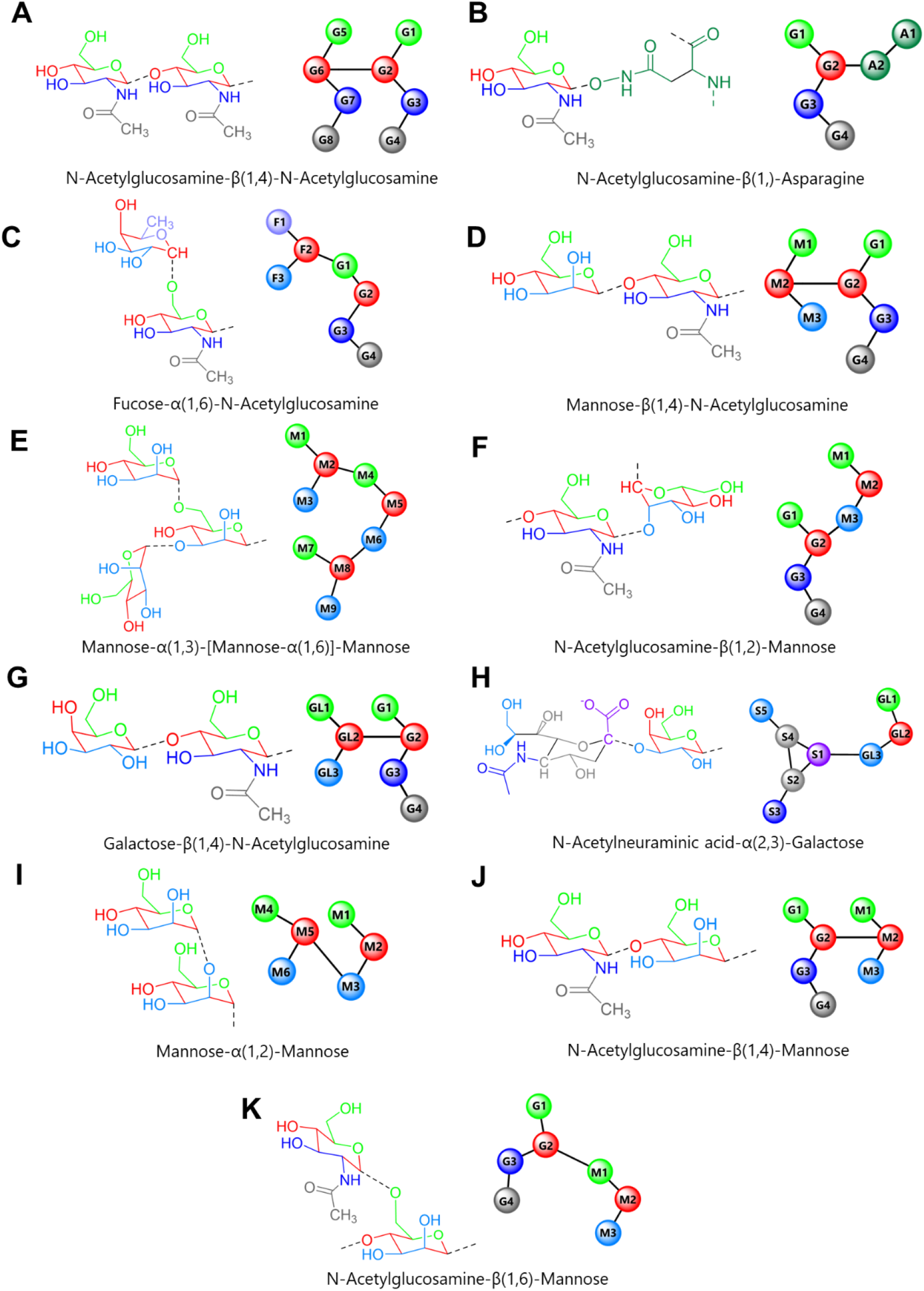
Disaccharide/trisaccharides used for developing parameters for N-glycans. Each image shows the atomistic representation of the saccharide (left) with mapped martini representation (right). The atoms which are mapped together are shown with the same colour as the beads in the CG model. The parameters for these di/trisaccharides are summarised in Table S1.

**Figure 2:**
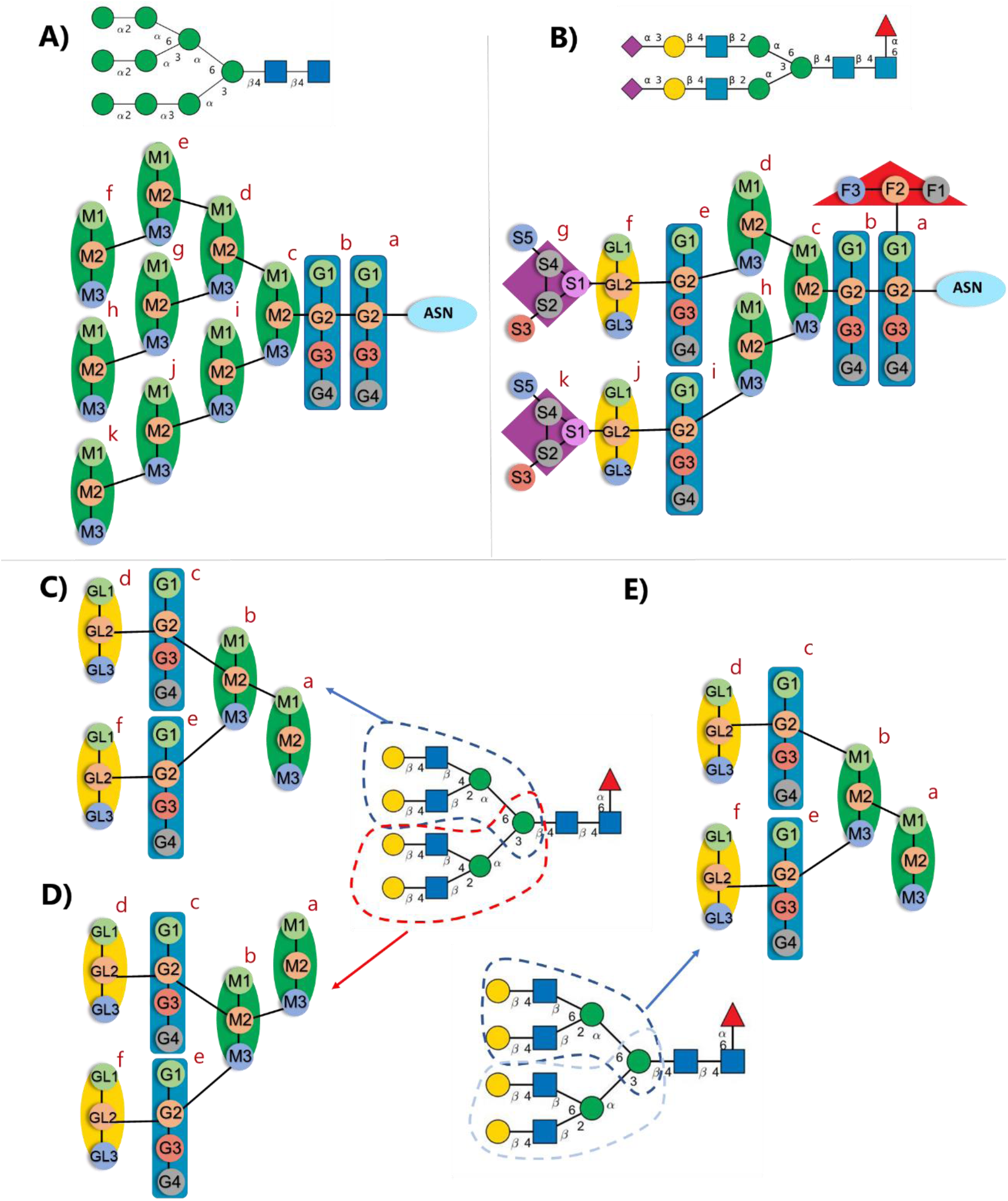
Parametrization of N-glycans constructed from disaccharides: (A) high mannose (M9) glycan; (B) sialylated bi-antennary (FA2G2S2) complex glycan; and (C) – (E) parts of tetra-antennary complex glycans parameterized separately for various linkages shown with dashed lines. All bonded parameters required to maintain the conformation of the glycans are summarised in the Table 1. Monosaccharides present in the glycans are represented by their symbolic representation: mannose (green circle), N-acetylglucosamine (blue square), galactose (yellow circle), and Neu5Ac/sialic acid (purple diamond).

### CG simulation setup

We followed the Martini mapping scheme for simple monosaccharides and unbranched oligosaccharides as suggested for carbohydrates^46^. In this mapping scheme, each sugar is modelled using three beads and the glycosidic bonds are made by connecting the central beads of two monosaccharides, regardless of the type of glycosidic linkage. However, this is harder to implement in the case of glycans with heavy branching where as many as four monosaccharides are attached to one monosaccharide, such as in bisected tetra-antennary complex glycans. Hence, for these heavily branched glycans, we employed a slightly different method for connecting the monosaccharides. Figure 1 shows the typical di/trisaccharide combinations observed in N-glycans. In the case of Fuc-α(1,6)-GlcNAc, the α(1,6) bond was represented by linking the 2^nd^ and 5^th^ beads. For trimannose Man-α(1,6)[Man-α(1,3)]-Man, the α(1,3) bond was represented by linking the 3^rd^ and 4^th^ beads and the same was done for other types of glycosidic linkages (Figure 1). The bonds were implemented between beads containing atoms originating from the glycosidic linkage in their ATM counterparts. All monosaccharides were linked using the above strategy for all the glycosidic bonds including α/β(1/2, 2/3/4/6) connections. The bead types used for most of the monosaccharides were chosen as suggested in the original carbohydrates Martini CG study^46^ and slightly modified depending upon the new mapping scheme (Figure 1). P1, SP2 and P4 polar beads were used for monosaccharides with a three bead model such as Man and Gal. Fuc, GlcNAc and Neu5Ac models were not previously available, so a five bead model was used to represent Neu5Ac using Qa, SP1, P4 and P5 beads. Fuc was parameterized using SP1, P2 and P4 beads. A four bead model was used for GlcNAc in which P1, SP2, P3 beads were used to model the core sugar while SP1 was used to model the acetyl group. The beads for the Fuc, GlcNAc and Neu5Ac were chosen based on chemical intuition and by analogy with previously parametrised carbohydrate-like molecules^46,63^, and further validated by calculating their partitioning data (see below). The mapping schemes are shown in Figure 1, while bead type selections for each of them are given in supplementary Table S2.

**Table 1:** Extra angles and elastic network parameters for N-glycans shown in Figure 2. Each bead is defined by its name as shown in Figure 2; in some cases, a superscript is used when there are two or more similar types of connections. i.e. G2a is a G2 bead belonging to the ‘a’ typed sugar in Figure 2. For angles greater than 140°, the restricted bending potential (ReB) was used.

| Glycan | Elastic Bonds | Rbond (nm) | Kbond (kJ mol<sup>-1</sup> nm<sup>-2</sup>) | Angles | Θ (°) | Kangle (kJ mol<sup>-1</sup>) |
| --- | --- | --- | --- | --- | --- | --- |
| High-Mannose (M9) (A) | G2a-M2f | 1.36 | 75 | A2ASN-G2a-G2b | 172 | 110 |
|  | G2a-M2h | 1.03 | 75 | G2a-G2b-M2c | 160 | 40 |
|  | G2a-M2k | 1.90 | 75 | M2f-M2d-M2h | 120 | 15 |
|  |  |  |  | M2f-M2c-M2k | 78 | 10 |
|  |  |  |  | M2h-M2c-M2k | 105 | 10 |
| Biantennary complex glycan (FA2G2S2) (B) | M2c-GL2f | 1.32 | 200 | A2ASN-G2a-G2b | 172 | 110 |
|  | M2c-GL2j | 1.35 | 200 | G2-G2-M2 | 160 | 40 |
|  |  |  |  | M3-G2-GL2 | 170 | 95 |
|  |  |  |  | G2-GL2-S1 | 112 | 30 |
|  |  |  |  | GL2-M2c-GL2 | 125 | 30 |
| Tetraantennary complex glycan (FA4G4) (C) | M2a-GL2d | 1.43 | 300 | M1a-M2b-G2c | 130 | 65 |
|  | M2a-GL2f | 1.32 | 200 | M2-G2-GL2 | 160 | 170 |
|  |  |  |  | M3-G2-GL2 | 178 | 70 |
| Tetraantennary complex glycan (FA4G4) (D) | M2a-GL2d | 1.43 | 300 | M3a-M2b-G2c | 138 | 120 |
|  | M2a-GL2f | 1.32 | 200 | M2-G2-GL2 | 162 | 170 |
|  |  |  |  | M3-G2-GL2 | 178 | 70 |
| Tetraantennary complex glycan (FA4G4) (E) | M2a-GL2d | 1.41 | 350 | M1-G2-GL2 | 166 | 180 |
|  | M2a-GL2f | 1.35 | 420 | M3-G2-GL2 | 178 | 50 |

### CG simulation parameters

The Martini FF^44,46,64^ was used for all the CG simulations performed in this study. ATM glycans were mapped according to the representation in Figure 1. Each di/tri-saccharide system was prepared similarly to the ATM system. Systems were solvated with Martini water beads and 10% antifreeze particles. Ions were added to neutralize the overall system charge before energy minimization using SD. A time step of 10 to 20 fs was used to integrate the equations of motion using the leap frog algorithm. A constant temperature of 310 K and a constant pressure of 1 bar were maintained, via the velocity rescale thermostat^65^ and the Parrinello-Rahman barostat^54^ with relaxation times of 1 ps and 12 ps, respectively. The non-bonded interactions were truncated at a distance cut-off of 1.1 nm. Electrostatics were handled using a reaction-field^66^ with a cut-off of 1.1 nm. Production runs were carried out for 1 µs with coordinates saved every 0.2 ns. These parameters were directly taken form the suggested mdp file settings available on the Martini website (http://cgmartini.nl/images/parameters/exampleMDP/martini_v2.x_new-rf.mdp).

### Parameterization of CG bonded interactions

Three types of bonded harmonic potentials were used. The potential *V*_*bond*_*(r)* was used to describes the bonds between the CG particles using:

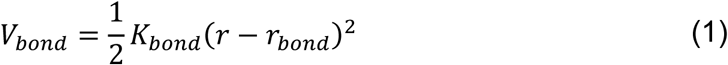

where *r*_*bond* and_ *K*_*bond*_ are the equilibrium distance and the force constant, respectively. A harmonic potential for angles was used for three consecutive beads:

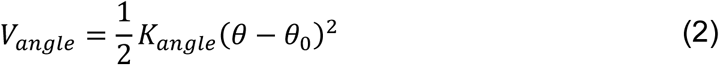

where *ϴ*_*0*_ and *K*_*angle*_ are the equilibrium angle and force constant, respectively. When the angle was found to be greater than 140°, the restricted bending potential (ReB) was used in order eliminate numerical instabilities associated with dihedral angles:

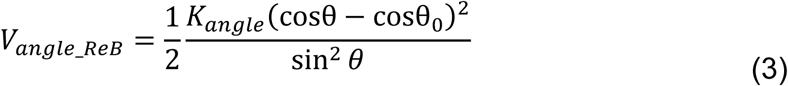

Dihedrals were described using the function:

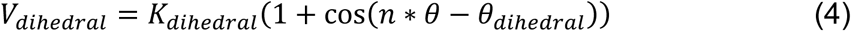

where *ϴ*_*dihedral*_ is the equilibrium angle between planes defined by the coordinates of the atoms *i, j, k* and *j, k, l* respectively, *K*_*dihedral*_ is the force constant, and *n* is the multiplicity. Most of the dihedrals with a single minimum were fitted using a multiplicity of 1, while those with two minima were fitted using a multiplicity of 2.

All the ATM trajectories for the systems in Figure 1 were converted to pseudo CG trajectories. Bonds, angles and dihedral distributions were obtained from these pseudo-CG ATM based trajectories and converted into potentials using the Boltzmann inversion method, and fitted with the respective bonded potential functions. CG simulations with these potentials were run and manually fine-tuned in an iterative fashion until they matched as closely as possible to the ATM distributions. Ultimately, these parameters were averaged for the molecules having the same types of bonds and angles within the same disaccharide (eg. GlcNAC-β(1,4)-GlcNAC) or different disaccharides (eg. Man-β(1,4)-GlcNAC and GlcNAC-β(1,4)-GlcNAC) since we observed a maximum of only 5-10% variation between them. The parameters for all di/tri-saccharides (Figure 1) are shown in Table S1 and the ATM vs CG distributions are shown in Figure S1.

### CG non-bonded interactions

In Martini, the van der Waals component of the non-bonded interactions is described by the Lennard Jones (LJ) 6-12 potential energy function:

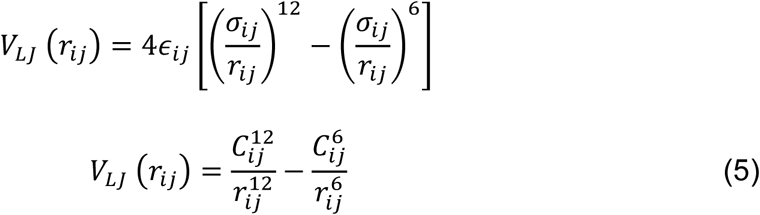

where σ_*ij*_ is the distance at which the potential crosses zero and epsilon ϵ_ij_ is the well depth. There are a total of 19 different bead types. Beads are divided into four categories according to their ϵ values: polar (P), nonpolar (N), apolar (C) and charged (Q). Each main type of bead is subdivided by its hydrogen bonding properties such as donor (d), acceptor (a), both (da) and none (0). The polarity of the bead ranges from low (=1) to high (=5) with interaction level (ε) ranging from 2.0 to 5.6 kJ mol^−1^ and an interaction distance (σ) of 0.47 nm. Smaller beads are used for ring structures with σ=0.43nm and 75% of the normal ε value. These values were previously parameterized to reproduce partition free energies of a library of small molecules^44^. In order to optimise the non-bonded interactions for our glycans of interest, we changed the well depth of the LJ potential i.e. modified the value of ϵ using the following equation,

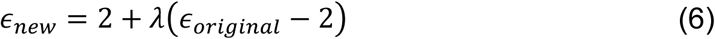

where *λ* is a scaling factor ranging from 0 to 1, whilst the value of ϵ remains unchanged when λ=1.0 and becomes 2 kJ/mol when λ=0, corresponding to the lowest value of ϵ for a bead in the Martini FF. A similar approach has been used in other studies to correct the non-bonded FF parameters^48,49^. Only the solute-solute (glycan-glycan or glycan-protein) interactions were scaled down while solute-solvent and solvent-solvent interactions were kept at their default level. This was done by adding special glycan bead types, and rescaling (by λ) the C^6^ and C^12^ terms (Equation 5) for their interaction with other solute particles accordingly. The down-scaling of glycan-glycan/glycan-protein interactions effectively makes the glycan-water interactions more favourable, consistent with experiments analysing the second virial coefficient (B_22_)^48^.

### Partitioning free energies

The choice of bead types used in Figure 1 was validated by calculating partitioning propensities. Solvation free energies of various disaccharides in the water (Δ*G*_*W*_) and octanol (Δ*G*_*O*_) phases were used for calculating partition coefficients (log *P*_*OW*_). The free energies of solvation Δ*G*_*W*_ and Δ*G*_*O*_ were calculated using thermodynamic integration^67^:

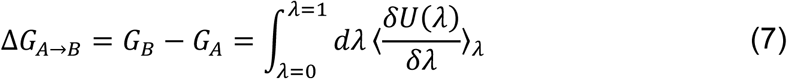

where the potential energy change (*δU*) for going from state A to B is calculated as a function of coupling parameter (*λ*) which goes from 0 (full interactions between beads and solvent) to 1 (no interaction). Non-bonded interactions were scaled linearly. A soft core potential was used to circumvent potential singularities which occur during annihilation or creation of atoms^68^. A total of 55 intermediate *λ* values were used, including additional ones in the high curvature regions. Each *λ* point was subjected to 40 ns of simulation time with the final 20-30 ns used for analysis, depending upon convergence. The free energy differences were estimated using the Bennett acceptance ratio method^69^ (BAR) implemented within GROMACS. The partitioning free energy ΔΔ*G*_*OW*_ is then the difference between Δ*G*_*W*_ and Δ*G*_*O*_, from which the partition coefficient (log *P*_*OW*_) may be calculated using:

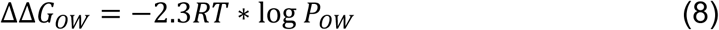

The *P*_*OW*_ values from simulations were compared to partition coefficient prediction methods such as ClogP and KOWWIN, which have been benchmarked against a wide variety of compounds^52^. The water-only simulations were composed of 1 disaccharide and 1000 water molecules, while the hydrated octanol simulations were composed of 1 disaccharide, 43 water molecules, and 519 octanol molecules, representing a 0.255 water/octanol molar fraction^70^. The vacuum-only simulations were composed of a single disaccharide in the simulation box.

### Umbrella sampling simulations to estimate binding affinities

Potential of mean force (PMF) profiles for the association of two solute molecules (glycan-glycan or lectin-glycan pairs) were calculated using umbrella sampling (US)^71^. Thus, multiple MD simulation windows were run along a pre-defined reaction coordinate – the separation distance between solute groups, along the *z*-axis of the simulation box – with an additional biasing harmonic potential. For a lectin-glycan pair, the groups were the center of mass of the glycan and the center of mass of residues defining the binding site in the protein. First, the two solutes were pulled away from each other at a rate of 10 nm ns^−1^, in order to generate the initial coordinates for the US windows. In the case of glycan-lectin PMFs, the glycan was pulled away from the lectin binding site. Both pulling simulations and US windows employed a harmonic potential between the centres of mass of the two groups of interest along the *z*-axis using a force constant of 1000 kJ mol^−1^ nm^−2^. Each complete pulling simulation corresponded to a 3 to 5 nm distance and resulted in 30 to 40 US windows with a 0.12 nm spacing. Additional windows were added in the regions of the minima to achieve greater overlap between windows, where necessary. Each window was subjected to production runs of 500 ns, leading to 40×500 ns (20 μs) of sampling per system per replica. For each system, at least two simulation replicas were performed. Block analysis was performed in order to assess the convergence. This was done by splitting the PMF trajectories into 100 ns windows. The PMFs were constructed using GROMACS’s inbuilt Weighted Histogram Analysis Method^72^ (WHAM) algorithm. The converged part of the trajectory was used for constructing the final PMF. 200 cycles of bootstrapping using the *b-hist* (Bayesian Bootstrapping) method with a tolerance of 10^−6^ was used to estimate the standard deviation across all replicas.

### Calculation of the binding free energy with standard state correction (ΔG^0^)

The binding free energy (Δ*G*^0^) for each glycan to its lectin was calculated from the one dimensional PMF *W*(*z*) by defining Δ*W* as zero at the minimum of the PMF curve minus the exponential average of the PMF over the unbound region, while a correction term (Δ*G*_*v*_) was added to account for the standard state volume *v*^0^ = 1661 Å^3^ based on the sampled unbound volume (*v*_*u*_)^73^:

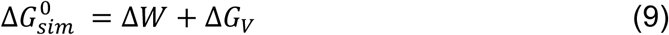

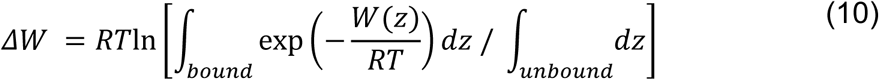

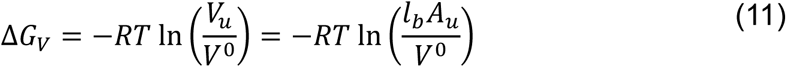

where *A*_*u*_ is the area sampled by the ligand in the unbound region. The protein backbone was restrained during the PMF calculation to prevent its rotation, which also prevented the ligand forming unproductive interactions with regions distant from the binding site in umbrella windows at increasing *z*-values. The unbound area was approximated as the cross-sectional area of the simulation box (i.e. the *xy* plane), following verification of complete sampling in *x* and *y* by each ligand across all unbound 500 ns umbrella windows (Figure S2). *l*_*b*_ is the bound length calculated by:

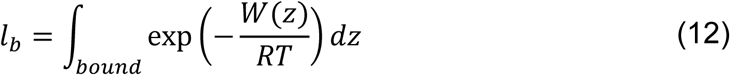

No further rotational entropy corrections were necessary, as the ligands were allowed full rotational sampling in the unbound region. The standard state free energies of binding were compared with the corresponding experimental values.

### Second virial coefficients (B_22_)

Osmotic data provides information about the nature of interactions between two solute molecules. This informs on how much the simulated association deviates from experimental measurements with molar concentration (*c*)^53^ and has previously been used to correct carbohydrate^74^ and protein^49,75^ FFs. In particular, the second virial coefficient (*B*_*22*_) comes from the virial expansion of pressure of many particle systems given by^57^:

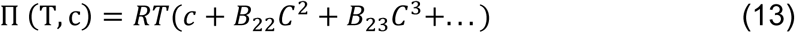

where *B*_*ij*_ are coefficients of virial expansion, *B*_*22*_ is the second virial coefficient, *T* is the temperature, and *R* is the gas constant. *B*_*22*_ > 0 indicates repulsive interactions between the two solutes while *B*_*22*_ < 0 indicates attractive interactions. The *B*_*22*_ value can be experimentally determined by self-interaction chromatography^76^ or diffraction studies^77^. McMillan and Mayer derived a method to calculate *B*_*22*_ values using MD simulations^53^ providing a powerful tool that can be used to optimise FFs. In this method, the PMF *W*(*z*) can be used to obtain the *B*_*22*_ value using the following expression:

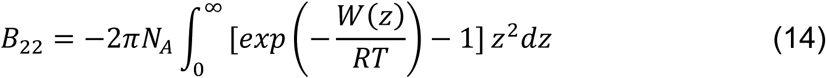

where *N*_*A*_ is the Avogadro number, *R* is the gas constant, *T* is the temperature of the system, and *z* is the distance between the solute molecules.

### N-glycans from di/trisaccharides

CG glycans such as high mannose, bi/tri/tetra-antennary and bisected complex glycans were constructed from component di/trisaccharide units (Figure 1). Parameters for the connecting angle between a sugar and its second neighbour were missing. To obtain these parameters, corresponding ATM glycans were constructed using the GLYCAM glycan builder^40^. Again, a similar methodology as used for di/trisaccharides was used to generate pseudo CG trajectories from 1 μs ATM simulations. As shown in Figure 2C-E, the glycosidic bonds between N-acetylglucosamine and mannose in a tetra-antennary complex glycan can be β(1,2/4/6). These branch patterns were parameterized separately. The angles obtained are summarized in Table 1. When constructing full length glycans these parameters are added to the disaccharide parameters depending upon the topology of the glycan constructed. The different types of glycans are named according to their Oxford notation^78^. This is based on building up N-glycan structures and it can be used to denote very complex glycans: all N-glycans have two core GlcNAcs; a given number of mannose sugars on the core GlcNAcs are denoted by Mx (e.g. M3, M5, M9); the number of antennae on the trimannosyl core are given by Ax (e.g. A2, A3, A4); Gx is the number of linked galactose units on antennae (eg. A2G1, A3G3); Sx is used to denote the number sialic acids linked to galactose (e.g. A2G2S2, A3G3S1); and an F start denotes the presence of fucose (e.g. FA2, FA3G2S1).

### Aggregation studies

It was previously reported^48,63^ that sugars in the Martini FF have a greater tendency to aggregate than observed experimentally. To investigate this further, systems containing 35 to 40 molecules of glycans were set up, with a ∼50 g L^−1^ concentration. Glycans should be readily soluble at concentrations of 50 g L^−1^, as dextran, a branched polymer of glucose is soluble even at a concentration of 400 g L^−1 79^. The simulations were performed using Martini v2.2, Martini 3.0b, GROMOS54a7 as well as CHARMM36m. To estimate the aggregation propensity of these glycans, RDFs were calculated. PMFs were also calculated to quantify the aggregation strength using Martini v2.2, GROMOS54a7 and CHARMM36m FFs. Triplicate simulations were used to construct the final PMFs at each scale factor. Scaling factors (λ) of 1.0, 0.9, 0.7, 0.5 and 0.3 were used to estimate second virial coefficients (B_22_) according to equation 14.

### Lectin binding studies

To supplement the partitioning and virial data, we pursued a complementary approach to validate non-bonded interactions by calculating binding free energies of different types of glycans with the lectins, which are selective for specific sugar patterns. Lectins are a class of proteins which selectively bind to mono or oligosaccharide molecules with specific glycosidic linkages^18^. This makes them good candidates for testing and validating the bonded as well as non-bonded parameters of our glycans. A total of seven candidate lectins, including cyanovirin-N (CVN), concanavalin-A (CONA), pterocarpus anolensis (PAL), ricinus communis agglutinin (RCA), wheat germ agglutinin (WGA), Maackia amurensis (MAA) and urtica dioicia agglutinin (UDA) were chosen; each has available crystal structures, along with either Isothermal Calorimetry (ITC) or Surface Plasmon Resonance (SPR) data for binding to various types of N-glycans. For every lectin, the ATM structure was converted to CG resolution using the *martinize*.*py* script from the Martini website. An elastic network with upper and lower cut-off values of 0.5 and 0.9 nm, respectively, and a force constant of 500 kJ mol^−1^ nm^− 2^ was implemented in addition to secondary structural bond/dihedral terms to maintain the higher order structures of all lectins. While setting up the lectin-glycan systems, the sugars which were resolved in the crystal structures were aligned with the CG glycan models, while non-interacting branches of the glycan were modelled pointing outwards, into solvent. The lectins used in this study, their PDB IDs and their binding affinities for various glycans obtained either from ITC or SPR experiments are summarized in Table 3. For reasons that will become apparent below, we calculated PMFs with scaling factors of 1, 0.95, or 0.9 for each of the 13 glycan-lectin pairs (Table 3, Figure 5).

**Table 2:** Thermodynamic properties of CG n-glycan di/trisaccharides. Free energies of solvation and partition coefficients for various disaccharides compared to predictions methods ClogP and KOWWIN. The errors estimated for the solvation free energies in water (Δ*G*_*W*_) and in octanol (Δ*G*_*O*_) for obtaining the partitioning free energies (ΔΔ*G*_*OW*_) were incorporated into the final reported partition coefficients (log *P*_*OW*_). These were compared against empirical predictions of log *P*_*OW*_ obtained from ClogP and KOWWIN^52^.

| Molecule | $\Delta G_W$<br>(kcal mol<sup>-1</sup>) | $\Delta G_O$<br>(kcal mol<sup>-1</sup>) | $\Delta\Delta G_{OW}$<br>(kcal mol<sup>-1</sup>) | $\log P_{OW}$ | $\log P_{OW}$<br>CLogP | $\log P_{OW}$<br>KOWWIN |
| --- | --- | --- | --- | --- | --- | --- |
| Fuc-α16-GlcNAc | -34.9 ± 0.2 | -29.1 ± 0.1 | 5.8 ± 0.3 | -4.1 ± 0.3 | -3.2 | -4.0 |
| GlcNAc-β14-GlcNAc | -36.2 ± 0.1 | -31.0 ± 0.2 | 5.2 ± 0.3 | -3.7 ± 0.2 | -4.1 | -4.1 |
| Man-β14-GlcNAc | -32.8 ± 0.2 | -27.2 ± 0.1 | 5.6 ± 0.3 | -4.0 ± 0.2 | -4.1 | -4.5 |
| Man-α16-[Man-α13-Man] | -41.7 ± 0.2 | -33.5 ± 0.2 | 8.2 ± 0.4 | -5.8 ± 0.3 | -5.9 | -6.5 |
| Man-α12-Man | -28.9 ± 0.1 | -23.5 ± 0.1 | 5.4 ± 0.2 | -3.8 ± 0.1 | -4.0 | -4.0 |
| Gal-β14-GlcNAc | -32.1 ± 0.1 | -26.8 ± 0.2 | 5.3 ± 0.3 | -3.7 ± 0.1 | -4.1 | -4.5 |
| Neu5Ac-α23-Gal | -35.4 ± 0.2 | -28.2 ± 0.4 | 7.2 ± 0.6 | -5.1 ± 0.4 | -5.3 | -6.0 |

**Table 3:**
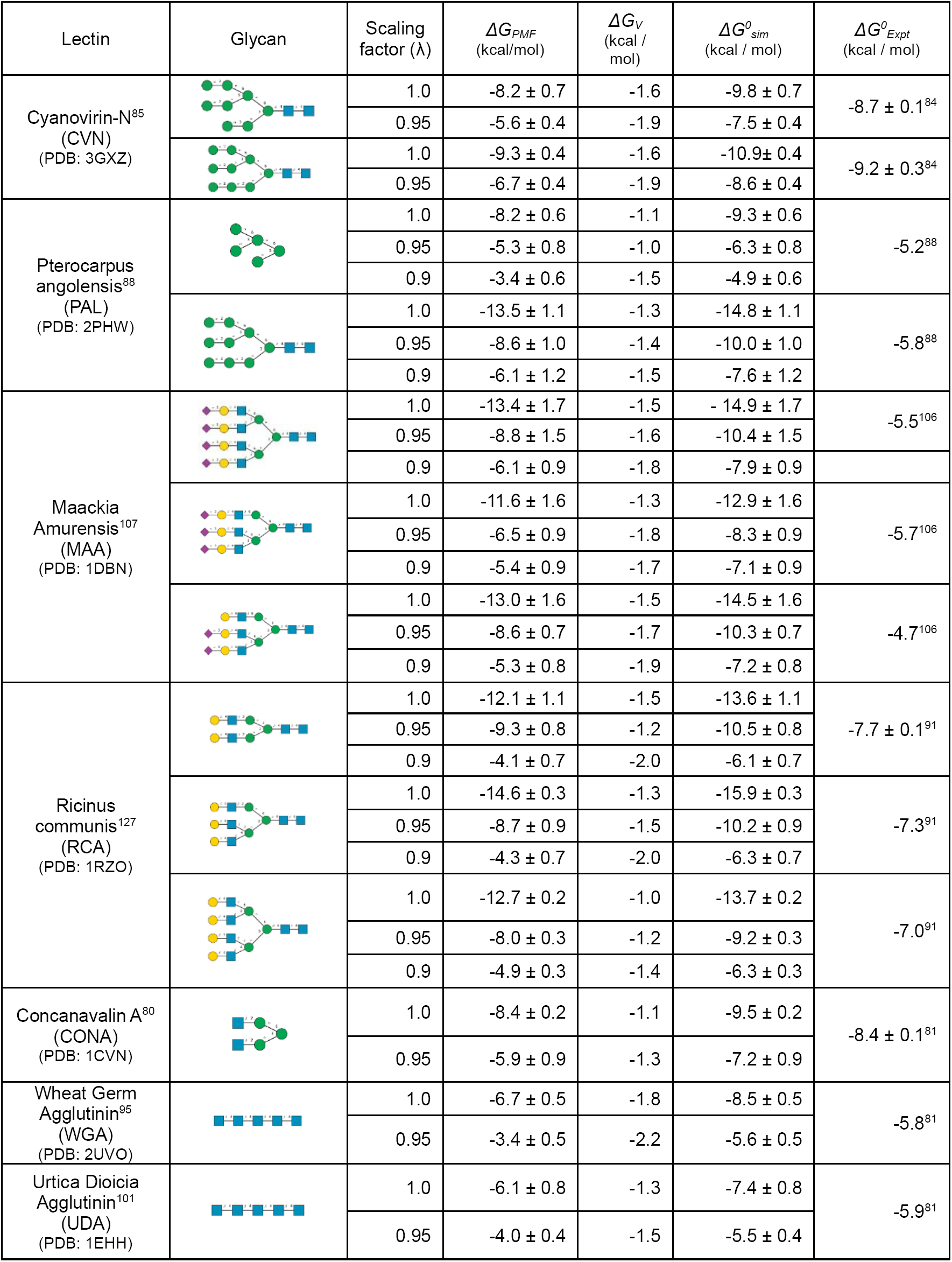
Lectins used for calculating the binding free energies (*ΔG*^*0*^_*sim*_) of various glycans to lectins. The binding affinities were obtained from either SPR or ITC experiments (*ΔG*^*0*^_*Expt*_). Errors calculated from PMFs (*ΔG*_*PMF*_) were obtained from 200 cycles of bootstrapping. The binding free energy Δ*G*^0^_*sim*_ was calculated upon addition to the *ΔG*_*PMF*_ of a correction term (Δ*G*_*v*_) to convert to standard state volume, for comparison with the experimental binding free energy Δ*G*^0^_*Expt*_. Monosaccharides present in the glycans are represented by their symbolic representation, including mannose (green circle), N-acetylglucosamine (blue square), galactose (yellow circle), and Neu5Ac/sialic acid (purple diamond).

| Lectin | Glycan | Scaling factor ($\lambda$) | $\Delta G_{PMF}$ (kcal/mol) | $\Delta G_V$ (kcal / mol) | $\Delta G^0_{sim}$ (kcal / mol) | $\Delta G^0_{Expt}$ (kcal / mol) |
| --- | --- | --- | --- | --- | --- | --- |
| Cyanovirin-N<sup>85</sup><br>(CVN)<br>(PDB: 3GXZ) |  | 1.0 | -8.2 ± 0.7 | -1.6 | -9.8 ± 0.7 | -8.7 ± 0.1<sup>84</sup> |
|  |  | 0.95 | -5.6 ± 0.4 | -1.9 | -7.5 ± 0.4 |  |
|  |  | 1.0 | -9.3 ± 0.4 | -1.6 | -10.9 ± 0.4 | -9.2 ± 0.3<sup>84</sup> |
|  |  | 0.95 | -6.7 ± 0.4 | -1.9 | -8.6 ± 0.4 |  |
| Pterocarpus angolensis<sup>88</sup><br>(PAL)<br>(PDB: 2PHW) |  | 1.0 | -8.2 ± 0.6 | -1.1 | -9.3 ± 0.6 | -5.2<sup>88</sup> |
|  |  | 0.95 | -5.3 ± 0.8 | -1.0 | -6.3 ± 0.8 |  |
|  |  | 0.9 | -3.4 ± 0.6 | -1.5 | -4.9 ± 0.6 |  |
|  |  | 1.0 | -13.5 ± 1.1 | -1.3 | -14.8 ± 1.1 | -5.8<sup>88</sup> |
|  |  | 0.95 | -8.6 ± 1.0 | -1.4 | -10.0 ± 1.0 |  |
|  |  | 0.9 | -6.1 ± 1.2 | -1.5 | -7.6 ± 1.2 |  |
| Maackia Amurensis<sup>107</sup><br>(MAA)<br>(PDB: 1DBN) |  | 1.0 | -13.4 ± 1.7 | -1.5 | -14.9 ± 1.7 | -5.5<sup>106</sup> |
|  |  | 0.95 | -8.8 ± 1.5 | -1.6 | -10.4 ± 1.5 |  |
|  |  | 0.9 | -6.1 ± 0.9 | -1.8 | -7.9 ± 0.9 |  |
|  |  | 1.0 | -11.6 ± 1.6 | -1.3 | -12.9 ± 1.6 | -5.7<sup>106</sup> |
|  |  | 0.95 | -6.5 ± 0.9 | -1.8 | -8.3 ± 0.9 |  |
|  |  | 0.9 | -5.4 ± 0.9 | -1.7 | -7.1 ± 0.9 |  |
|  |  | 1.0 | -13.0 ± 1.6 | -1.5 | -14.5 ± 1.6 | -4.7<sup>106</sup> |
|  |  | 0.95 | -8.6 ± 0.7 | -1.7 | -10.3 ± 0.7 |  |
|  |  | 0.9 | -5.3 ± 0.8 | -1.9 | -7.2 ± 0.8 |  |
| Ricinus communis<sup>127</sup><br>(RCA)<br>(PDB: 1RZO) |  | 1.0 | -12.1 ± 1.1 | -1.5 | -13.6 ± 1.1 | -7.7 ± 0.1<sup>91</sup> |
|  |  | 0.95 | -9.3 ± 0.8 | -1.2 | -10.5 ± 0.8 |  |
|  |  | 0.9 | -4.1 ± 0.7 | -2.0 | -6.1 ± 0.7 |  |
|  |  | 1.0 | -14.6 ± 0.3 | -1.3 | -15.9 ± 0.3 | -7.3<sup>91</sup> |
|  |  | 0.95 | -8.7 ± 0.9 | -1.5 | -10.2 ± 0.9 |  |
|  |  | 0.9 | -4.3 ± 0.7 | -2.0 | -6.3 ± 0.7 |  |
|  |  | 1.0 | -12.7 ± 0.2 | -1.0 | -13.7 ± 0.2 | -7.0<sup>91</sup> |
|  |  | 0.95 | -8.0 ± 0.3 | -1.2 | -9.2 ± 0.3 |  |
|  |  | 0.9 | -4.9 ± 0.3 | -1.4 | -6.3 ± 0.3 |  |
| Concanavalin A<sup>80</sup><br>(CONA)<br>(PDB: 1CVN) |  | 1.0 | -8.4 ± 0.2 | -1.1 | -9.5 ± 0.2 | -8.4 ± 0.1<sup>81</sup> |
|  |  | 0.95 | -5.9 ± 0.9 | -1.3 | -7.2 ± 0.9 |  |
| Wheat Germ Agglutinin<sup>95</sup><br>(WGA)<br>(PDB: 2UVO) |  | 1.0 | -6.7 ± 0.5 | -1.8 | -8.5 ± 0.5 | -5.8<sup>81</sup> |
|  |  | 0.95 | -3.4 ± 0.5 | -2.2 | -5.6 ± 0.5 |  |
| Urtica Dioica Agglutinin<sup>101</sup><br>(UDA)<br>(PDB: 1EHH) |  | 1.0 | -6.1 ± 0.8 | -1.3 | -7.4 ± 0.8 | -5.9<sup>81</sup> |
|  |  | 0.95 | -4.0 ± 0.4 | -1.5 | -5.5 ± 0.4 |  |

**Figure 3:**
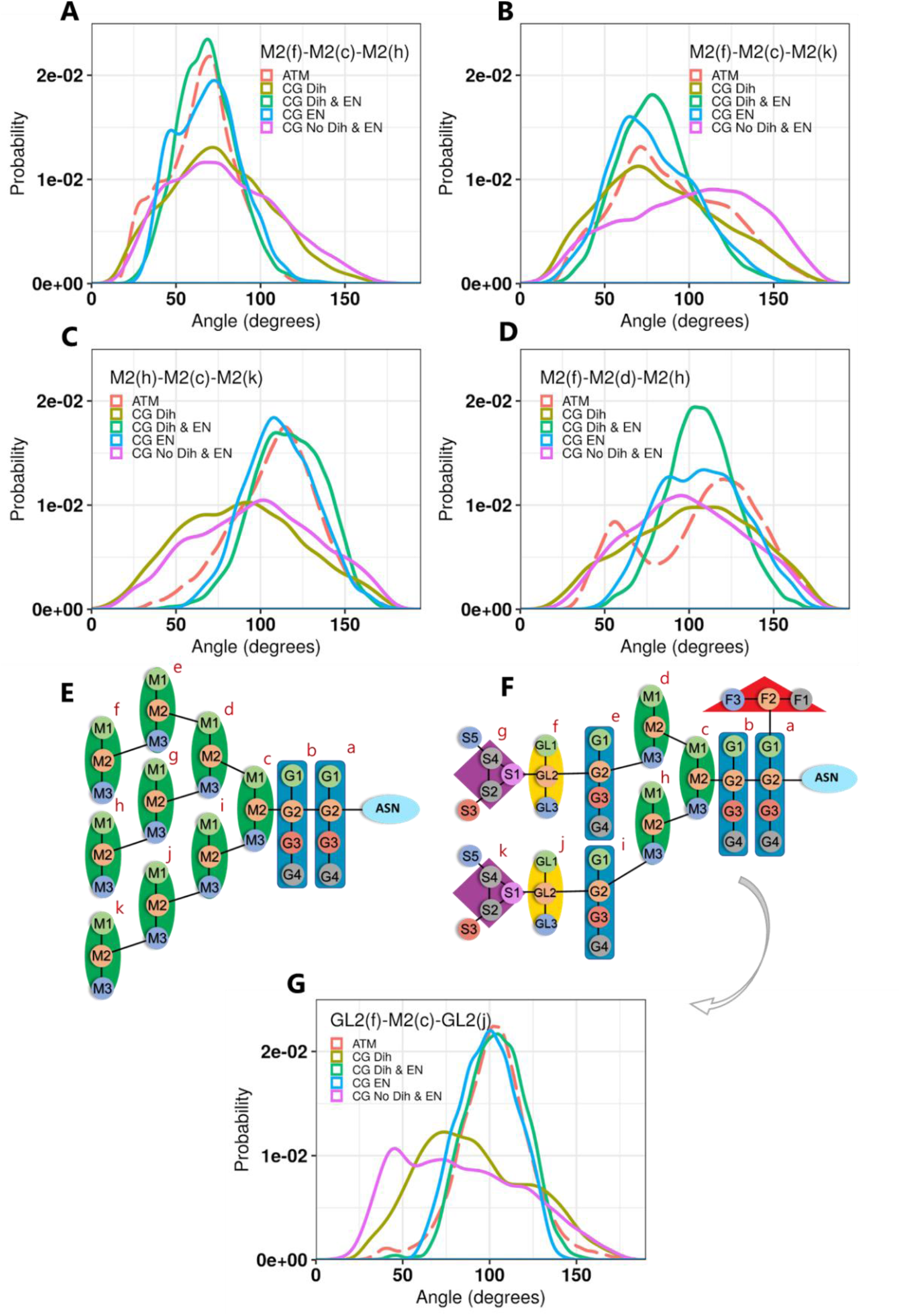
Branch angle distributions for M9 and FA2G2S2 type glycans. Distributions compare data from ATM simulations versus those for CG with dihedrals (CG Dih), CG with dihedrals and elastic network (CG Dih + EN), CG with elastic network only (CG EN), or CG with neither dihedrals or elastic network (CG No Dih & EN). The angles in plots A, B, C and D are for M9 as illustrated in (E). The angle in plot G is for FA2G2S2 complex type glycan as illustrated in (F).

**Figure 4:**
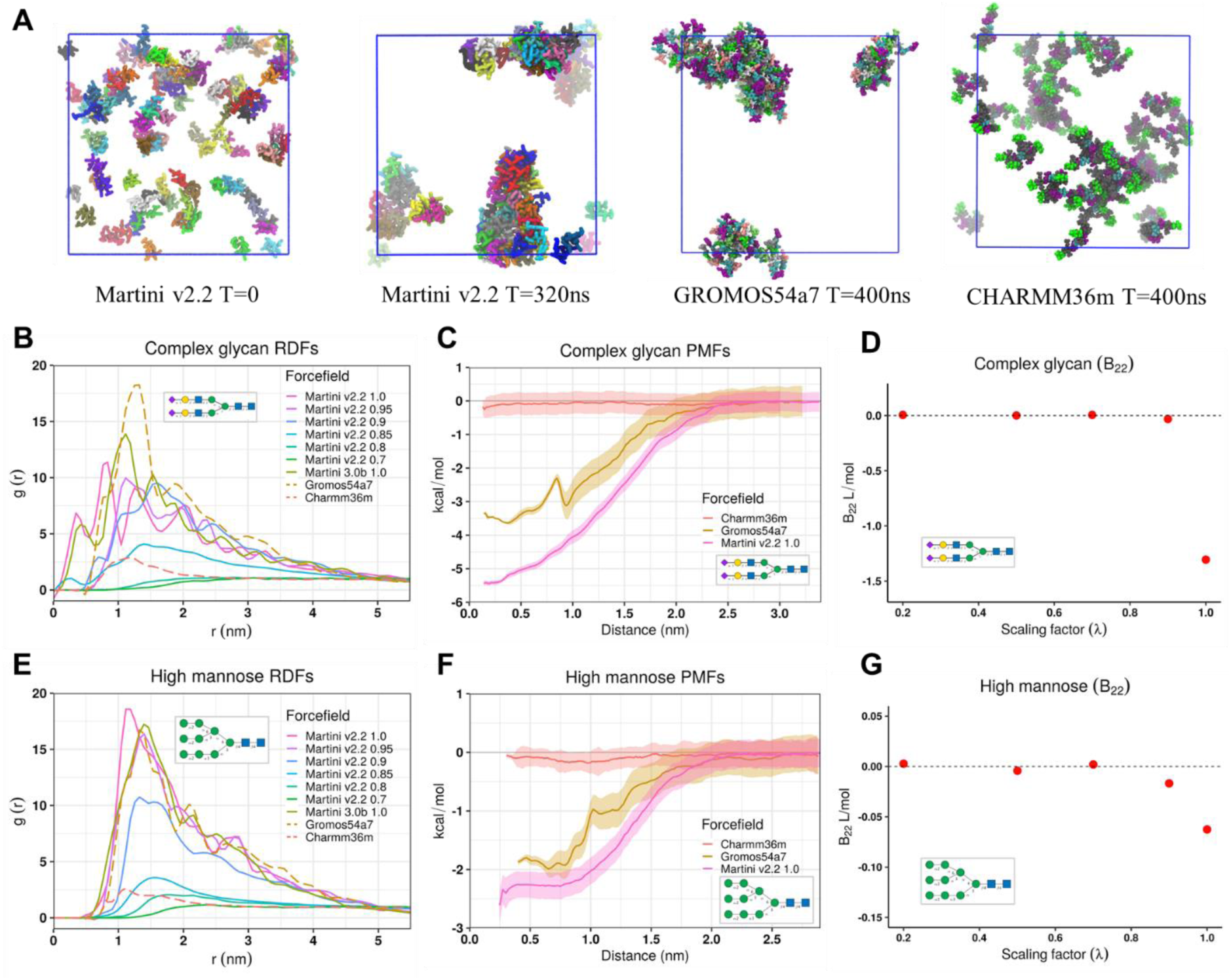
Aggregation propensity for complex and high mannose glycans using a range of FFs. (A) Initial and aggregated stages of glycan simulations for Martini v2.2, GROMOS54a7 and CHARMM36m forcefield. (B), (E) Radial distribution functions (RDFs) of glycans with various FFs. (C), (F) Potential mean of forces (PMFs) for glycans with various FFs. (D), (G) Partial virial coefficients (B_22_) for Martini v2.2. Monosaccharides present in the glycans are represented by their symbolic representation: mannose (green circle), N-acetylglucosamine (blue square), galactose (yellow circle), and Neu5Ac/sialic acid (purple diamond).

**Figure 5:**
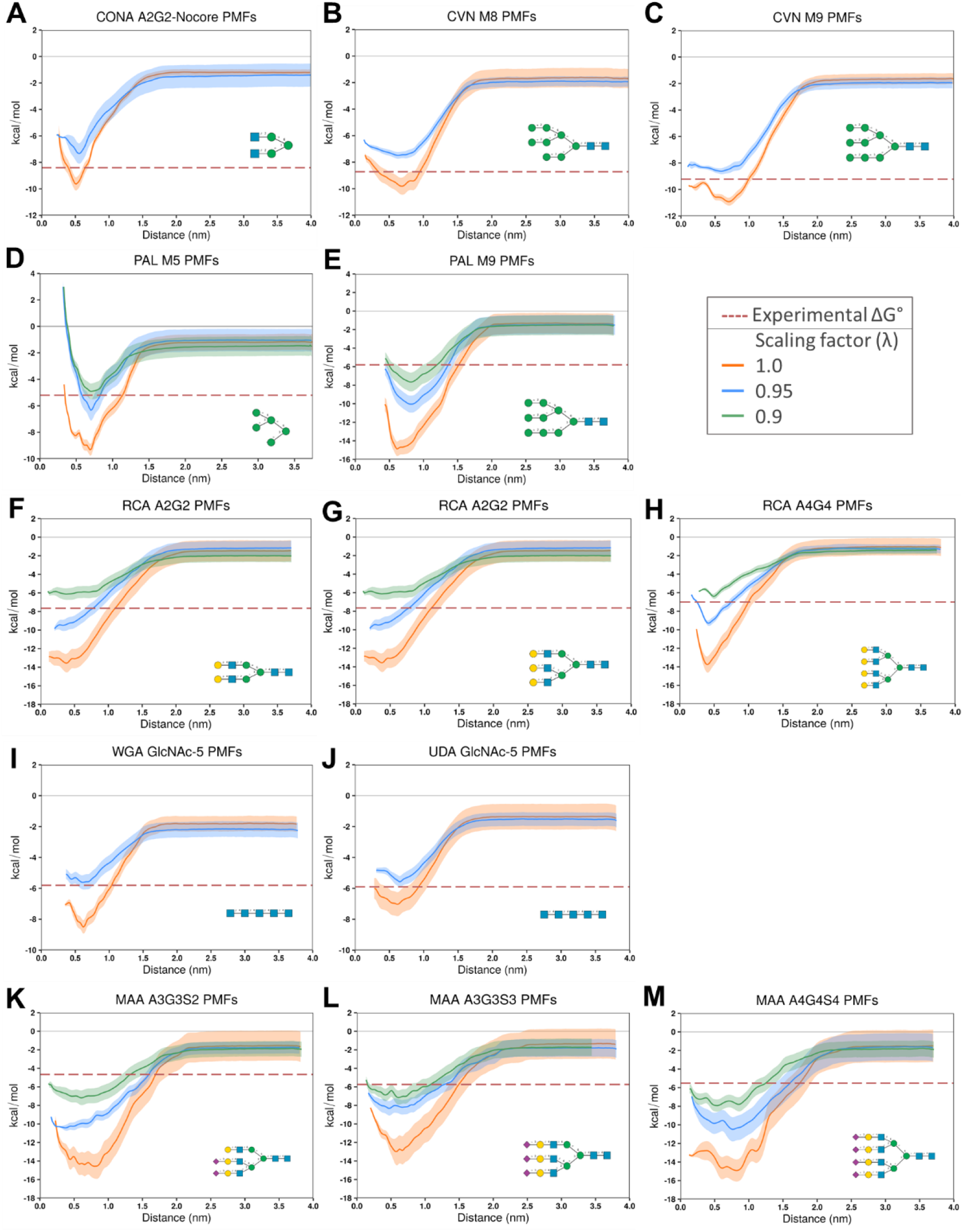
PMFs for different lectin-glycan systems. The glycan used for binding studies in each case is given in the right bottom corner of each plot. Error estimates are shown with the shaded region obtained from 200 cycles of bootstrapping. (A) Concanavalin A (CONA); B), (C) Cyanovirin-N (CVN); (D), (E) Pterocarpus Anolensis (PAL); (F), (G), (H) Ricinus Communis Agglutinin (RCA); (I) Wheat Germ Agglutinin (WGA); (J) Urtica Dioicia Agglutinin (UDA); and (K), (L), (M) Maackia Amurensis (MAA). Monosaccharides present in the glycans are represented by their symbolic representation: mannose (green circle), N-acetylglucosamine (blue square), galactose (yellow circle), and Neu5Ac/sialic acid (purple diamond). The volume corrections *ΔG*_*v*_ were added to the total PMF in order to visually compare with the experimental data (dashed lines).

## Results

### Elastic network in extended N-glycans

As previously reported for studies of glycolipids and some oligosaccharides^48,63^, when applying dihedral potentials in simulations of our extended CG glycans, numerical instabilities limited the maximum time step to 5 fs. This was alleviated by using ReB angle potentials, enabling a timestep of 10 fs. To retain the overall conformation in accordance with the ATM models, elastic connections were also implemented between the central bead of the first mannose of the trimannosyl (M3) core and the last monosaccharide of each branch of the glycans shown in Figure 2. In addition, harmonic angle potentials (Table 1) were added between the branch ends and the M3 core. Looking at the branch angle distributions of high mannose (M9) and complex type (FA2G2S2) glycans (Figure 3), it was observed that dihedrals alone could not reproduce the AA distributions. The distributions in CG simulations with only dihedrals were wider compared to their ATM counterparts. The distributions with either dihedrals plus elastic network or elastic network alone could, however, reproduce the ATM data closely, confirming the requirement of an elastic network to maintain the overall conformations of the glycans. This allowed us to run our simulations of glycans having an elastic network with dihedrals switched off using a stable time step of 15 fs. All the elastic network parameters are summarised in Table 1.

### Partitioning behaviour

The Martini forcefield building block parameters were derived in part according to their partitioning behaviour. In previous carbohydrate development efforts, experimental partitioning energies were accurately reproduced for various small sugars including glucose, fructose, sucrose, maltose, cellobiose, kojibiose, sophorose, nigerose, laminaraibose and trehalose^46^. Although the bead type selection is very similar to these sugars, new parameters were derived for variants including GlcNAc, Neu5Ac and Fuc. Thus, we calculated solvation free energies of various n-glycan disaccharides in water (Δ*G*_*W*_) and octanol (Δ*G*_*o*_) phases to obtain partition coefficients (log *P*_*OW*_) for comparison with corresponding values from empirical fragment-based models (Table 2). The log *P*_*OW*_ values were negative for all sugars, consistent with their preference towards the water phase. Overall, the simulated values are in reasonable agreement with those obtained from the empirical models, with slightly closer accordance with the ClogP data compared to KOWWIN, consistent with previous studies^52^.

### Aggregation of glycans

Simulations of both high mannose and complex glycans at ∼50 g L^−1^ concentration, a concentration at which all of them should be soluble, led to aggregation within a few hundred nanoseconds at both CG and AA resolution, except when using the CHARMM36m FF (Figure 4). These aggregates remained irreversibly associated even after extending the simulations to 2 µs. When comparing against ATM FFs, GROMOS54a7 showed very similar behaviour to Martini v2.2 and 3.0b in terms of both RDFs and PMFs. However, CHARMM36m did not form clusters in any of the simulations, and also resulted in a very shallow PMF well depth (Figure 4B-C, E-F). Martini 3.0b is still in the early stages of its development and the new mapping scheme is not yet available for simple carbohydrates. So, with the added bead types and interaction levels, the final sugar mapping may still improve the results in future. Similar to findings of a previous study^48^, non-bonded interactions in Martini v2.2 were found to be too attractive for the glycans. This is as reported for other Martini-parameterized molecules like glycolipids and proteins such as lysozyme^48,49,63^. Thus, we attempted to optimize the non-bonded parameters of the glycans, firstly by tuning the second virial coefficient (B_22_)^53^ and secondly by comparing the glycan binding free energies calculated from simulations with either ITC or SPR experiments. Scaling down the interactions alleviated the aggregation propensity, but required a scaling factor of 0.7 or below, to reach B_22_ coefficient values of ≥ 0 L mol^−1^, in both cases (Figure 4D,G). The experimental value of the B_22_ for the complex glycan (A2G2S2) is around 40 L mol^−1 48^, which could not be achieved even after reducing the scaling factor to as low as 0.3. Nevertheless, aggregates were not formed during 1 µs simulations of high mannose and complex glycans when a scaling factor of 0.85 was used (Figure 4B,E). PMFs calculated with these scaling factors implemented for both high mannose and complex glycan were flat for a scaling factor 0.9 or below, indicating dominating water-glycan interactions as observed in the experimental conditions (Figure S5). Importantly, we also observed that aggregation was dependent upon the size of the glycans (Figure S6). Monosaccharides did not require any scaling while a scaling of 0.95-0.9 was required for disaccharides. For sugars bigger than disaccharides, a maximum scaling factor of 0.85 was required to reproduce ATM RDFs including high mannose and complex glycans.

### Binding of glycans with different lectins

Based on the results of PMF calculations, described in detail below, every lectin-glycan pair overpredicted binding free energies from unscaled simulations. Scaling by 0.95 was sufficient for most of the pairs, except for the MAA and PAL lectins, where 0.9 scaling was required. The calculated binding free energies across all systems are summarised in the Table 3, with detailed information about the binding pocket in given in Table S3.

#### 1. Mannose binding lectins

##### 1.1 Concanavalin A (CONA)

The mannose binding specificity of CONA is dependent on the trimannoside core found in most N-glycans. The crystal structure^80^ (Figure S7A) shows tetrameric CONA interacting with four core trimannoside Man-α(1,6)-[Man-α(1,3)-]Man or M3. The interaction site includes residues Y12, N14, T15, D16, G98, L99, Y100, A207, D208, G227 and R228. ITC experiments with GlcNAc-β(1,2)-Man-α(1,6)[GlcNAc-β(1,2)-Man-α(1,3)]Man-OH glycan, which will be referenced as A2-nocore hereafter, yielded a ΔG of −8.4 ± 0.1 kcal mol^−1 81^. In our studies, the M3 sugars of the glycan were aligned with the crystal structure. Our simulations predicted a binding free energy of −9.5 ± 0.2 kcal mol^−1^ (Figure 5A) for λ=1.0, which is an overprediction of ∼13%.

##### 1.2 Cyanovirin-N (CVN)

CVN is a widely studied 110 kDa lectin because of its role in inactivation of many strains of Human Immunodeficiency virus (HIV)^82^. CVN preferentially binds to high-mannose oligosaccharides^83^. ITC experiments show that two mannose oligosaccharides M8 and M9 bind to CVN with binding affinities of −8.7 ± 0.1 and −9.2 ± 0.3 kcal mol^−1^ respectively^84^. The crystal structure for CVN with M9 (Figure S7B) reveals a binding interface of three stacked α(1,2) linked mannose sugars interacting with residues L1, G2, K3, T7, E23, N93, D95 and E101 of CVN while the rest of the chain is exposed to solvent^85^. In our studies, both M8 and M9 glycans were used to estimate the binding free energy. M8 and M9 have a terminal α(1,2) linked mannose which is important for the interaction. Similar behaviour to CONA was observed in the case of the CVN lectin. The calculated PMFs predict binding free energies of −9.8 ± 0.7 and −10.9 ± 0.4 kcal mol^−1^ (Figure 5B & 5C) for M8 and M9 respectively, which are overpredicted by ∼12% and ∼18% respectively.

##### 1.3 Pterocarpus Anolensis lectin (PAL)

PAL is a Mannose/Glucose specific lectin and multiple crystal structures showing its interactions with mono, di and trisaccharides are available^86,87^. Recent studies revealed PAL’s interactions with M9 and M5 high mannose glycans^88^ (Figure S7C). It was observed crystallographically that PAL interacts with M5 and M9 in the same unique way in which the Man-α(1,2)-Man-α(1,6)-[Man-α(1,3)-]Man-α(1,-) motif binds to PAL via a combination of van der Waal’s contacts and hydrogen bonds^88^. The glycan interacts with residues D36, N83, G106, D136, N138, E221. Both M5 and M9 interacts strongly with PAL, with binding affinities of −5.2 and −5.8 kcal mol^−1^ respectively, as shown by ITC^88^. Our simulations led to an overestimation of the interactions in the non-scaled systems for both glycans (Figure 5D & 5E). In the case of M5 and M9 the binding free energy was overestimated by ∼77% and ∼155% respectively. To achieve reasonable agreement with the experimental binding affinities, the interactions needed to be scaled by 0.9.

#### 2. Galactose/N-acetylgalactosamine binding lectins

##### 2.1 Ricinus Communis Agglutinin (RCA_**120**_)

RCA_120_ is a hemagglutinin and is tetrameric in nature. It has two α and two β subunits that are 29.5 and 37 kD in size, respectively. Out of the two types of subunits, it has been shown that the oligosaccharides interact only with the β subunits of the lectin^89^. RCA_120_ specifically recognises Gal-β(1,4) with very similar affinities for Gal-β(1,4)-GlcNAc, Gal-β(1,4)-Glc and Gal-β(1,4)-Man terminal residues^90^. SPR studies were performed with bi (A2G2), tri (A3G3) and tetra-antennary (A4G4) complex glycans (Table 3) and resulted in binding affinities of −7.7, −7.3, and −7.0 kcal mol^−1^, respectively^91^. Although there are no crystal structures showing direct interactions with any of the A2G2, A3G3 or A4G4 glycans, there is a crystallographic study showing interactions with two terminal galactoses (PDB 1RZO). In the structure (Figure S7D) of RCA_120_, the first galactose interacts with D22, G25, E26, Q35, K40 and N46, while the other galactose interacts with N95 and Y125. Considering the higher number of interactions of the first GlcNAc with polar and charged residues, it was used for the PMF calculations. For all branched glycans, including bi, tri, and tetra-antennary glycans, we found that the FF overestimates the binding free energy without any scaling of interactions (Figure 5F, 5G & 5H). The energies obtained from the simulation with a scaling factor of 0.9 were −6.1 ± 0.7, −6.3 ± 0.7 and −6.3 ± 0.3 kcal mol^−1^ with 20%, 14% and 10% deviation from the experimental values for A2G2, A3G3 and A4G4 glycans, respectively.

#### 3. Sialic acid/N-acetylglucosamine binding lectins

##### 3.1 Wheat Germ agglutinin (WGA)

WGA is an Neu5Ac and GlcNAc specific lectin which is antifungal in nature and has three isoforms^92,93^. The crystal structure reveals a stable dimer with each polypeptide showing four hevein domains responsible for GlcNAc recognition^94^, though not all eight binding sites were observed to be occupied in a single crystal^95^. In this structure (Figure S7G), the first GlcNAc occupied the region defined by residues D86, S105, F109 from monomer 1 and A71, E72 of monomer 2, while the region defined by residues S62, Y64, Y66, E72, Y73 of monomer 1 and S114, E115 of monomer 2 were occupied by the second GlcNAc. Binding studies revealed that WGA has the highest affinity (5.8 kcal mol^−1^) for (GlcNAc)_5_ and decreases to −3.7 kcal mol^−1^ as the number of sugars reduces from (GlcNAc)_5_ to (GlcNAc)_1_^81^. ITC experiments of single domain recombinant WGA with (GlcNAc)_3_ and (GlcNAc)_4_ resulted in a 10-fold lower binding constant than the wild type oligomer, emphasizing the importance of the dimer interface in the binding of oligosaccharides^96^. Considering the selectivity of (GlcNAc)_5_, it was used for the calculation of PMFs, based on the first dimeric interface from the crystal structure (Figure S7G). In our studies, the binding free energies for (GlcNAc)_5_ were overpredicted and scaling 0.95 was required to obtain a binding free energy of - 5.6 kcal mol^−1^ (Figure 5I) which represents a 4% deviation from experiment.

### 3.2 Urtica Dioicia agglutinin (UDA)

UDA is a chitin and is a GlcNAc oligomer specific lectin derived from plants^97,98^. UDA is speculated to be antifungal and insecticidal in nature^99,100^. Binding experiments suggest that the lectin has two binding sites with a preference for (GlcNAc)_5_, with an affinity of −5.9 kcal mol^−1^ that decreases to −3.9 kcal mol^−1^ upon a reduction in chain length to (GlcNAc)_2_.^81^ The crystal structure of UDA isolectin VI is available, revealing its interactions with (GlcNAc)_3_^101^ (Figure S7F). The GlcNAc oligomer interacts with UDA at residues S19, C24 and Y30. In our simulations, the binding free energy with the (GlcNAc)_5_ oligosaccharide – modelled based on the (GlcNAc)_3_ coordinates for the middle three GlcNAc groups – was overpredicted by ∼25% in the absence of scaling. A scaling of 0.95 yielded close agreement with the experimental value of −5.9 kcal mol^− 1^ (Figure 5J) with only a 7 % deviation.

### 3.3 Maackia Amurensis (MAA)

Maackia Amurensis has two isolectins, hemagglutinin and leukoagglutinin (MAA), which were identified by their agglutination activity against different blood cell types and their binding properties with either O-linked or N-linked oligosaccharides^102,103^. Subsequently, it was shown that MAA is specific towards Neu5Ac units, especially towards the NeuAc-α(2,3)-Gal-β(1,4)-GlcNAc/Glc oligosaccharide^104,105^. SPR binding studies of MAA with different N-glycans such as sialylated tri-antennary (A3G3S2), fully sialylated tri-antennary (A3G3S3) and fully sialylated tetra-antennary (A4G4S4) yielded similar affinities of −4.7, −5.7 and −5.5 kcal mol^−1^ respectively, suggesting a slight preference for the NeuAc-α(2,3)-Gal-β(1,4)-GlcNAc/Glcβ motif^106^. The crystal structure of MAA is dimeric with each monomer folding into large β-pleated sheets^107^. Each monomer shows interactions with sialyllactose at residues T45, D87, S104, L107, T131, T136 and T221 (Figure S7E). The sialyllactose coordinates were used as the initial coordinates for modelling all three glycans (A3G3S2, A3G3S3 and A4G4S4) with other branches pointing towards the solvent. PMF calculations revealed that sialic acid containing glycans result in overprediction of binding free energies by ∼125-200 % (Figure 5K, 5L & 5M). Scaling the interactions by 0.9 was required to obtain reasonable agreement with the experimental values.

## Discussion

The highly flexible nature of glycans limits detailed structure-dynamics-function studies using experimental techniques, but this gap may be supplemented by MD simulations. The behaviour of molecules in these simulations depends upon the FF used. In this work, we have extended the Martini parameters towards N-glycans. A slightly different mapping scheme was used where bonds were made between beads originating from ATM models compared to the one used by Lopez et al^46^ in which only the central beads were connected to describe the glycosidic linkage. Parameterization of disaccharides using this scheme was convenient for making highly branched patterns of N-glycan like bisected tetra-antennary complex glycans. Although all the N-glycan glycosidic linkages were parameterized using this mapping scheme, one should note that coarse-graining still leads to reduced accuracy. Firstly, it results in a loss of the explicit stereochemical nature and the hydrogen bonding network of sugars, which is important for the specific interactions with carbohydrate binding proteins (CBPs)^88,92,101,107^. The water network around the sugars is lost, and in turn can affect the local translational and rotational dynamics^108^. Although coarse-graining leads to loss of information, the Martini CG approach^45,109–112^ nevertheless implicitly maintains chemical identities by using appropriate polar bead types that have been experimentally validated against water-octanol partitioning free energies^46^.

Monosaccharides undergo chair-to-boat and chair-to-chair transitions, also referred to as “ring puckering”, making the choice of FF for calibration important. The ATM FF used in this study, GROMOS54a7^113^, reproduces these conformations very well. But in the Martini CG model, this effect is neglected. Although the ring can transition between ^4^C_1_ and ^1^C_4_ conformations in the ATM model, the overall shape of the sugar remains the same in the CG model as a result of the grouping of atoms into unified particles, making the sugar effectively linear (Figure 1). Therefore, the CG model should not be affected by the preference of the ATM FF towards a specific conformation showing the average structure of these puckering transitions. Also, the more common dextrorotatory (D) form of sugars was considered for parametrization^114–116^.

The CG approach also affects the degree to which one can distinguish between α and β anomers, which were not considered here as the anomeric form of most of the sugars is already fixed for N-glycans. Multiple rotameric states of the hydroxymethyl group and its preference towards gg(−60°) and gt(60°)^117,118^ conformations was observed in the ATM simulations. We modelled the bimodal distributions using a single harmonic potential by fitting them to the most populated conformation observed. It should be noted that in future, they could be modelled via tabulated potentials^119^. The glycosidic linkages in di/tri-saccharides were represented by using dihedral potentials which orient the monosaccharides relative to each other. But in the case of N-glycans, dihedrals resulted in numerical instabilities due to geometric tension between the glycosidic bonds, as observed in other studies^48,63^. The problem could either be solved by using a smaller timestep, which cripples the efficiency of the CG approach, or by not using the dihedral potentials at all. The first approach has been used for glycolipids^63^ while the latter has been used for some oligosaccharides^48^. A timestep of 5 fs or less is manageable but seriously limits the benefits of the CG approach, becoming computationally inefficient for larger biologically relevant systems such as glycoprotein-antibody complexes. This was partially alleviated here by using ReB angle potentials, which improves the associated numerical instabilities. Furthermore, an elastic network proved beneficial in maintaining the overall conformations of the glycans (Figure 3). This ultimately allowed the simulations to be run using a timestep of 15 fs. An alternative approach to the elastic network that could be investigated in future would be to introduce virtual sites, within the trimannose core, the disaccharides, or at other branch sites, in order to maintain key conformations via effective dihedrals^120^.

A first step towards validating the non-bonded interactions in new Martini parameters involves calculating partition coefficients for the building blocks of the molecules of interest. We observed that the selection of bead types for our di/saccharide combinations correlated well with the partition data from empirical prediction methods^46^. However, constructing full length glycans from these building blocks showed some serious issues in terms of self-aggregation. Comparing the effect of the elastic network on the self-interaction energies of M9 and A2G2 complex type glycans suggests that the elastic network does result in slightly higher binding free energies (Figure S4). It has recently been shown that this is likely because of weak force constants in the elastic network resulting in a short bond length effect which creates “superinteraction” centres^121^. This observed effect is small in our model as we have a maximum of just three elastic bonds, compared to 24 in the polyleucine model used in that study. But even without using an elastic network, the predicted self-binding energies for the glycans were observed to be negative, which resulted in the well-documented “sticky” behaviour. Similar observations were made in a related study on few oligosaccharides^48^. Thus, Schmalhorst et al^48^ showed that carbohydrates including glucose (monosaccharide), sucrose (disaccharide), α/β-cyclodextrin (cyclic), and sialylated biantennary glycan (A2G2S2) spuriously aggregate within a few hundred nanoseconds and proposed a 50% scaling down of non-bonded interactions for oligosaccharides. Likewise, for our glycan models including A2G2S2 and high mannose (M9) (Figure 4), carbohydrate molecules aggregated within a few hundred nanoseconds. The calculated B_22_ values were negative (Figure 4) consistent with interactions between sugars being attractive in the Martini representation, compared to positive and hence repulsive experimental B_22_ values^48^. This sticky behaviour is suggested to be in part the result of mixing smaller sized beads with regular (R) beads creating artificial energy barriers^121^. The effect is significant with the “tiny” (T type) beads while much lower with “small” beads (S type). So, depending upon the type and length of the glycan, the added artificial barriers will potentially aggravate the sticky nature of Martini FF.

The overly attractive behaviour observed for oligosaccharides in the Martini FF calls for adjustments in the underlying non-bonded interactions. A simple way of solving this issue is to make the water-glycan interactions stronger or make glycan-glycan interactions weaker. The latter approach is generally preferred^48,49^ so as to keep central properties of the Martini FF constant such as the partitioning behaviour between water and apolar solvents, hence avoiding the need for complete reparameterization of the entire FF. Thus, the solvent-solvent and solute-solvent interactions were not changed. Weakening the glycan-glycan interactions partly solved the issues (Figure 4). Scaling by as much as λ=0.7 was required to reach B_22_ coefficients of 0 L mol^−1^ for the high mannose and complex type glycans. The experimental B_22_ value for A2G2S2 glycan was not attainable by scaling down the interactions drastically. Visual inspection and examination of RDFs suggested that aggregation was reversible when a scaling factor of 0.85 or lower was used, suggesting that B_22_ is a somewhat problematic choice of parameter for optimization of non-bonded interactions (Figure 4). It is also noteworthy that the scaling factor required for reversing the aggregation is dependent upon the size of the glycan studied (Figure S6).

While solution properties of the glycans are important, their interactions with other biomolecules are equally critical. The binding of N-glycans to lectins, carbohydrate binding proteins that make specific interactions with terminal sugars of glycan chains^92,97,99,101–105,122^, are crucial in many biological phenomena. Thus, optimizing the binding properties of these glycans with proteins is important for their applicability in multi-protein complexes, and reproduction of experimental affinities of glycans with proteins represents a useful way of optimizing the non-bonded parameters, as shown here. Binding free energies obtained for a total of thirteen candidate lectin-glycan pairs showed that our glycan models can reproduce the experimental binding affinities without the need for drastic corrections in non-bonded interactions (Figure 5).

It should be noted that for many of the systems, the exact binding mode of the whole glycan was often unresolved. The partially resolved sugars in available crystal structures were thus used for aligning and constructing the whole glycan which can result in multiple initial conformations. Replicates ensured that the initial structure bias was reduced. Cluster analysis showed that these glycans could maintain the overall binding pose during umbrella sampling simulations (Figure S10). A simple glycan such as (GlcNAc)_5_ could distinguish between a favourable as well random surfaces on a UDA lectin (Figure S11). Unrestrained simulations of high affinity CVN+M9 maintained a very similar binding pose when compared to ATM simulations, whereas in the case of the low affinity PAL and M5 pair, the glycan was more dynamic and could drift from the pocket in both ATM and CG representations, consistent with the weak binding of the ligand (Figure S12, S13). All these observations support the applicability of the newly derived CG N-glycan models for specifically quantifying energetics with given protein-ligand pairs.

Out of 13 different lectin-glycan systems, every one of them overpredicted the binding free energies calculated during unscaled simulations. A 0.95 scaling was enough to reduce the gap between the predicted and experimental binding free energies by a large factor. It was observed that charged complex glycans – which contain an explicit charge as well as a higher number of S type beads – required a higher scaling of 0.9. In the PAL lectin, which also required 0.9 scaling, the glycan interacts with the lectin via a relatively high number of polar residues in the binding pocket compared to any other lectins (Table S3). Thus, the scaling required appears to be in part glycan type dependent, particularly when electrostatics and a greater number of small type beads are involved.

The scaling approach for Martini was first proposed by Stark et al^49^ due to the imbalance of the non-bonded interactions in the Martini FF for protein-protein systems, and has since been used for glycan-glycan^48^ interactions as well. In studies of dimerization of different receptors such as ErbB1 and EphA1, binding free energies were again overestimated^123–126^. Javanainen et al^50^ predicted the binding free energies of dimerization of TM domains of five candidate receptor tyrosine kinases (RTKs) and suggested a relatively modest correction of 10% in the well depth (ε) to achieve better agreement with FRET studies^50^ compared to the 60% correction suggested by Stark et al^49^ where they compared their PMFs against the B_22_ coefficient. In the present work, we also obtained data pointing towards the imbalance in the Martini FF, but this was not drastic and was alleviated by scaling the non-bonded interactions by a relatively small value.

## Conclusions

In summary, we have extended the Martini CG model parameters to N-glycans with various branching patterns. An elastic network was found to be advantageous in maintaining the conformations of branched glycans. The spurious self-aggregation of glycans could be alleviated by scaling the non-bonded interaction, and when working with glycans in solution, we recommend a scaling factor of 0.85. On the other hand, when protein-binding is involved, free energy calculations with a wide variety of lectins revealed that only modest scaling was needed to achieve experimental *ΔG* values from SPR or ITC experiments. Thus, in initial studies of novel carbohydrates, we would in general recommend that the N-glycan parameters developed herein should be implemented with a non-bonded scaling factor of 0.9 for charged and/or highly polar complex type glycans whereas 0.95 is sufficient for the simpler high mannose type glycans. The parameters presented here should be useful for others interested in studying the role of glycans in the dynamics of various large glycoproteins and glycoprotein complexes which would benefit from a CG representation. Although the open beta version of Martini 3 has been released for phospholipid bilayers and proteins^51^, the bonded parameters and mapping schemes outlined in this study should be sufficiently robust for the future optimization of new compatible N-glycan parameters.

## Acknowledgments

ATS acknowledges NUS Research Scholarship funded by Ministry of Education (AcRF grant R-154-100-580-112). CSV, PJB, and PM acknowledge funding by the Ministry of Education of Singapore (MOE2012-T3-1-008). PJB and JKM further acknowledge funding from the National Research Foundation (NRF2017NRF-CRP001-027). We wish to thank the Singapore National Supercomputing Centre (https://www.nscc.sg) for computational resources.

## References

1. Cossart, P. & Sansonetti, P. J. Bacterial Invasion: The Paradigms of Enteroinvasive Pathogens. Science 304, 242–248 (2004).

2. Piwnica-Worms, D., Schuster, D. P. & Garbow, J. R. Molecular imaging of host-pathogen interactions in intact small animals. Cellular Microbiology 6, 319–331 (2004).

3. Bernimoulin, M. P. et al. Molecular basis of leukocyte rolling on PSGL-1: Predominant role of core-2 O-glycans and of tyrosine sulfate residue 51. J. Biol. Chem. 278, 37–47 (2003).

4. Vo, L. H., Yen, T.-Y., Macher, B. a & Hedrick, J. L. Identification of the ZPC oligosaccharide ligand involved in sperm binding and the glycan structures of Xenopus laevis vitelline envelope glycoproteins. Biol. Reprod. 69, 1822–30 (2003).

5. Green, R. S. et al. Mammalian N-Glycan Branching Protects against Innate Immune Self-Recognition and Inflammation in Autoimmune Disease Pathogenesis. Immunity 27, 308–320 (2007).

6. Johnson, C. P., Fujimoto, I., Rutishauser, U. & Leckband, D. E. Direct evidence that Neural Cell Adhesion Molecule (NCAM) polysialylation increases intermembrane repulsion and abrogates adhesion. J. Biol. Chem. 280, 137–145 (2005).

7. Langer, M. D., Guo, H., Shashikanth, N., Pierce, J. M. & Leckband, D. E. N-glycosylation alters cadherin-mediated intercellular binding kinetics. J. Cell Sci. 125, 2478–2485 (2012).

8. Moremen, K. W., Tiemeyer, M. & Nairn, A. V. Vertebrate protein glycosylation: Diversity, synthesis and function. Nature Reviews Molecular Cell Biology 13, 448–462 (2012).

9. Jaeken, J. Congenital disorders of glycosylation (CDG): It’s (nearly) all in it! Journal of Inherited Metabolic Disease 34, 853–858 (2011).

10. Michele, D. E. et al. Post-translational disruption of dystroglycan-ligand interactions in congenital muscular dystrophies. Nature 418, 417–422 (2002).

11. Yoshida, A. et al. Muscular Dystrophy and Neuronal Migration Disorder Caused by Mutations in a Glycosyltransferase, POMGnT1. Dev. Cell 1, 717–724 (2001).

12. Mcgowan, K. A. & Marinkovich, M. P. Laminins and human disease. Microscopy Research and Technique 51, 262–279 (2000).

13. Birklé, S., Zeng, G., Gao, L., Yu, R. K. & Aubry, J. Role of tumor-associated gangliosides in cancer progression. Biochimie 85, 455–463 (2003).

14. Stanley, P., Schachter, H. & Taniguchi, N. *N-Glycans*. Essentials of Glycobiology (Cold Spring Harbor Laboratory Press, 2009).

15. Shinkawa, T. et al. The absence of fucose but not the presence of galactose or bisecting N-acetylglucosamine of human IgG1 complex-type oligosaccharides shows the critical role of enhancing antibody-dependent cellular cytotoxicity. J. Biol. Chem. 278, 3466–3473 (2003).

16. Okazaki, A. et al. Fucose Depletion from Human IgG1 Oligosaccharide Enhances Binding Enthalpy and Association Rate between IgG1 and FcγRIIIa. J. Mol. Biol. 336, 1239–1249 (2004).

17. Hodoniczky, J., Yuan, Z. Z. & James, D. C. Control of recombinant monoclonal antibody effector functions by Fc N-glycan remodeling in vitro. Biotechnol. Prog. 21, 1644–1652 (2005).

18. Goldstein, I. J. & Hayes, C. E. The Lectins: Carbohydrate-Binding Proteins of Plants and Animals. Adv. Carbohydr. Chem. Biochem. 35, 127–340 (1978).

19. Misevic, G. N. & Burger, M. M. Reconstitution of high cell binding affinity of a marine sponge aggregation factor by cross-linking of small low affinity fragments into a large polyvalent polymer. J. Biol. Chem. 261, 2853–2859 (1986).

20. Del Valle, E. M. M. Cyclodextrins and their uses: A review. Process Biochem. 39, 1033–1046 (2004).

21. Varki, Ajit; Cummings, R. D.; Esko, J. D.; Freeze, H. H.; Stanley, P.; Bertozzi, C. R.; Hart, G. W.; Etzler, M. & E. *Essentials of Glycobiology, 3rd edition*. Cold Spring Harbor (NY) (Cold Spring Harbor Laboratory Press, 2015).

22. Chandler, K. B., Pompach, P., Goldman, R. & Edwards, N. Exploring site-specific N-glycosylation microheterogeneity of haptoglobin using glycopeptide CID tandem mass spectra and glycan database search. J. Proteome Res. 12, 3652–3666 (2013).

23. Mariño, K., Bones, J., Kattla, J. J. & Rudd, P. M. A systematic approach to protein glycosylation analysis: a path through the maze. Nat. Chem. Biol. 6, 713–723 (2010).

24. Rouvinski, A. et al. Recognition determinants of broadly neutralizing human antibodies against dengue viruses. Nature 520, 109–113 (2015).

25. Mesters, J. R. & Hilgenfeld, R. Protein glycosylation, sweet to crystal growth? in Crystal Growth and Design 7, 2251–2253 (American Chemical Society, 2007).

26. Han, L. & Costello, C. E. Mass spectrometry of glycans. Biochemistry. Biokhimiia 78, 710–20 (2013).

27. Leymarie, N. & Zaia, J. Effective use of mass spectrometry for glycan and glycopeptide structural analysis. Anal. Chem. 84, 3040–3048 (2012).

28. Kirschner, K. N. et al. GLYCAM06: A generalizable biomolecular force field. carbohydrates. J. Comput. Chem. 29, 622–655 (2008).

29. Hatcher, E., Guvench, O. & MacKerell, A. D. Charmm additive all-atom force field for aldopentofuranoses, methyl-aldopentofuranosides, and fructofuranose. J. Phys. Chem. B 113, 12466–12476 (2009).

30. Momany, F. A. & Willett, J. L. Computational studies on carbohydrates: In vacuo studies using a revised AMBER force field, AMB99C, designed for α-(1→4) linkages. Carbohydr. Res. 326, 194–209 (2000).

31. Pol-Fachin, L., Verli, H. & Lins, R. D. Extension and validation of the GROMOS 53A6(GLYC) parameter set for glycoproteins. J. Comput. Chem. 35, 2087–2095 (2014).

32. Allinger, N. L., Rahman, M. & Lii, J. H. A Molecular Mechanics Force Field (MM3) for Alcohols and Ethers. J. Am. Chem. Soc. 112, 8293–8307 (1990).

33. Dauchez, M., Derreumaux, P., Lagant, P. & Vergoten, G. A vibrational molecular force field of model compounds with biological interest. IV. Parameters for the different glycosidic linkages of oligosaccharides. J. Comput. Chem. 16, 188–199 (1995).

34. Momany, F. A., Willett, J. L. & Schnupf, U. Molecular dynamics simulations of a cyclic-DP-240 amylose fragment in a periodic cell: Glass transition temperature and water diffusion. Carbohydr. Polym. 78, 978–986 (2009).

35. Guvench, O., Hatcher, E., Venable, R. M., Pastor, R. W. & MacKerell, A. D. CHARMM additive all-atom force field for glycosidic linkages between hexopyranoses. J. Chem. Theory Comput. 5, 2353–2370 (2009).

36. Lins, R. D. & Hünenberger, P. H. A new GROMOS force field for hexopyranose-based carbohydrates. J. Comput. Chem. 26, 1400–1412 (2005).

37. Durier, V., Tristram, F. & Vergoten, G. Molecular force field development for saccharides using the SPASIBA spectroscopic potential. Force field parameters for α-D-glucose. J. Mol. Struct. THEOCHEM 395–396, 81–90 (1997).

38. Jo, S., Song, K. C., Desaire, H., MacKerell, A. D. & Im, W. Glycan reader: Automated sugar identification and simulation preparation for carbohydrates and glycoproteins. J. Comput. Chem. 32, 3135–3141 (2011).

39. Jo, S., Kim, T., Iyer, V. G. & Im, W. CHARMM-GUI: A web-based graphical user interface for CHARMM. J. Comput. Chem. 29, 1859–1865 (2008).

40. Woods, R. J. GLYCAM-Web | Utilities for molecular modeling of carbohydrates. Available at: http://glycam.org/tools/molecular-dynamics/glycoprotein-builder/upload-pdb. (Accessed: 8th January 2018)

41. Xiong, X. et al. Force fields and scoring functions for carbohydrate simulation. Carbohydrate Research 401, 73–81 (2015).

42. Sauter, J. & Grafmüller, A. Procedure for transferable coarse-grained models of aqueous polysaccharides. J. Chem. Theory Comput. 13, 223–236 (2017).

43. Sinitskiy, A. V. & Pande, V. S. Simulated Dynamics of Glycans on Ligand-Binding Domain of NMDA Receptors Reveals Strong Dynamic Coupling between Glycans and Protein Core. J. Chem. Theory Comput. 13, 5496–5505 (2017).

44. Marrink, S. J., Risselada, H. J., Yefimov, S., Tieleman, D. P. & De Vries, A. H. The MARTINI force field: Coarse grained model for biomolecular simulations. J. Phys. Chem. B 111, 7812–7824 (2007).

45. Monticelli, L. et al. The MARTINI coarse-grained force field: Extension to proteins. J. Chem. Theory Comput. 4, 819–834 (2008).

46. López, C. A. et al. Martini coarse-grained force field: Extension to carbohydrates. J. Chem. Theory Comput. 5, 3195–3210 (2009).

47. Uusitalo, J. J., Ingólfsson, H. I., Akhshi, P., Tieleman, D. P. & Marrink, S. J. Martini Coarse-Grained Force Field: Extension to DNA. J. Chem. Theory Comput. 11, 3932–3945 (2015).

48. Schmalhorst, P. S., Deluweit, F., Scherrers, R., Heisenberg, C. P. & Sikora, M. Overcoming the Limitations of the MARTINI Force Field in Simulations of Polysaccharides. J. Chem. Theory Comput. 13, 5039–5053 (2017).

49. Stark, A. C., Andrews, C. T. & Elcock, A. H. Toward optimized potential functions for protein-protein interactions in aqueous solutions: Osmotic second virial coefficient calculations using the MARTINI coarse-grained force field. J. Chem. Theory Comput. 9, 4176–4185 (2013).

50. Javanainen, M., Martinez-Seara, H. & Vattulainen, I. Excessive aggregation of membrane proteins in the Martini model. PLoS One 12, e0187936 (2017).

51. Marrink, S. J. Martini 3 open-beta release. Available at: http://cgmartini.nl/index.php/martini3beta. (Accessed: 16th October 2018)

52. Machatha, S. G. & Yalkowsky, S. H. Comparison of the octanol/water partition coefficients calculated by ClogP®, ACDlogP and KowWin® to experimentally determined values. Int. J. Pharm. 294, 185–192 (2005).

53. Vafaei, S., Tomberli, B. & Gray, C. G. McMillan-Mayer theory of solutions revisited: Simplifications and extensions. J. Chem. Phys. 141, 154501 (2014).

54. Schmid, N. et al. Definition and testing of the GROMOS force-field versions 54A7 and 54B7. Eur. Biophys. J. 40, 843–856 (2011).

55. Hess, B., Kutzner, C., Van Der Spoel, D. & Lindahl, E. GRGMACS 4: Algorithms for highly efficient, load-balanced, and scalable molecular simulation. J. Chem. Theory Comput. 4, 435–447 (2008).

56. Berendsen H.J.C., Postma J.P.M., van Gunsteren W.F. H. J. Interaction models for water in relation to protein hydration. in Intermolecular Forces 331–342 (Springer, Dordrecht, 1981). doi:10.1007/978-94-015-7658-1_21

57. Hünenberger, P. H. Thermostat algorithms for molecular dynamics simulations. Adv. Polym. Sci. 173, 105–147 (2005).

58. Berendsen, H. J. C., Postma, J. P. M., Van Gunsteren, W. F., Dinola, A. & Haak, J. R. Molecular dynamics with coupling to an external bath. J. Chem. Phys. 81, 3684–3690 (1984).

59. Hess, B., Bekker, H., Berendsen, H. J. C. & Fraaije, J. G. E. M. LINCS: A Linear Constraint Solver for molecular simulations. J. Comput. Chem. 18, 1463–1472 (1997).

60. Darden, T., Perera, L., Li, L. & Lee, P. New tricks for modelers from the crystallography toolkit: The particle mesh Ewald algorithm and its use in nucleic acid simulations. Structure 7, R55–60 (1999).

61. Huang, J. et al. CHARMM36m: An improved force field for folded and intrinsically disordered proteins. Nat. Methods (2016). doi:10.1038/nmeth.4067

62. Van Der Spoel, D. et al. GROMACS: Fast, flexible, and free. Journal of Computational Chemistry 26, 1701–1718 (2005).

63. López, C. A., Sovova, Z., Van Eerden, F. J., De Vries, A. H. & Marrink, S. J. Martini force field parameters for glycolipids. J. Chem. Theory Comput. 9, 1694–1708 (2013).

64. De Jong, D. H. et al. Improved parameters for the martini coarse-grained protein force field. J. Chem. Theory Comput. 9, 687–697 (2013).

65. Bussi, G., Donadio, D. & Parrinello, M. Canonical sampling through velocity rescaling. J. Chem. Phys. 126, 014101 (2007).

66. Tironi, I. G., Sperb, R., Smith, P. E. & Van Gunsteren, W. F. A generalized reaction field method for molecular dynamics simulations. J. Chem. Phys. 102, 5451–5459 (1995).

67. van Gunsteren, W. F. & Berendsen, H. J. C. Thermodynamic cycle integration by computer simulation as a tool for obtaining free energy differences in molecular chemistry. J. Comput. Aided. Mol. Des. 1, 171–176 (1987).

68. Beutler, T. C., Mark, A. E., van Schaik, R. C., Gerber, P. R. & van Gunsteren, W. F. Avoiding singularities and numerical instabilities in free energy calculations based on molecular simulations. Chem. Phys. Lett. 222, 529–539 (1994).

69. Bennett, C. H. Efficient estimation of free energy differences from Monte Carlo data. J. Comput. Phys. 22, 245–268 (1976).

70. Best, S. A., Merz, K. M. & Reynolds, C. H. Free energy perturbation study of octanol/water partition coefficients: Comparison with continuum GB/SA calculations. Journal of Physical Chemistry B 103, 714–726 (1999).

71. Torrie, G. M. & Valleau, J. P. Nonphysical sampling distributions in Monte Carlo free-energy estimation: Umbrella sampling. J. Comput. Phys. 23, 187–199 (1977).

72. Kumar, S., Rosenberg, J. M., Bouzida, D., Swendsen, R. H. & Kollman, P. A. THE weighted histogram analysis method for free-energy calculations on biomolecules. I. The method. J. Comput. Chem. 13, 1011–1021 (1992).

73. Doudou, S., Burton, N. A. & Henchman, R. H. Standard free energy of binding from a one-dimensional potential of mean force. J. Chem. Theory Comput. 5, 909–918 (2009).

74. Lay, W. K., Miller, M. S. & Elcock, A. H. Optimizing Solute-Solute Interactions in the GLYCAM06 and CHARMM36 Carbohydrate Force Fields Using Osmotic Pressure Measurements. J. Chem. Theory Comput. 12, 1401–1407 (2016).

75. Blanco, M. A., Sahin, E., Robinson, A. S. & Roberts, C. J. Coarse-grained model for colloidal protein interactions, B22, and protein cluster formation. J. Phys. Chem. B 117, 16013–16028 (2013).

76. Tessier, P. M. et al. Self-interaction chromatography: A novel screening method for rational protein crystallization. in Acta Crystallographica Section D: Biological Crystallography 58, 1531–1535 (2002).

77. Otwinowski, Z. & Minor, W. Macromolecular Crystallography Part A. Methods in Enzymology 276, (1997).

78. Harvey, D. J. et al. Proposal for a standard system for drawing structural diagrams of N- and O-linked carbohydrates and related compounds. Proteomics 9, 3796–3801 (2009).

79. Ioan, C. E., Aberle, T. & Burchard, W. Structure properties of dextran. 2. Dilute solution. Macromolecules 33, 5730–5739 (2000).

80. Naismith, J. H. & Field, R. A. Structural basis of trimannoside recognition by concanavalin A. J. Biol. Chem. 271, 972–976 (1996).

81. Mandal, D. K., Kishore, N. & Brewer, C. F. Thermodynamics of Lectin- Carbohydrate Interactions. Titration Microcalorimetry Measurements of the Binding of N-Linked Carbohydrates and Ovalbumin to Concanavalin A. Biochemistry 33, 1149–1156 (1994).

82. Boyd, M. R. et al. Discovery of cyanovirin-N, a novel human immunodeficiency virus-inactivating protein that binds viral surface envelope glycoprotein gp120: potential applications to microbicide development. Antimicrob. Agents Chemother. 41, 1521–30 (1997).

83. Bolmstedt, A. J., O’Keefe, B. R., Shenoy, S. R., McMahon, J. B. & Boyd, M. R. Cyanovirin-N defines a new class of antiviral agent targeting N-linked, high-mannose glycans in an oligosaccharide-specific manner. Mol. Pharmacol. 59, 949–54 (2001).

84. Shenoy, S. R., O’Keefe, B. R., Bolmstedt, A. J., Cartner, L. K. & Boyd, M. R. Selective interactions of the human immunodeficiency virus-inactivating protein cyanovirin-N with high-mannose oligosaccharides on gp120 and other glycoproteins. J. Pharmacol. Exp. Ther. 297, 704–710 (2001).

85. Botos, I. et al. Structures of the complexes of a potent anti-HIV protein cyanovirin-N and high mannose oligosaccharides. J. Biol. Chem. 277, 34336–34342 (2002).

86. Loris, R. et al. Structural Basis of Oligomannose Recognition by the Pterocarpus angolensis Seed Lectin. J. Mol. Biol. 335, 1227–1240 (2004).

87. Loris, R. et al. Crystal structure of Pterocarpus angolensis lectin in complex with glucose, sucrose, and turanose. J. Biol. Chem. 278, 16297–16303 (2003).

88. Garcia-Pino, A., Buts, L., Wyns, L., Imberty, A. & Loris, R. How a plant lectin recognizes high mannose oligosaccharides. Plant Physiol. 144, 1733–1741 (2007).

89. Green, E. D., Brodbeck, R. M. & Baenziger, J. U. Lectin affinity high-performance liquid chromatography. Interactions of N-glycanase-released oligosaccharides with Ricinus communis agglutinin I and Ricinus communis agglutinin II. J. Biol. Chem. 262, 12030–9 (1987).

90. Wu, A. M. et al. Recognition factors of Ricinus communis agglutinin 1 (RCA1). Mol. Immunol. 43, 1700–1715 (2006).

91. Shinohara, Y. et al. Use of a biosensor based on surface plasmon resonance and biotinyl glycans for analysis of sugar binding specificities of lectins. J. Biochem. 117, 1076–82 (1995).

92. Asensio, J. L. et al. Structural basis for chitin recognition by defense proteins: GIcNAc residues are bound in a multivalent fashion by extended binding sites in hevein domains. Chem. Biol. 7, 529–543 (2000).

93. Mirelman, D., Galun, E., Sharon, N. & Lotan, R. Inhibition of fungal growth by wheat germ agglutinin. Nature 256, 414–416 (1975).

94. Harata, K., Nagahora, H. & Jigami, Y. X-ray structure of wheat germ agglutinin isolectin 3. Acta Crystallogr. Sect. D Biol. Crystallogr. 51, 1013–1019 (1995).

95. Schwefel, D. et al. Structural basis of multivalent binding to wheat germ agglutinin. J. Am. Chem. Soc. 132, 8704–8719 (2010).

96. Rice, A. C. Cloning, expression, purification and characterization of domain B of wheat germ agglutinin. (Virginia Commonwealth University, 1994).

97. Shibuya, N., Goldstein, I. J., Shafer, J. A., Peumans, W. J. & Broekaert, W. F. Carbohydrate binding properties of the stinging nettle (Urtica dioica) rhizome lectin. Arch. Biochem. Biophys. 249, 215–224 (1986).

98. Peumans, W. J., De Ley, M. & Broekaert, W. F. An unusual lectin from stinging nettle (Urtica dioica) rhizomes. FEBS Lett. 177, 99–103 (1984).

99. Huesing, J. E., Murdock, L. L. & Shade, R. E. Rice and stinging nettle lectins: Insecticidal activity similar to wheat germ agglutinin. Phytochemistry 30, 3565–3568 (1991).

100. Broekaert, W. F., Van Parijs, J. A. N., Leyns, F., Joos, H. & Peumans, W. J. A chitin-binding lectin from stinging nettle rhizomes with antifungal properties. Science (80-.). 245, 1100–1102 (1989).

101. Harata, K. & Muraki, M. Crystal structures of Urtica dioica agglutinin and its complex with tri-N-acetylchitotriose. J. Mol. Biol. 297, 673–681 (2000).

102. Kawaguchi, T., Matsumoto, I. & Osawa, T. Studies on Hemagglutinins from Maackia amurensis seeds. J. Biol. Chem. 249, 2786–2792 (1974).

103. Kawaguchi, T. & Osawa, T. Elucidation of Lectin Receptors by Quantitative Inhibition of Lectin Binding to Human Erythrocytes and Lymphocytes. Biochemistry (1976). doi:10.1021/bi00666a006

104. Knibbs, R. N., Goldstein, I. J., Ratcliffe, R. M. & Shibuya, N. Characterization of the carbohydrate binding specificity of the leukoagglutinating lectin from Maackia amurensis. Comparison with other sialic acid-specific lectins. J. Biol. Chem. 266, 83–88 (1991).

105. Wang, W. C. & Cummings, R. D. The immobilized leukoagglutinin from the seeds of Maackia amurensis binds with high affinity to complex-type Asn-linked oligosaccharides containing terminal sialic acid-linked alpha-2,3 to penultimate galactose residues. J. Biol. Chem. 263, 4576–4585 (1988).

106. Haseley, S. R., Talaga, P., Kamerling, J. P. & Vliegenthart, J. F. G. Characterization of the carbohydrate binding specificity and kinetic parameters of lectins by using surface plasmon resonance. Anal. Biochem. 274, 203–210 (1999).

107. Imberty, A. et al. An unusual carbohydrate binding site revealed by the structures of two Maackia amurensis lectins complexed with sialic acid-containing oligosaccharides. J. Biol. Chem. 275, 17541–17548 (2000).

108. Ramadugu, S. K., Chung, Y. H., Xia, J. & Margulis, C. J. When sugars get wet. A comprehensive study of the behavior of water on the surface of oligosaccharides. J. Phys. Chem. B 113, 11003–11015 (2009).

109. Chebaro, Y., Pasquali, S. & Derreumaux, P. The coarse-grained OPEP force field for non-amyloid and amyloid proteins. J. Phys. Chem. B 116, 8741–8752 (2012).

110. Maupetit, J., Tuffery, P. & Derreumaux, P. A coarse-grained protein force field for folding and structure prediction. Proteins Struct. Funct. Bioinforma. 69, 394–408 (2007).

111. Bereau, T. & Deserno, M. Generic coarse-grained model for protein folding and aggregation. J. Chem. Phys. (2009). doi:10.1063/1.3152842

112. Liwo, A. et al. A united-residue force field for off-lattice protein-structure simulations. I. Functional forms and parameters of long-range side-chain interaction potentials from protein crystal data. J. Comput. Chem. 18, 849–873 (1997).

113. Pol-Fachin, L., Rusu, V. H., Verli, H. & Lins, R. D. GROMOS 53A6 GLYC, an improved GROMOS force field for hexopyranose-based carbohydrates. J. Chem. Theory Comput. 8, 4681–4690 (2012).

114. Kaufmann, M., Mügge, C. & Kroh, L. W. NMR analyses of complex d -glucose anomerization. Food Chem. 265, 222–226 (2018).

115. Shwe, T., Pratchayasakul, W., Chattipakorn, N. & Chattipakorn, S. C. Role of D-galactose-induced brain aging and its potential used for therapeutic interventions. Exp. Gerontol. 101, 13–36 (2018).

116. Huang, J. et al. Production of d-mannose from d-glucose by co-expression of d-glucose isomerase and d-lyxose isomerase in Escherichia coli. J. Sci. Food Agric. 98, 4895–4902 (2018).

117. Ohrui, H., Nishida, Y. & Higuchi, H. The preferred rotamer about the C5-C6 bond of D-galactopyranoses and the stereochemistry of dehydrogenation by D-galactose oxidase. Can. J. Chem. 65, 1145–1153 (1987).

118. Nishida, Y., Ohrui, H. & Meguro, H. 1H-NMR studies of (6r)- and (6s)- deuterated d-hexoses: assignment of the preferred rotamers about C5-C6 bond of D-glucose and D-galactose derivatives in solutions. Tetrahedron Lett. 25, 1575–1578 (1984).

119. Wolff, D. & Rudd, W. G. Tabulated potentials in molecular dynamics simulations. Comput. Phys. Commun. 120, 20–32 (1999).

120. Feenstra, K. A., Hess, B. & Berendsen, H. J. C. Improving efficiency of large time-scale molecular dynamics simulations of hydrogen-rich systems. J. Comput. Chem. 20, 786 (1999).

121. Alessandri, R. et al. Pitfalls of the Martini Model. J. Chem. Theory Comput. 15, 5448–5460 (2019).

122. Lis, H. & Sharon, N. Lectins: Carbohydrate-specific proteins that mediate cellular recognition. Chem. Rev. 98, 637–674 (1998).

123. Chavent, M., Chetwynd, A. P., Stansfeld, P. J. & Sansom, M. S. P. Dimerization of the EphA1 receptor tyrosine kinase transmembrane domain: Insights into the mechanism of receptor activation. Biochemistry 53, 6641–6652 (2014).

124. Artemenko, E. O., Egorova, N. S., Arseniev, A. S. & Feofanov, A. V. Transmembrane domain of EphA1 receptor forms dimers in membrane-like environment. Biochim. Biophys. Acta - Biomembr. 1778, 2361–2367 (2008).

125. Prakash, A., Janosi, L. & Doxastakis, M. Self-association of models of transmembrane domains of erbb receptors in a lipid bilayer. Biophys. J. 99, 3657–3665 (2010).

126. Chen, L., Merzlyakov, M., Cohen, T., Shai, Y. & Hristova, K. Energetics of ErbB1 transmembrane domain dimerization in lipid bilayers. Biophys. J. 96, 4622–4630 (2009).

127. Gabdoulkhakov, A. G. et al. Structure-function investigation complex of Agglutinin from Ricinus communis with galactoaza. TO BE PUBLISHED Available at: https://www.rcsb.org/structure/1rzo. (Accessed: 23rd April 2018)

