## Supplementary information for "Extending the Martini coarse-grained forcefield to N-glycans"

### Extending the Martini coarse-grained force field to N-glycans – Supplementary information

**Table S1: Bonded parameters for all the di/trisaccharides.** Bonds, angles and dihedral parameters are shown according to mapping in Figure 1. For angles greater than 140°, the restricted bending potential (ReB) was used. Dihedral angles marked with * were optimized with a multiplicity of 2.

| Di/Trisaccharide | Bonds | Rbond (nm) | Kbond (kJ $mol^{-1}$ $nm^{-2}$) | Angles | Θ (deg) | Kangle (kJ $mol^{-1}$) | Dihedrals | Φpd (°) | Kpd (kJ $mol^{-1}$) |
|---|---|---|---|---|---|---|---|---|---|
| N-acetylglucosamine-β-(1, 4)-N-acetylglucosamine (A) | G1 - G2 | 0.238 | 30000 | G1 - G2 - G3 | 135 | 300 | G3 - G2 - G6 - G7 | 45 | 10 |
| | G2 - G3 | 0.222 | 30000 | G2 - G3 - G4 | 145 | 250 | G1 - G2 - G3 - G4 | 180 | 15 |
| | G3 - G4 | 0.312 | 30000 | G5 - G6 - G7 | 135 | 300 | G5 - G6 - G7 - G8 | 180 | 15 |
| | G2 - G6 | 0.524 | 5000 | G6 - G7 - G8 | 145 | 250 | | | |
| | G5 - G6 | 0.238 | 30000 | G2 - G6 - G5 | 70 | 240 | | | |
| | G6 - G7 | 0.222 | 30000 | G2 - G6 - G7 | 74 | 370 | | | |
| | G7 - G8 | 0.312 | 30000 | G1 - G2 - G6 | 110 | 100 | | | |
| | | | | G3 - G2 - G6 | 100 | 145 | | | |
| N-acetylglucosamine-Asparagine (B) | A1 - A2 | 0.320 | 5000 | A1 - A2 - G2 | 115 | 170 | A1 - A2 - G2 - G1 | 80 | 8 |
| | A2 - G2 | 0.474 | 30000 | G1 - G2 - A2 | 45 | 50 | A1 - A2 - G2 - G3 | 300 | 8 |
| | G1 - G2 | 0.238 | 30000 | A2 - G2 - G3 | 35 | 120 | G1 - G2 - G3 - G4 | 180 | 15 |
| | G2 - G3 | 0.222 | 30000 | G1 - G2 - G3 | 135 | 300 | | | |
| | G3 - G4 | 0.312 | 30000 | G2 - G3 - G4 | 145 | 250 | | | |
| Fucose-α-(1,6)-N-acetylglucosamine (C) | G1 - G2 | 0.238 | 30000 | G1 - G2 - G3 | 135 | 300 | G1 - G2 - G3 - G4 | 180 | 15 |
| | G2 - G3 | 0.222 | 30000 | G2 - G3 - G4 | 145 | 250 | F2 - G1 - G2 - G3 | 30 | 7 |
| | G3 - G4 | 0.312 | 30000 | F2 - G1 - G2 | 145 | 100 | F3 - F2 - G1 - G2* | 180 | 5 |
| | G1 - F2 | 0.499 | 5000 | F3 - F2 - G1 | 82 | 180 | | | |
| | F2 - F3 | 0.224 | 30000 | F1 - F2 - G1 | 70 | 180 | | | |
| | F1 - F2 | 0.220 | 30000 | | | | | | |
| Mannose-β-(1, 4)-N-Acetylglucosamine (D) | G1 - G2 | 0.238 | 30000 | G1 - G2 - G3 | 135 | 300 | G1 - G2 - G3 - G4 | 180 | 15 |
| | G2 - G3 | 0.222 | 30000 | G2 - G3 - G4 | 145 | 250 | G1 - G2 - M2 - M3 | 0 | 8 |
| | G3 - G4 | 0.312 | 30000 | G1 - G2 - M2 | 95 | 80 | G1 - G2 - M2 - M1 | 95 | 7 |
| | G2 - M2 | 0.474 | 12000 | G3 - G2 - M2 | 125 | 140 | | | |
| | M1 - M2 | 0.245 | 30000 | M1 - M2 - G2 | 77 | 230 | | | |
| | M2 - M3 | 0.222 | 30000 | G2 - M2 - M3 | 82 | 240 | | | |
| | | | | M1 - M2 - M3 | 132 | 220 | | | |
| Mannose-α-(1, 3)-[Mannose-α-(1, 6)]-Mannose (E) | M4 - M5 | 0.230 | 30000 | M1 - M2 - M3 | 132 | 220 | M3 - M2 - M4 - M5 | 0 | 15 |
| | M5 - M6 | 0.222 | 30000 | M4 - M5 - M6 | 132 | 220 | M5 - M6 - M8 - M9 | 340 | 13 |
| | M6 - M8 | 0.485 | 8500 | M7 - M8 - M9 | 132 | 220 | M4 - M5 - M6 - M8 | 260 | 20 |

| | | | | | | | | | |
|---|---|---|---|---|---|---|---|---|---|
| | M8 - M9 | 0.222 | 30000 | M3 - M2 - M4 | 84 | 170 | M2 - M4 - M5 - M6 | 200 | 7 |
| | M7 - M8 | 0.230 | 30000 | M1 - M2 - M4 | 74 | 60 | | | |
| | M2 - M4 | 0.465 | 3000 | M6 - M8 - M9 | 68 | 210 | | | |
| | M2 - M3 | 0.222 | 30000 | M6 - M8 - M7 | 77 | 70 | | | |
| | M1 - M2 | 0.230 | 30000 | M2 - M4 - M5 | 157 | 60 | | | |
| | | | | M5 - M6 - M8 | 110 | 70 | | | |
| N-Acetylglucosamine-β-(1, 2)-Mannose (F) | M1 - M2 | 0.230 | 30000 | M1 - M2 -M3 | 132 | 220 | M1 - M2 - M3 - G2 | 55 | 25 |
| | M2 - M3 | 0.222 | 30000 | M2 - M3 - G2 | 110 | 150 | M2 - M3 - G2 - G3 | -25 | 17 |
| | M3 - G2 | 0.535 | 17000 | G1 - G2 - M3 | 67 | 300 | G1 - G2 - G3 - G4 | 180 | 15 |
| | G1 - G2 | 0.238 | 30000 | M3 - G2 - G3 | 70 | 400 | | | |
| | G2 - G3 | 0.222 | 30000 | G2 - G3 - G4 | 145 | 250 | | | |
| | G3 - G4 | 0.312 | 30000 | G1 - G2 - G3 | 135 | 300 | | | |
| Galactose-β-(1, 4)-N-Acetylglucosamine (G) | G1 - G2 | 0.238 | 30000 | G1 - G2 - G3 | 135 | 300 | G1 - G2 - G3 - G4 | 180 | 15 |
| | G2 - G3 | 0.222 | 30000 | G2 - G3 - G4 | 145 | 250 | G1 - G2 - GL2 - GL3 | 245 | 5 |
| | G3 - G4 | 0.312 | 30000 | GL1 - GL2 - GL3 | 140 | 170 | | | |
| | G2 - GL2 | 0.495 | 14000 | G2 - GL2 - GL1 | 80 | 160 | | | |
| | GL1 - GL2 | 0.222 | 30000 | G2 - GL2 - GL3 | 78 | 250 | | | |
| | GL2 - GL3 | 0.222 | 30000 | GL2 - G2 - G3 | 100 | 70 | | | |
| | | | | G1 - G2 - GL2 | 115 | 70 | | | |
| N-Acetylneuraminic acid-β-(1, 2)-Galactose (H) | GL1 - GL2 | 0.222 | 30000 | GL3 - S1 - S4 | 40 | 80 | GL3 - S1 - S4 - S5 | 90 | 5 |
| | GL2 - GL3 | 0.222 | 30000 | GL3 - S1 - S2 | 80 | 150 | GL3 - S1 - S2 - S3 | 120 | 15 |
| | GL3 - S1 | 0.300 | 2500 | S1 - S4 - S5 | 130 | 40 | GL2 - GL3 - S1 - S2 | 40 | 17 |
| | S1 - S4 | 0.370 | 3000 | S1 - S4 - S2 | 100 | 80 | GL1 - GL2 - GL3 - S1 | 100 | 25 |
| | S2 - S3 | 0.350 | 12000 | S1 - S2 - S4 | 50 | 180 | | | |
| | S4 - S5 | 0.300 | 7000 | S2 - S4 - S5 | 150 | 20 | | | |
| | | | | S4 - S1 - SS2 | 45 | 20 | | | |
| | S1 - S2 | 0.290 | CONSTRAINT | S1 - S2 - S3 | 140 | 230 | | | |
| | S2 - S4 | 0.260 | CONSTRAINT | S4 - S2 - S3 | 60 | 50 | | | |
| | | | | GL1 - GL2 - GL3 | 140 | 170 | | | |
| Mannose-α-(1, 2)-Mannose (I) | M1 - M2 | 0.230 | 30000 | M1 - M2 - M3 | 132 | 220 | M2 - M3 - M5 - M6 | 0 | 5 |
| | M2 - M3 | 0.222 | 30000 | M4 - M5 - M6 | 132 | 220 | M1 - M2 - M3 - M5 | 135 | 60 |
| | M3 - M5 | 0.478 | 5000 | M2 - M3 - M5 | 120 | 150 | | | |
| | M4 - M5 | 0.230 | 30000 | M3 - M5 - M4 | 85 | 120 | | | |
| | M5 - M6 | 0.222 | 30000 | M3 - M5 - M6 | 50 | 60 | | | |
| | M1 - M2 | 0.230 | 30000 | M1 - M2 - M3 | 117 | 220 | G1 - G2 - G3 - G4 | 180 | 15 |

| | | | | | | | | | |
|---|---|---|---|---|---|---|---|---|---|
| N-Acetylglucosamine-β-(1, 4)-Mannose (J) | M2 - M3 | 0.222 | 30000 | G1 - G2 - G3 | 135 | 300 | M3 - M2 - G2 - G3 | 45 | 20 |
| | M2 - G2 | 0.440 | 10000 | G2 - G3 - G4 | 145 | 250 | | | |
| | G1 - G2 | 0.238 | 30000 | G1 - G2 - M2 | 30 | 25 | | | |
| | G2 - G3 | 0.222 | 30000 | M2 - G2 - G3 | 20 | 40 | | | |
| | G3 - G4 | 0.312 | 30000 | M1 - M2 - G2 | 150 | 180 | | | |
| | | | | G2 - M2 - M3 | 130 | 90 | | | |
| N-Acetylglucosamine-β-(1, 6)-Mannose (K) | M1 - M2 | 0.230 | 30000 | M1 - M2 - M3 | 132 | 220 | G1 - G2 - G3 - G4 | 180 | 15 |
| | M2 - M3 | 0.222 | 30000 | G2 - M1 - M2 | 155 | 180 | G3 - G2 - M1 - M2 | 345 | 12 |
| | M1 - G2 | 0.534 | 10000 | G1 - G2 - M1 | 70 | 240 | G2 - M1 - M2 - M3 | 100 | 9 |
| | G1 - G2 | 0.238 | 30000 | G3 - G2 - M1 | 68 | 550 | | | |
| | G2 - G3 | 0.222 | 30000 | G1 - G2 - G3 | 135 | 300 | | | |
| | G3 - G4 | 0.312 | 30000 | G2 - G3 - G4 | 145 | 250 | | | |

**Table S2: Bead type selection for N-glycans.** Bead types used for each disaccharide in Figure 1 are shown.

| | | | | | | | | | | |
|---|---|---|---|---|---|---|---|---|---|---|
| N-acetylglucosamine-β-(1, 4)-N-acetylglucosamine (A) | Name | G1 | G2 | G3 | G4 | G5 | G6 | G7 | G8 | |
| | Type | P1 | SP2 | P3 | SP1 | P1 | SP2 | P3 | SP1 | |
| N-acetylglucosamine-Aspargine (B) | Name | G1 | G2 | G3 | G4 | A1 | A2 | | | |
| | Type | P1 | SP2 | P3 | SP1 | P5 | P5 | | | |
| Fucose-α-(1,6)-N-acetylglucosamine (C) | Name | G1 | G2 | G3 | G4 | F1 | F2 | F3 | | |
| | Type | P1 | SP2 | P3 | SP1 | SP1 | SP2 | P4 | | |
| Mannose-β-(1, 4)-N-Acetylglucosamine (D) | Name | G1 | G2 | G3 | G4 | M1 | M2 | M3 | | |
| | Type | P1 | SP2 | P3 | SP1 | P1 | SP2 | P4 | | |
| Mannose-α-(1, 3)-[Mannose-α-(1, 6)]-Mannose (E) | Name | M1 | M2 | M3 | M4 | M5 | M6 | M7 | M8 | M9 |
| | Type | P1 | SP2 | P4 | P1 | SP2 | P4 | P1 | SP2 | P4 |
| N-Acetylglucosamine-β-(1, 2)-Mannose (F) | Name | M1 | M2 | M3 | G1 | G2 | G3 | G4 | | |
| | Type | P1 | SP2 | P4 | P1 | SP2 | P3 | SP1 | | |
| Galactose-β-(1, 4)-N-Acetylglucosamine (G) | Name | GL1 | GL2 | GL3 | G1 | G2 | G3 | G4 | | |
| | Type | P1 | SP2 | P4 | P1 | SP2 | P3 | SP1 | | |
| N-Acetylneuraminic acid-β-(1, 2)-Galactose (H) | Name | S1 | S2 | S3 | S4 | S5 | GL1 | GL2 | GL3 | |
| | Type | Qa | SP1 | P3 | SP1 | P4 | P1 | SP2 | P4 | |
| Mannose-α-(1, 2)-Mannose (I) | Name | M1 | M2 | M3 | M4 | M5 | M6 | | | |
| | Type | P1 | SP2 | P4 | P1 | SP2 | P4 | | | |
| N-Acetylglucosamine-β-(1, 4)-Mannose (J) | Name | M1 | M2 | M3 | G1 | G2 | G3 | G4 | | |
| | Type | P1 | SP2 | P4 | P1 | SP2 | P3 | SP1 | | |
| N-Acetylglucosamine-β-(1, 6)-Mannose (K) | Name | M1 | M2 | M3 | G1 | G2 | G3 | G4 | | |
| | Type | P1 | SP2 | P4 | P1 | SP2 | P3 | SP1 | | |

**Table S3: Lectins with their residues interacting with glycans.** In the lectin, column, their names are colored according to their specificity: mannose specific lectins (green), galactose specific lectins (blue), and sialic acid/N-acetylglucosamine specific lectins (red). In the residues column, charged residues are indicated in red colour. Monosaccharides present in the glycan column are represented by their symbolic representation, including mannose (green circle), N-acetylglucosamine (blue square), galactose (yellow circle), and Neu5Ac/sialic acid (purple diamond).

| Lectin | Pocket Residues (Charged in Red) | No of charged residues | No of polar residues | Glycan | Scaling factor (λ) | $\Delta G^0_{sim}$ (kcal / mol) | $\Delta G^0_{Expt}$ (kcal / mol) |
|---|---|---|---|---|---|---|---|
| Cyanovirin-N (CVN) (PDB: 3GXZ) | L1, G2, **K3**, T7, E23, N93, **D95**, **E101** | 3 | 2 | 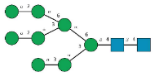 | 1.0 | -9.8 ± 0.7 | -8.7 ± 0.1[1] |
| | | | | | 0.95 | -7.5 ± 0.4 | |
| | | | | 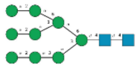 | 1.0 | -10.9 ± 0.4 | -9.2 ± 0.3[1] |
| | | | | | 0.95 | -8.6 ± 0.4 | |
| Pterocarpus angolensis (PAL) (PDB: 2PHW) | N83, **D86**, G106, **D136**, S137, N138, **E221**, Q222 | 3 | 4 | 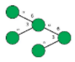 | 1.0 | -9.3 ± 0.6 | -5.2[2] |
| | | | | | 0.95 | -6.3 ± 0.8 | |
| | | | | | 0.9 | -4.9 ± 0.6 | |
| | | | | 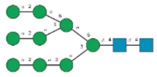 | 1.0 | -14.8 ± 1.1 | -5.8[2] |
| | | | | | 0.95 | -10.0 ± 1.0 | |
| | | | | | 0.9 | -7.6 ± 1.2 | |
| Maackia Amurensis (MAA) (PDB: 1DBN) | Y45, **D87**, S104, S106, **K107**, Y131, **D137**, **E224** | 4 | 2 | 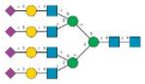 | 1.0 | - 14.9 ± 1.7 | -5.5[3] |
| | | | | | 0.95 | -10.4 ± 1.5 | |
| | | | | | 0.9 | -7.9 ± 0.9 | |
| | | | | 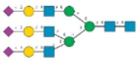 | 1.0 | -12.9 ± 1.6 | -5.7[3] |
| | | | | | 0.95 | -8.3 ± 0.9 | |
| | | | | | 0.9 | -7.1 ± 0.9 | |
| | | | | 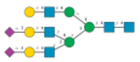 | 1.0 | -14.5 ± 1.6 | -4.7[3] |
| | | | | | 0.95 | -10.3 ± 0.7 | |
| | | | | | 0.9 | -7.2 ± 0.8 | |
| Ricinus communis (RCA) (PDB: 1RZO) | **D2022**, G2025, **E2026**, Q2035, **K2040**, N2046 | 3 | 2 | 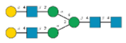 | 1.0 | -13.6 ± 1.1 | -7.7 ± 0.1[4] |
| | | | | | 0.95 | -10.5 ± 0.8 | |
| | | | | | 0.9 | -6.1 ± 0.7 | |
| | | | | 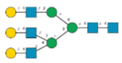 | 1.0 | -15.9 ± 0.3 | -7.3[4] |
| | | | | | 0.95 | -10.2 ± 0.9 | |
| | | | | | 0.9 | -6.3 ± 0.7 | |
| | | | | 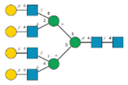 | 1.0 | -13.7 ± 0.2 | -7.0[4] |
| | | | | | 0.95 | -9.2 ± 0.3 | |
| | | | | | 0.9 | -6.3 ± 0.3 | |

| | | | | | | | |
|---|---|---|---|---|---|---|---|
| Concanavalin A (CONA) (PDB: 1CVN) | Y12, N14, T15, **D16**, G98, L99, Y100, A207, **D208**, G227, **R228** | 3 | 2 | 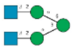 | 1.0 | -9.5 ± 0.2 | -8.4 ± 0.1[5] |
| | | | | | 0.95 | -7.2 ± 0.9 | |
| Wheat Germ Agglutinin (WGA) (PDB: 2UVO) | S62, **E72**, Y73, **D86**, S105, S114, **E115** | 3 | 2 |  | 1.0 | -8.5 ± 0.5 | -5.8[5] |
| | | | | | 0.95 | -5.6 ± 0.5 | |
| Urtica Dioicia Agglutinin (UDA) (PDB: 1EHH) | S19, C24, Y30 | 0 | 1 | 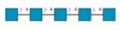 | 1.0 | -7.4 ± 0.8 | -5.9[5] |
| | | | | | 0.95 | -5.5 ± 0.4 | |

**Table S4: Residues and their Martini bead composition at each of the interacting glycan surfaces for PMFs in Urtica dioica agglutinin, shown in Figure S11.**

| Surface type | Favourable | Unfavourable 1 | Unfavourable 2 | Unfavourable 3 |
|---|---|---|---|---|
| Residues | C24, S19, W21, Y30 | S5, Q6, I20, E36 | G79, G80, C82, Q83, S89 | S45, D46, H47, W69 |
| Martini sidechain bead types for above residues | SC4, SC4, SP1, C5, P1, SC4, SNd, SC5, SC5 | P1, P4, AC1, Qa | C5, P4, P1 | P1, Qa, P5, SC4, SP1, SP1, P5, SC4, SNd, SC5 SC5 |

**Table S5: Window spacing used for thermodynamic integration calculations for coulombic and Van der Walls components.**

coul-lambdas/vdw-lambdas = 0 0.03 0.06 0.09 0.12 0.15 0.18
0.21 0.24 0.27 0.3 0.33 0.36 0.39 0.42 0.45 0.48 0.51
0.54 0.57 0.6 0.63 0.66 0.69 0.72 0.75 0.78 0.81 0.84
0.87 0.9 0.93 0.96 0.962 0.964 0.966 0.968 0.97 0.972 0.974
0.976 0.978 0.98 0.982 0.984 0.986 0.988 0.99 0.992 0.994 0.996
0.998 0.9985 0.999 0.9995

**Figure S1: Bonded distributions (CG vs ATM).** Distributions are shown for all the disaccharide/trisaccharide combinations observed in the common N-glycans: (A) N-acetylglucosamine-β(1,4)-N-acetylglucosamine; (B) N-acetylglucosamine-β(1,)-asparagine; (C) fucose-α-(1,6)-N-acetylglucosamine; (D) mannose-β-(1,4)-N-acetylglucosamine; (E) mannose-α-(1,3)-[mannose-α-(1,6)]-mannose; (F) N-acetylglucosamine-β-(1,2)-mannose; (G) galactose-β-(1,4)-N-acetylglucosamine; (H) N-acetylneuraminic acid-β-(1,2)-galactose; (I) mannose-α-(1,2)-mannose; (J) N-acetylglucosamine-β(1, 4)-mannose; and (K) N-acetylglucosamine-β(1, 6)-mannose.

#### A) N-Acetylglucosamine-β(1,4)-N-Acetylglucosamine:

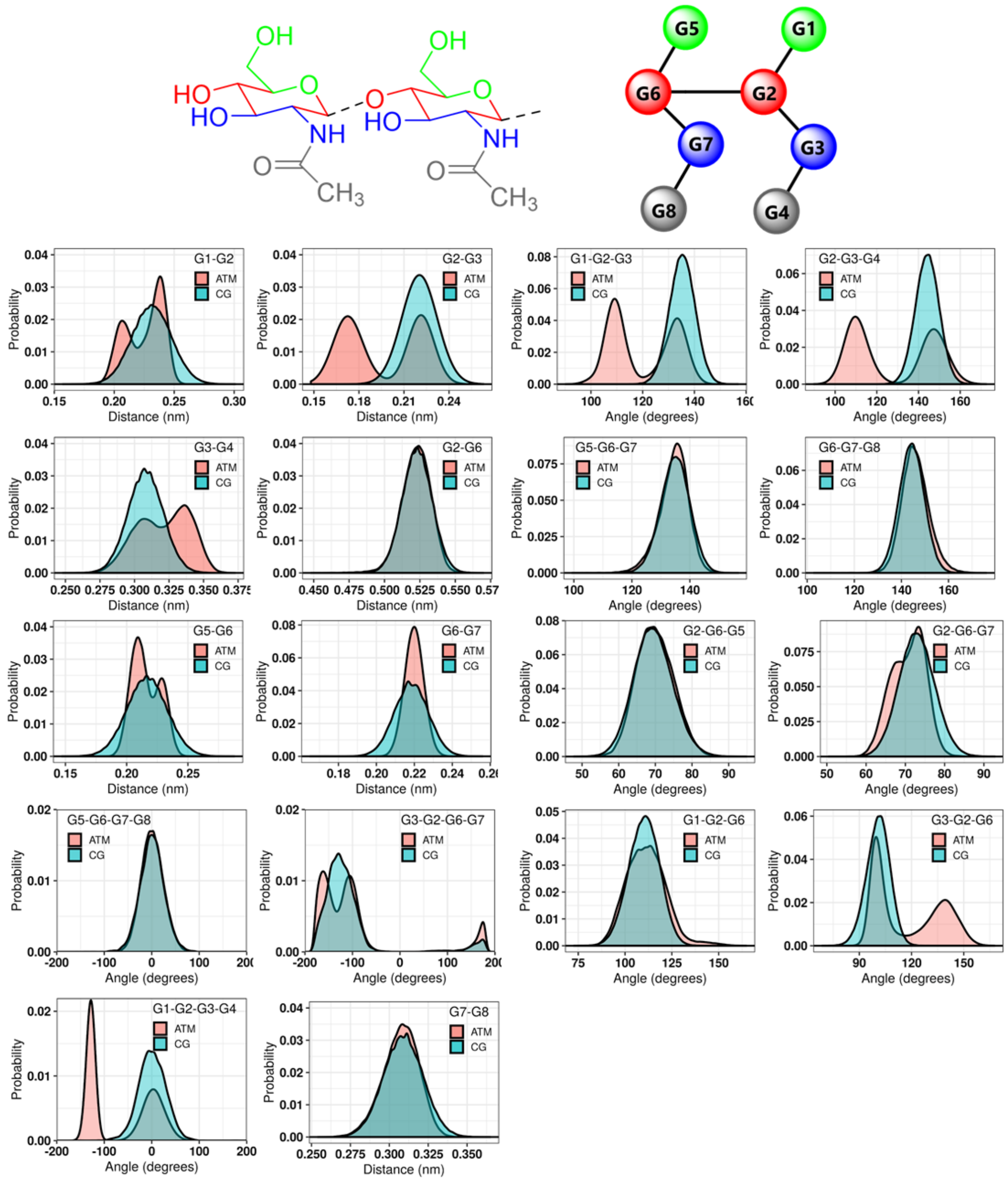

B) N-Acetylglucosamine-β(1,)-Asparagine:

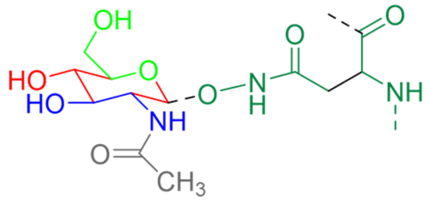

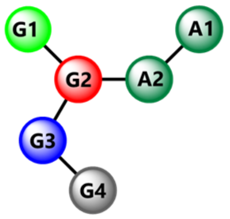

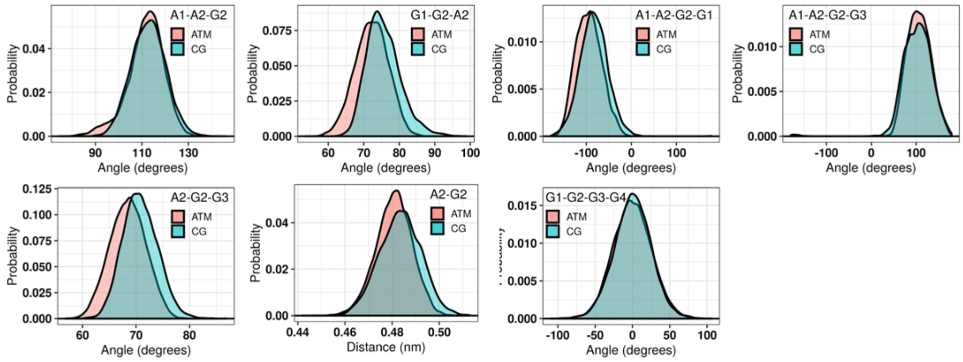

C) Fucose-α(1,6)-N-Acetylglucosamine:

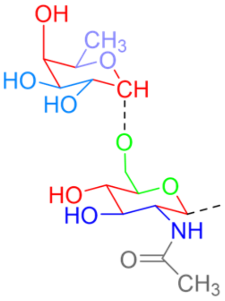

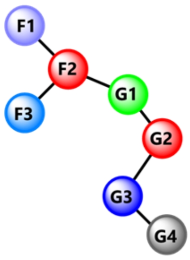

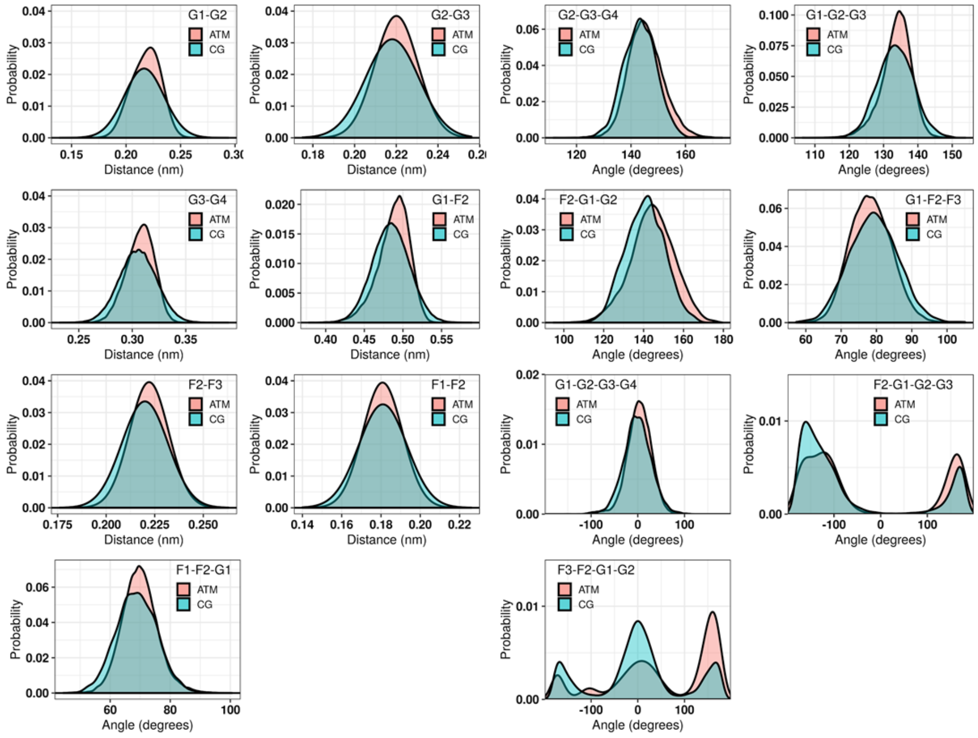

D) Mannose-β(1, 4)-N-Acetylglucosamine

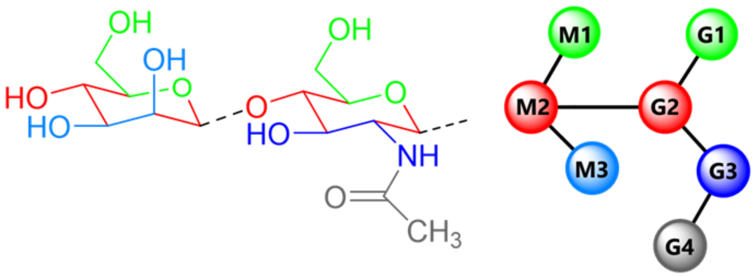

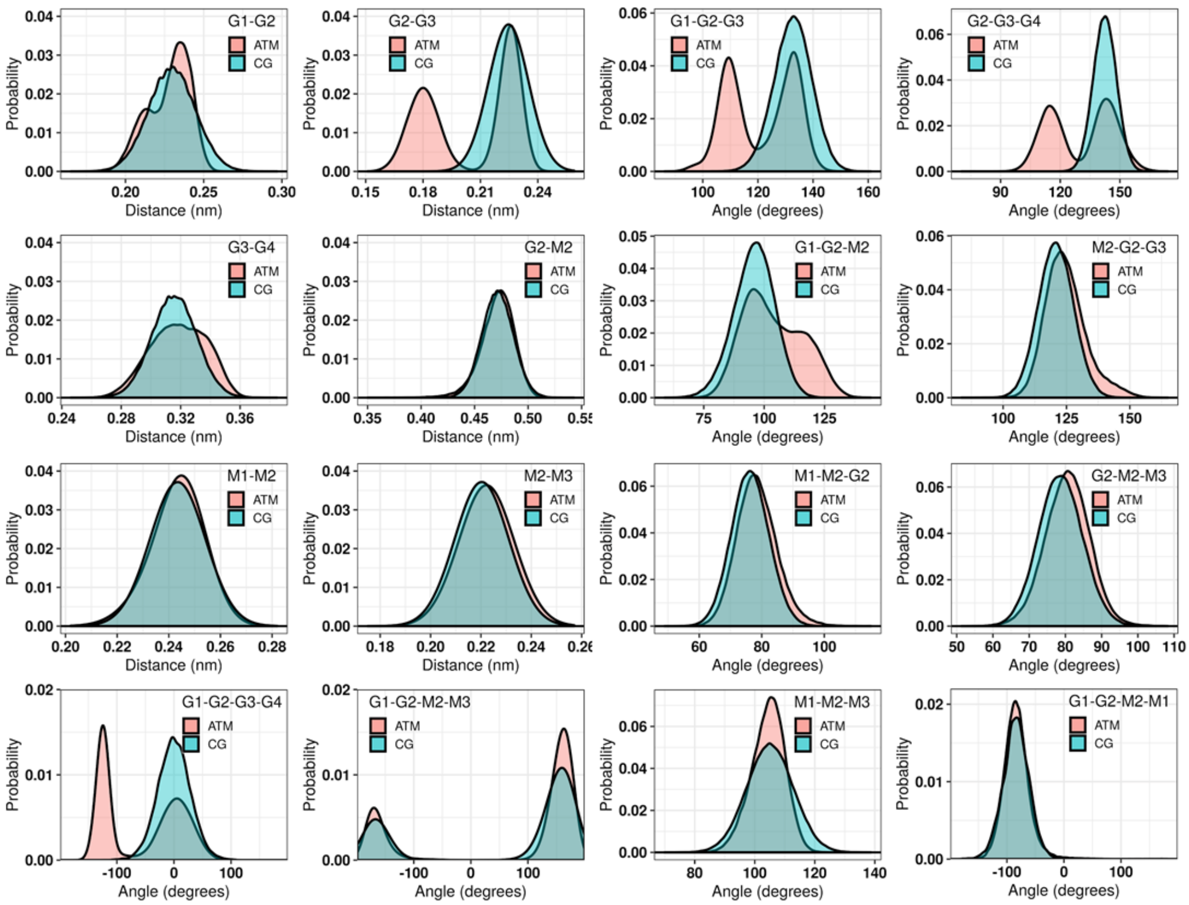

E) Mannose-α(1, 3)-[Mannose-α(1, 6)]-Mannose

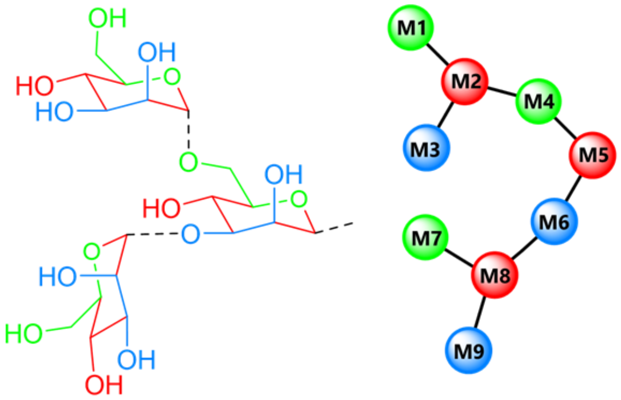

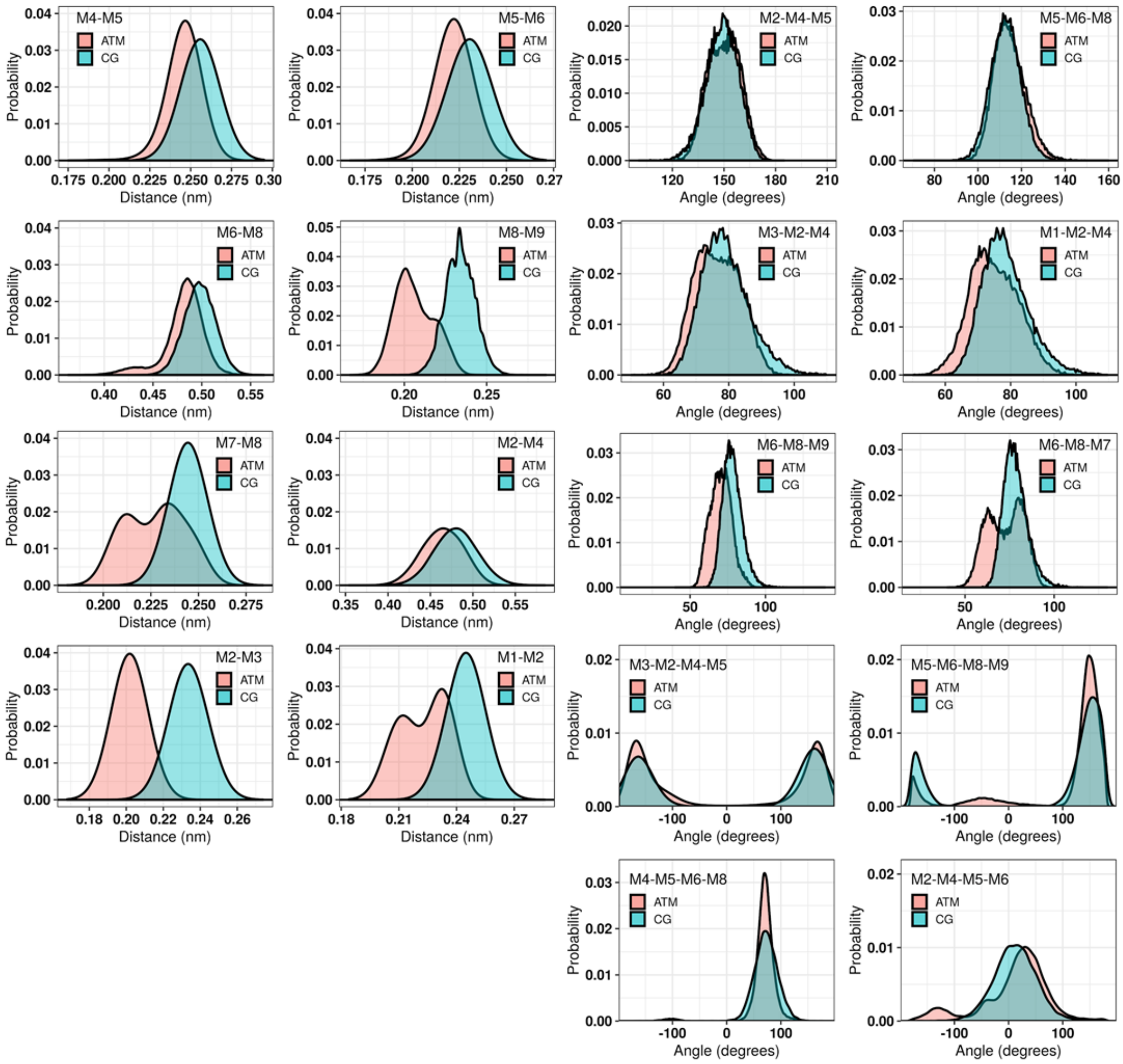

F) N-Acetylglucosamine-β(1,2)-Mannose

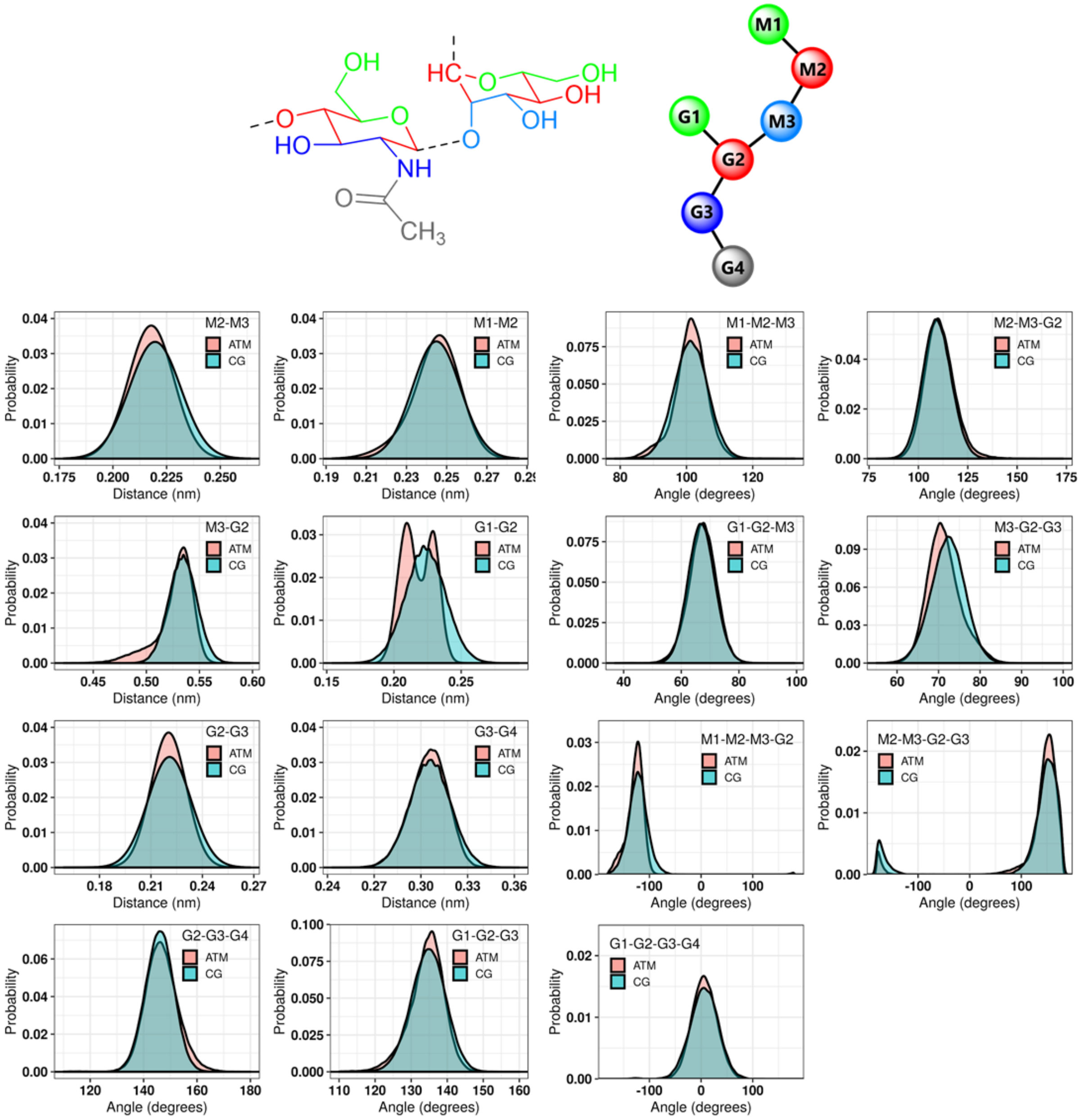

G) Galactose-β(1,4)-N-Acetylglucosamine

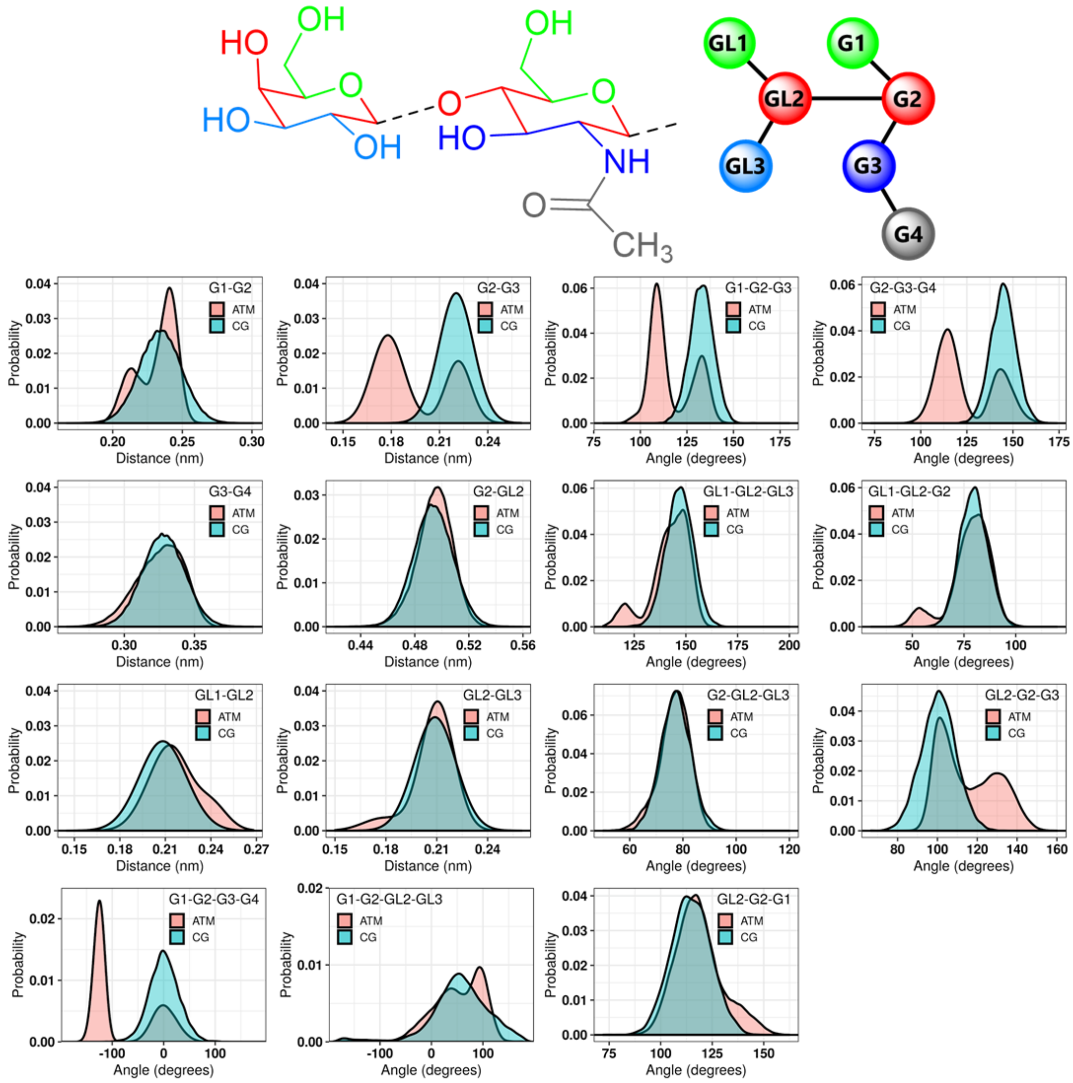

#### H) N-acetylneuraminic acid-α(2,3)-Galactose

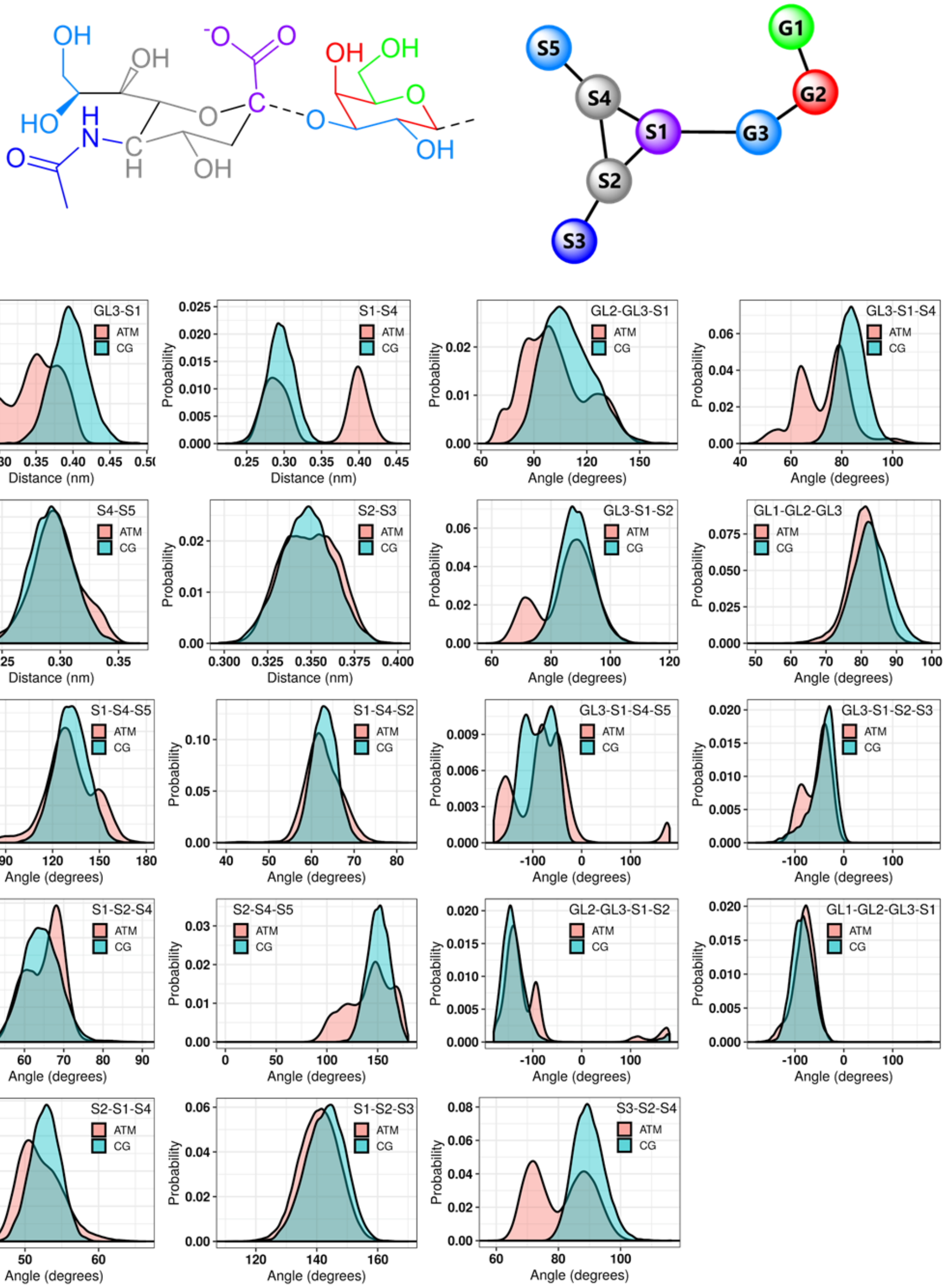

#### I) Mannose-α(1, 2)-Mannose:

(Only new bonded distributions which were not parametrized before are shown)

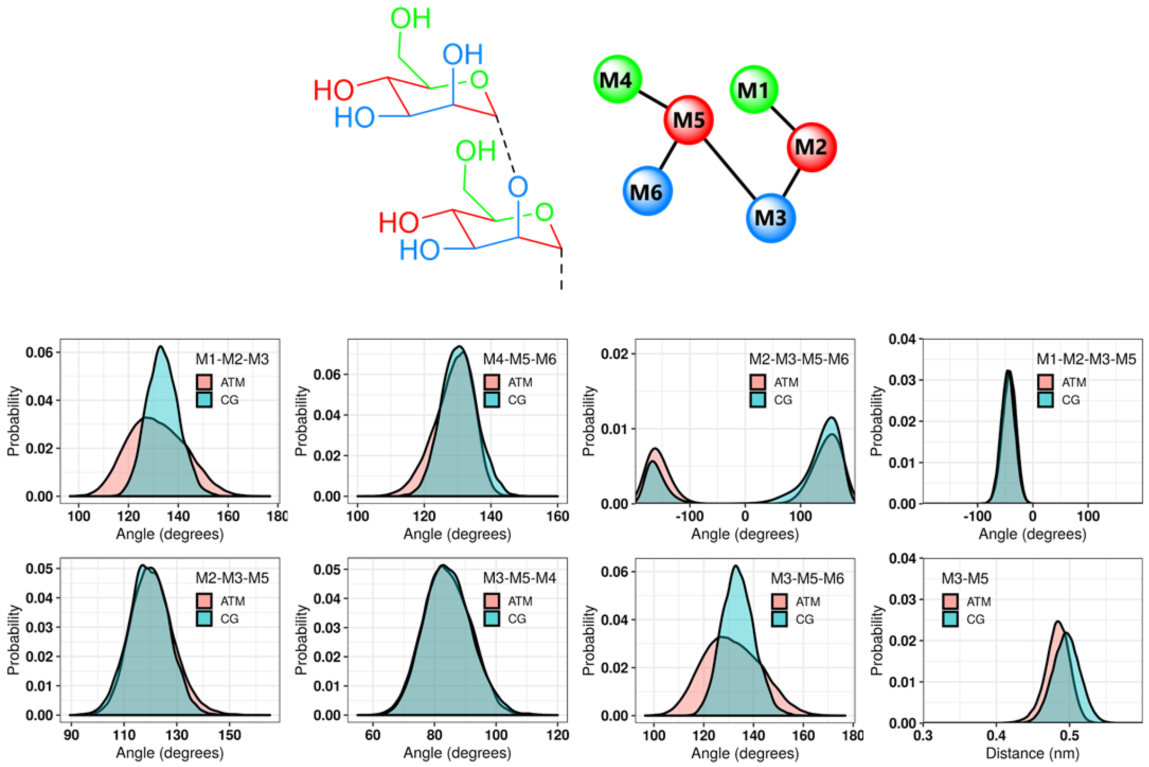

#### J) N-Acetylglucosamine-β(1, 4)-Mannose

(Only new bonded distributions which were not parametrized before are shown)

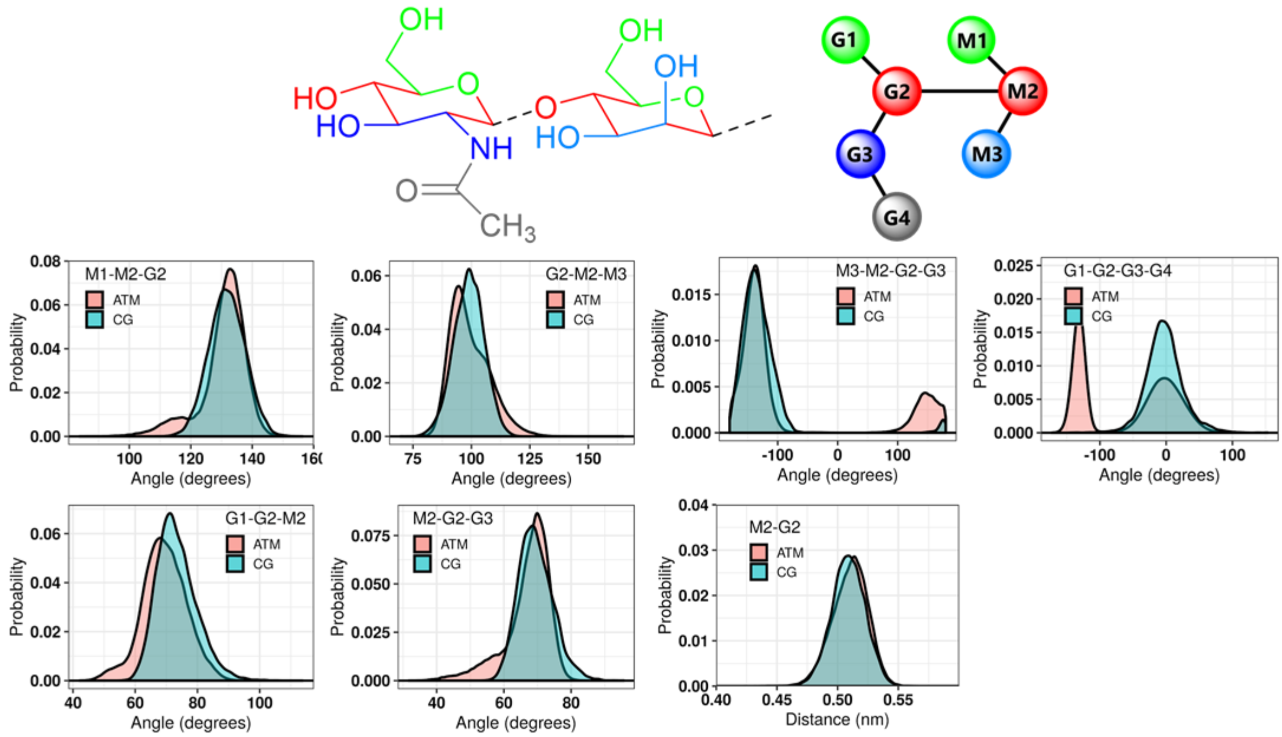

K) N-Acetylglucosamine-β(1, 6)-Mannose

(Only new bonded distributions which were not parametrized before are shown)

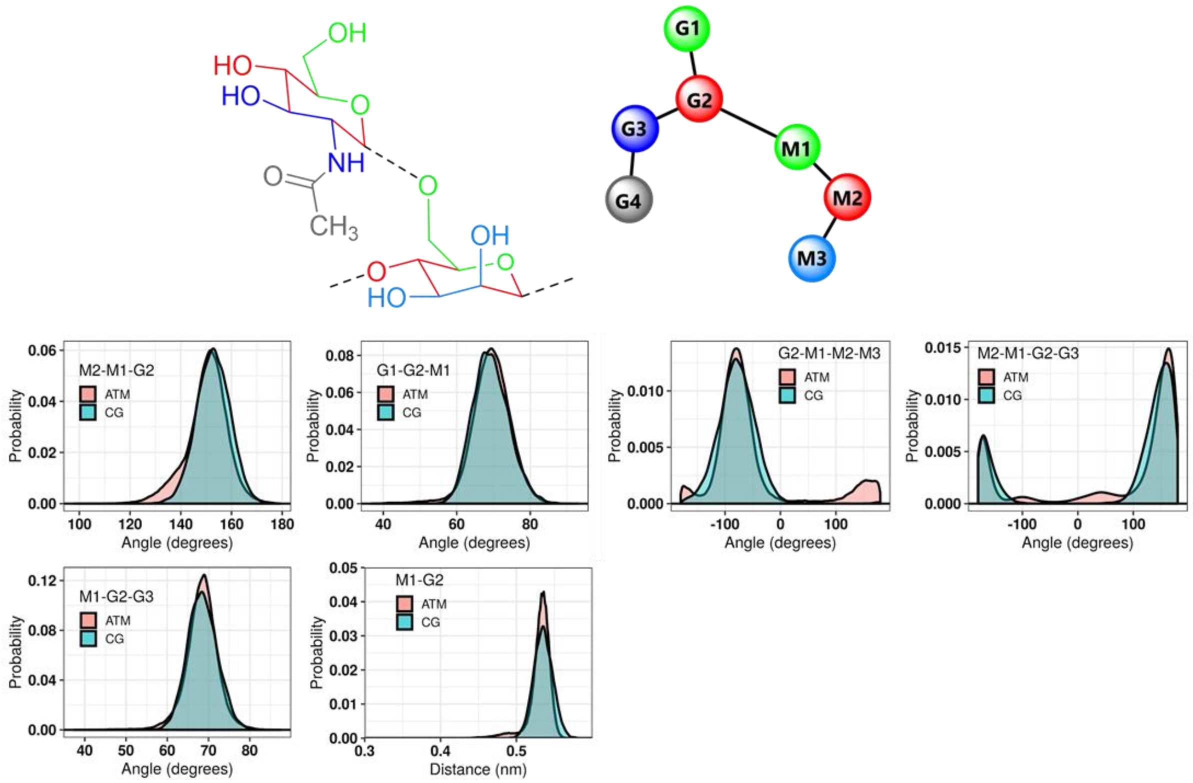

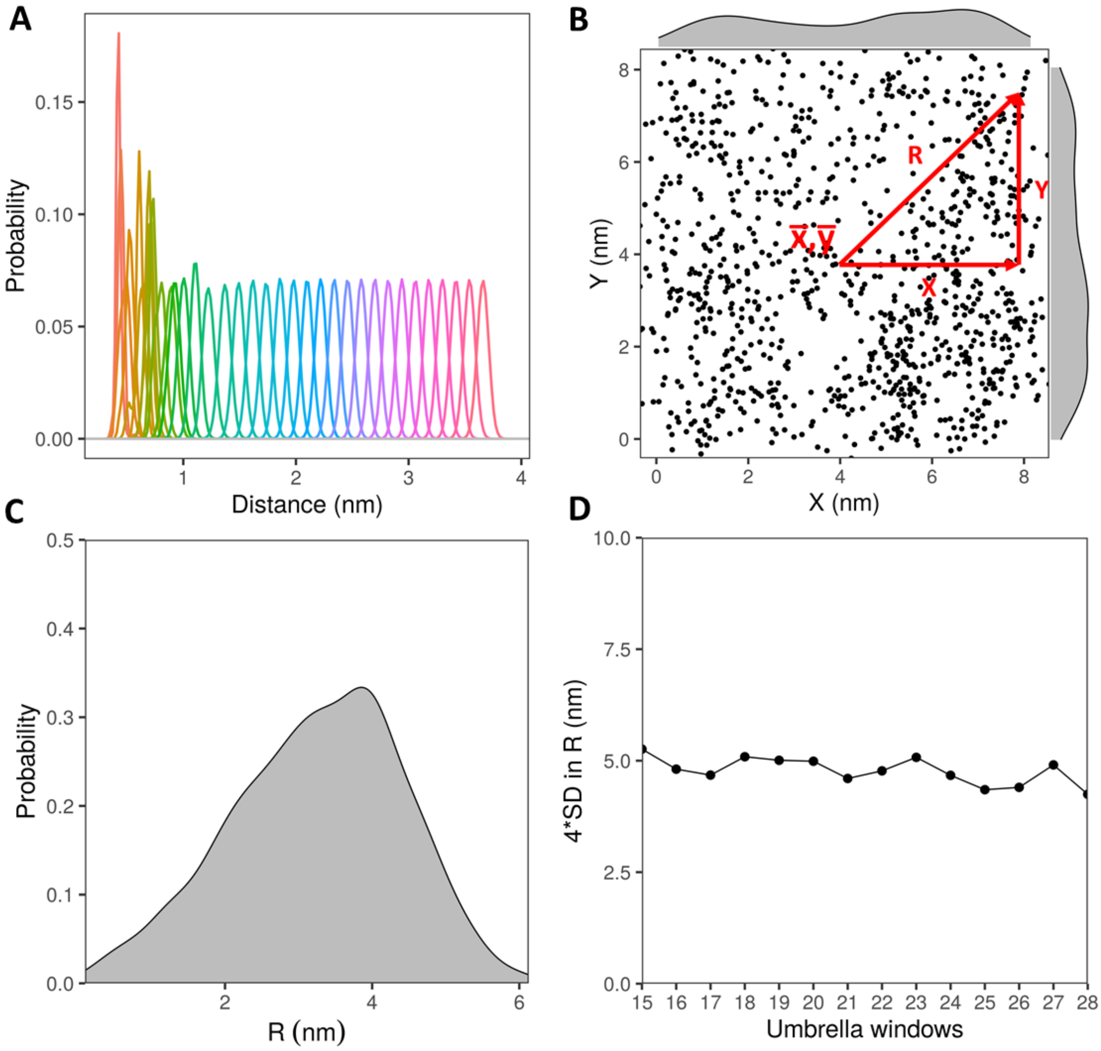

**Figure S2: Verification of ligand sampled area ($A_u$) for umbrella windows in the unbound region, using the example of Pterocarpus angolensis (PAL) interacting with M5 ligand for 0.95 scaling factor system.** (A) Overlap of umbrella sampling windows for ligand along *Z* coordinate. (B) *X* and *Y* coordinates of centre of mass of M5 ligand extracted from each frame of the final umbrella sampling window. Spread of *R* [$= \sqrt{(X^2 + Y^2)}$] was calculated from average position sampled by the ligand ($\bar{X},\ \bar{Y}$). (C) Distribution of *R* for the data in panel B (D) 95% confidence interval in *R* based on the standard deviation (SD), shown for all the windows in the unbound region. Similar ligand sampling was observed across all windows, yielding an estimate of spread equivalent to the orthogonal box area (i.e. $\pi * (4 \bullet SD)^2 \ \approx X \bullet Y$).

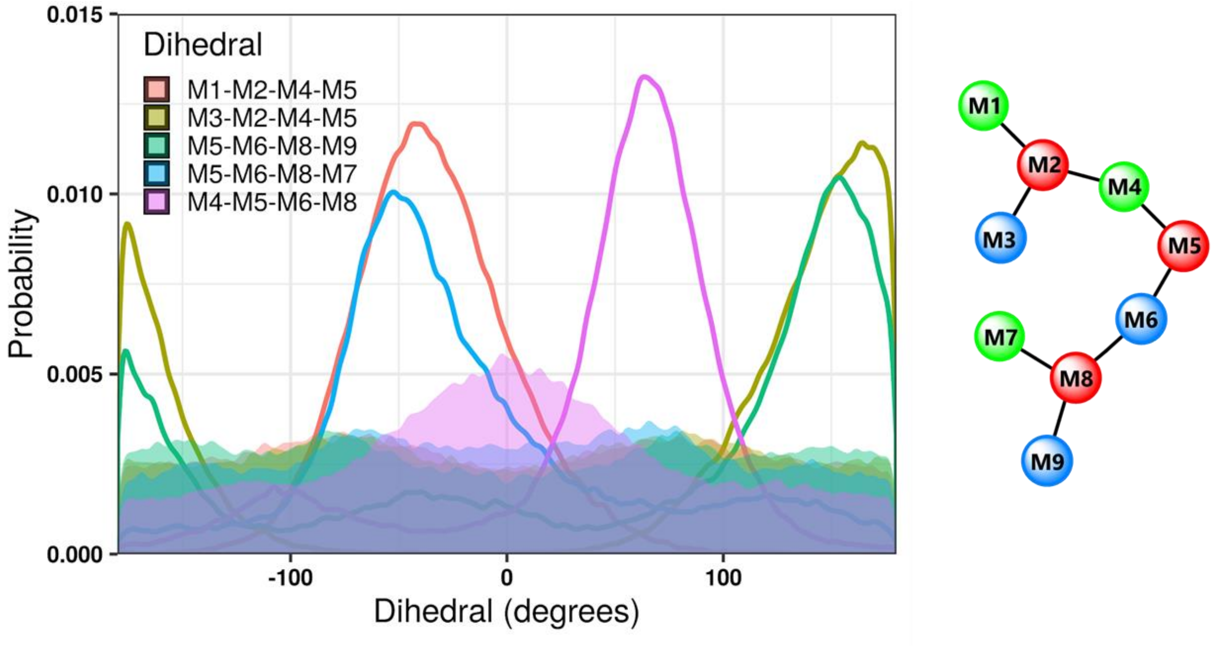

**Figure S3: M3 dihedral distributions.** Distributions obtained when dihedrals potentials are switched on (solid lines) compared to when they are switched off (filled).

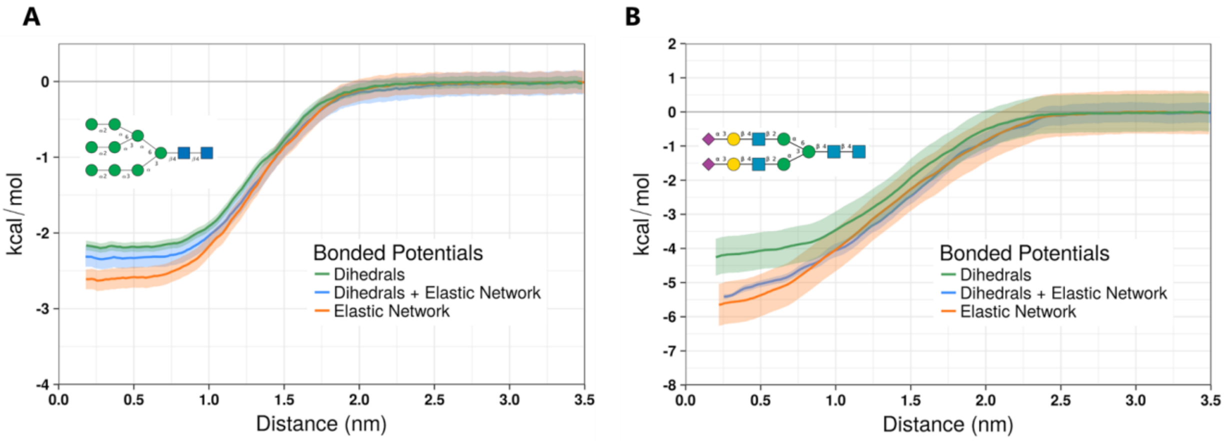

**Figure S4: Effect of elastic network on self-interaction properties.** Data are shown for (A) high mannose M9 and (B) A2G2S2 complex type glycans with dihedrals (orange), dihedrals + elastic network (green), or elastic network alone (blue). Error bars were calculated from 200 cycles of bootstrapping. Monosaccharides present in the glycans are represented by their symbolic representation, including mannose (green circle), N-acetylglucosamine (blue square), galactose (yellow circle), and Neu5Ac/sialic acid (purple diamond).

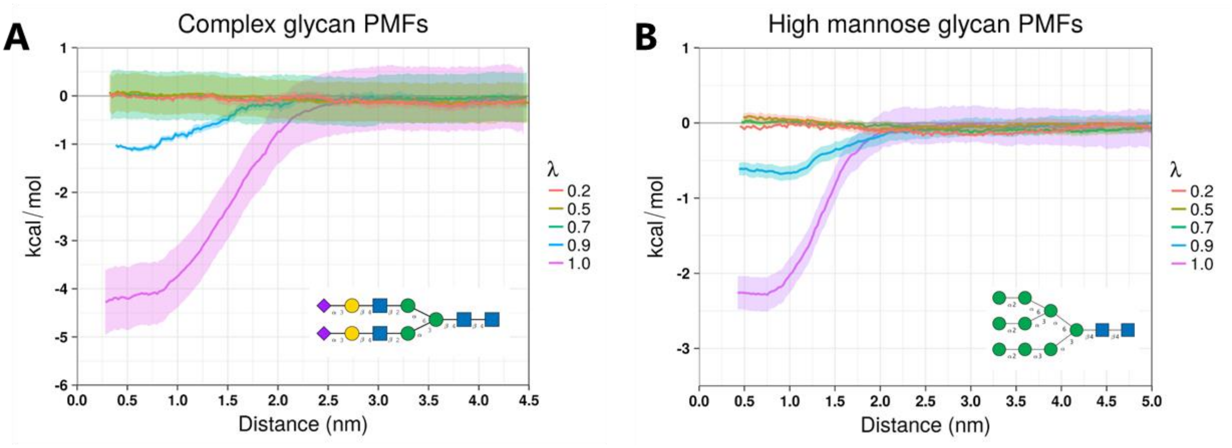

**Figure S5: Dependence of potential mean of force (PMF) on scaling the non-bonded interactions in Martini v2.2.** Monosaccharides present in the glycans are represented by their symbolic representation, including mannose (green circle), N-acetylglucosamine (blue square), galactose (yellow circle), and Neu5Ac/sialic acid (purple diamond).

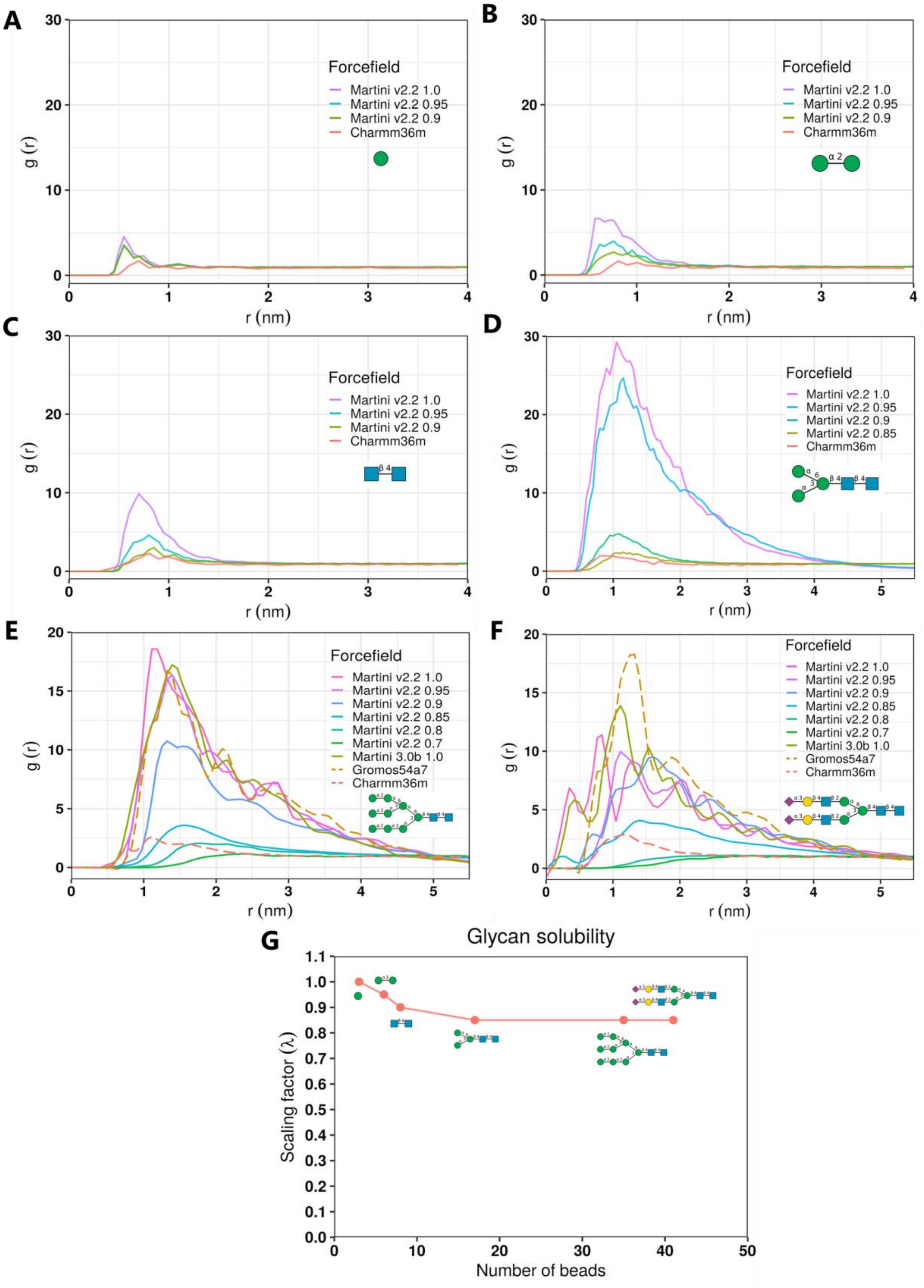

**Figure S6: Size dependence on the solubility of glycans.** Radial distribution functions (RDFs) of (A) Man, (B) Man-α12-Man, (C) GlcNAc-β14-GlcNAc, (D) M3, (E) M9, and (F) A2G2S2 glycans, calculated for Martini 2.2 and Charmm36m FFs. In (G), the scaling factor for which the RDFs match best with the Charmm36m simulations are plotted, as a function of number of beads present in the glycan.

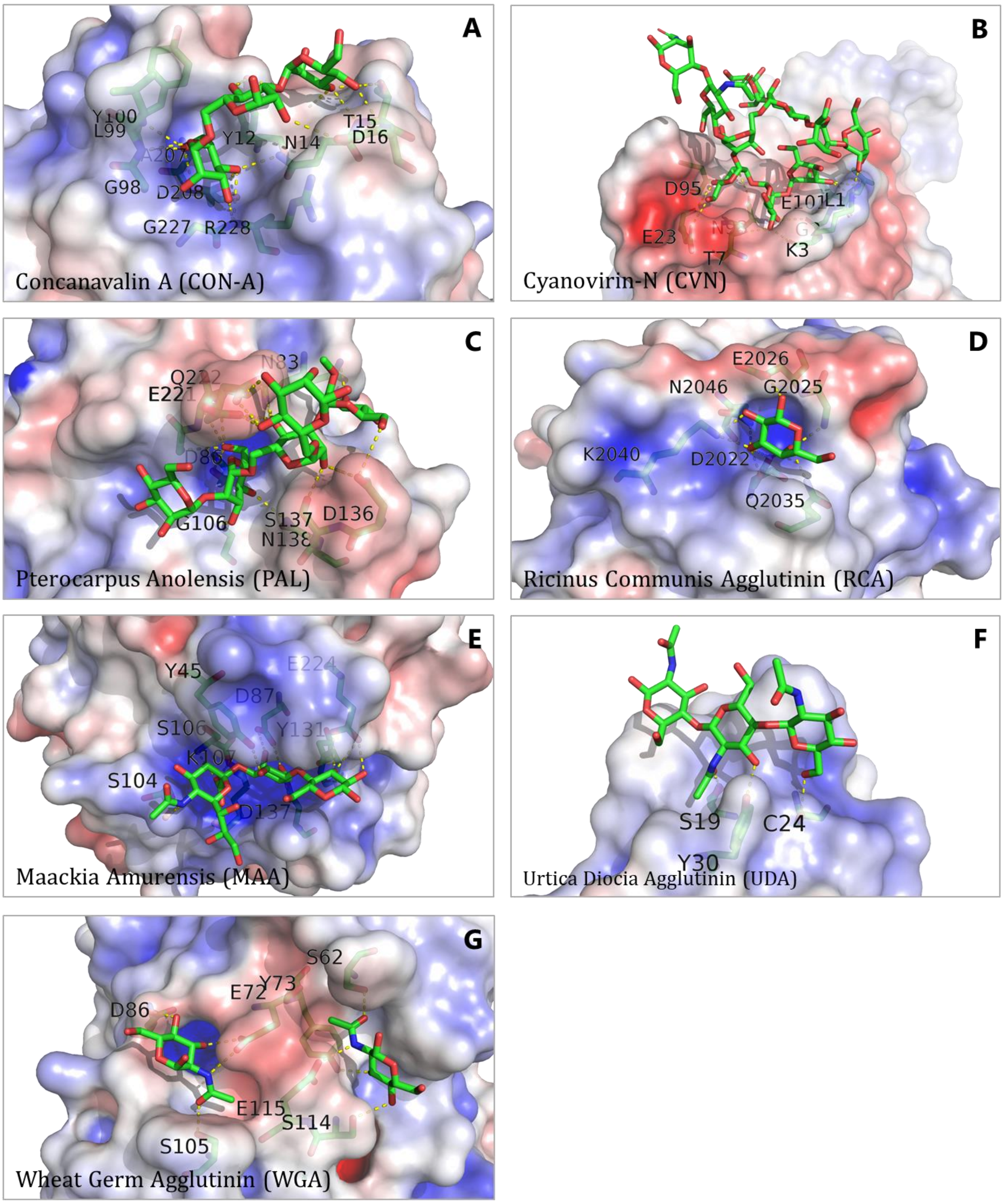

**Figure S7: Crystal structures of lectins showing the residues engaging glycans.** (A) Concanavalin A (CONA) with trimannose ($Man_3$) [PDB: 1CVN] (B) Cyanovirin-N (CVN) with high mannose (M9) [PDB: 3GXZ] (C) Pterocarpus Anolensis (PAL) with M9 [PDB: 2PHW] (D) Ricinus Communis Agglutinin (RCA) with galactose [PDB: 1RZO] (E) Maackia Amurensis (MAA) with Sialyllactose [PDB: 1DBN] (F) Urtica Diocia Agglutinin (UDA) with tri-N-Acetylchitotriose [PDB: 1EHH] (G) Wheat Germ Agglutinin (WGA) with N-Acetyl-D-glucosamine [PDB: 2UVO]. The colour coded electrostatic surface of the protein was generated using the Adaptive Poisson-Boltzmann Solver (APBS)[6] with maximum and minimum values set to +5 and -5 (units $k_B T/e$) respectively. Ligands are shown in liquorice representation.

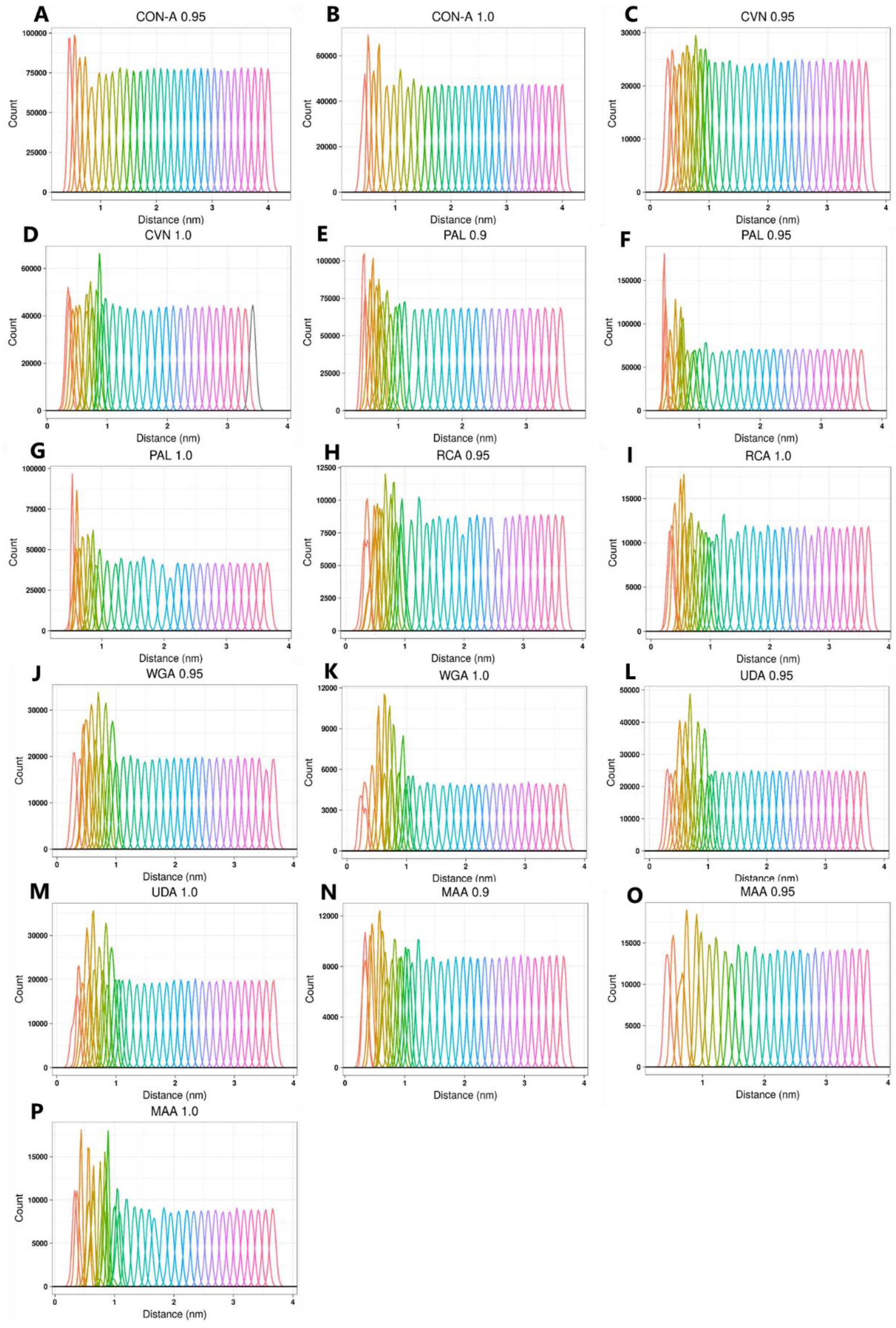

**Figure S8: Overlap of umbrella sampling windows used for calculating PMFs with different scaling factors.** Data are shown for (A)-(B) Concanavalin A (CONA), (C)-(D) Cyanovirin-N (CVN), (E)-(G) Pterocarpus Anolensis (PAL), (H)-(I) Ricinus Communis Agglutinin (RCA), (J)-(K) Wheat Germ Agglutinin (WGA), (L)-(M) Urtica Dioicia Agglutinin (UDA), and (N)-(P) Maackia Amurensis (MAA). Additional windows were added in the minima regions for better overlap of umbrella windows, where necessary.

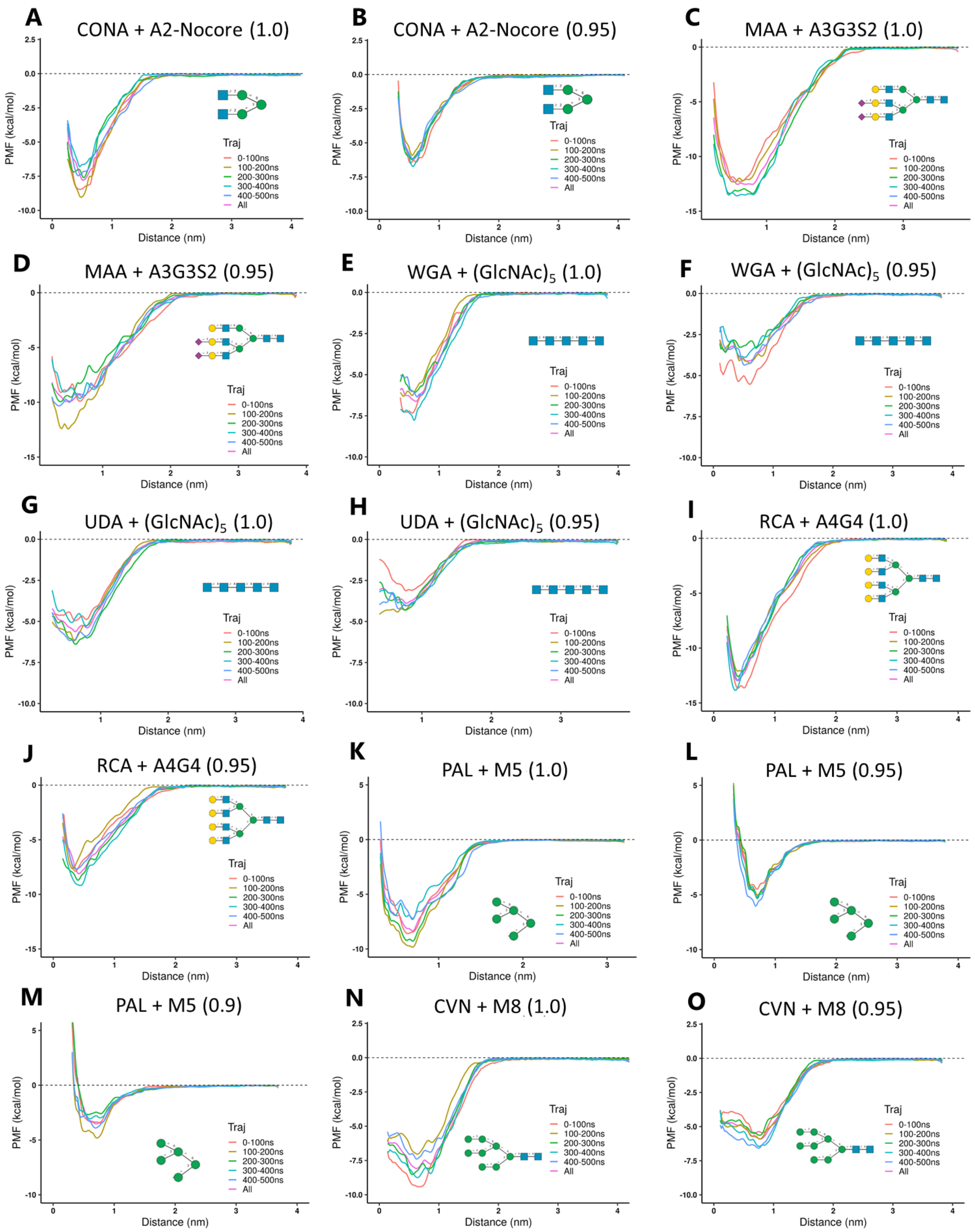

**Figure S9: Convergence analysis for the PMFs with different scaling factors.** Data are shown for (A,B) Concanavalin A (CONA), (C,D) Maackia Amurensis (MAA), (E,F) Wheat Germ Agglutinin (WGA), (G,H) Urtica Dioicia Agglutinin (UDA), (I, J) Ricinus Communis Agglutinin (RCA), (K,L,M) Pterocarpus Anolensis (PAL), (N,O) Cyanovirin-N. The glycan and the scaling factor used is denoted in the title of the plot.

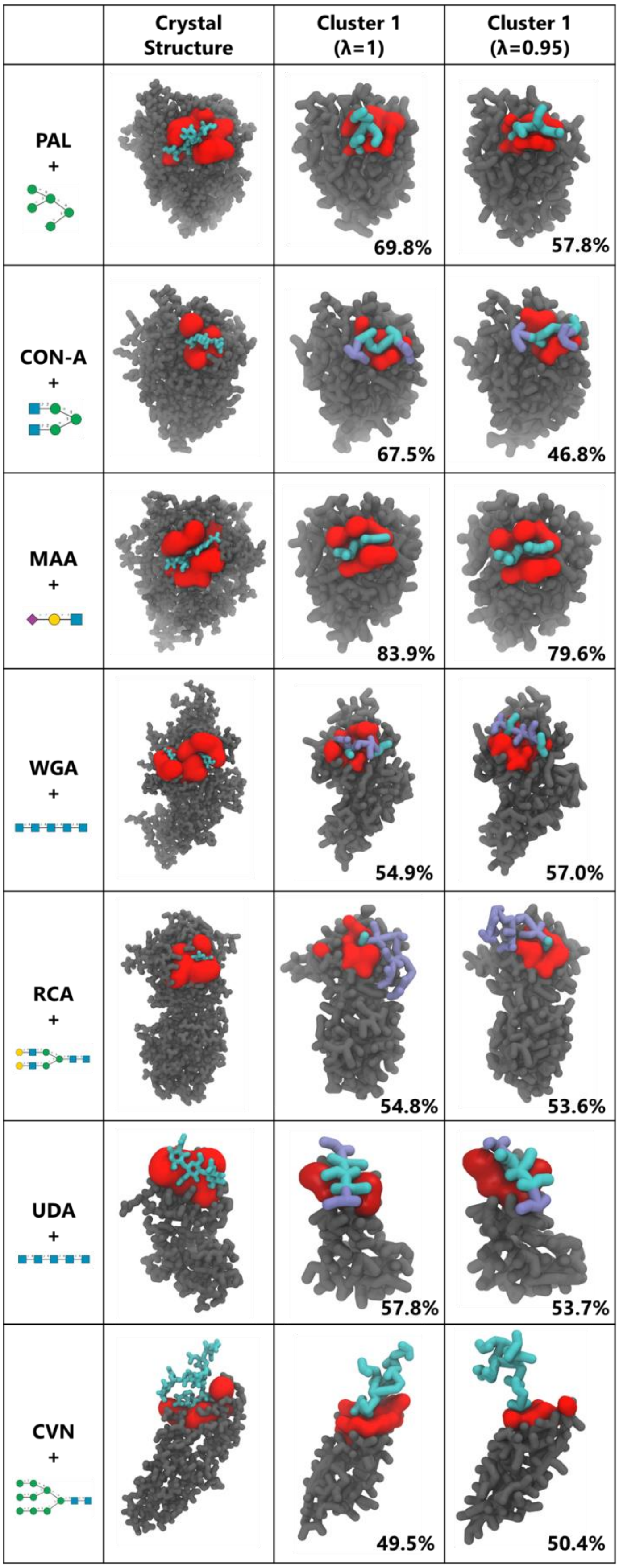

**Figure S10: Cluster analysis of lectin-glycan binding simulations.** Each lectin and its glycan binding partners are shown: Pterocarpus Anolensis (PAL) with M5;

Concanavalin A (CONA) with A2G2-Nocore;  Maackia Amurensis (MAA) with Neu5Ac-α(2,3)-Gal-β(1,4)-GlcNAc; Wheat Germ Agglutinin (WGA) with $(GlcNAc)_5$; Ricinus Communis Agglutinin (RCA) with A2G2; Urtica Dioicia Agglutinin (UDA) with $(GlcNAc)_5$; Cyanovirin-N (CVN) with M9. The top clusters from RMSD-based clustering are shown for each scaling factor (λ) and compared against the corresponding crystal structure. For every cluster the central frame of the cluster is shown. For each protein, the lectin binding region is shown in surface representation while the remainder is shown in licorice representation. Glycans are sown in licorice representation, with fragments resolved or unresolved in each crystal structure coloured in cyan or blue, respectively. Cluster populations are noted in the right bottom corner of each cluster snapshot. Monosaccharides present in the glycans for each system are represented by their symbolic representation: mannose (green circle), N-acetylglucosamine (blue square), galactose (yellow circle), and Neu5Ac/sialic acid (purple diamond)

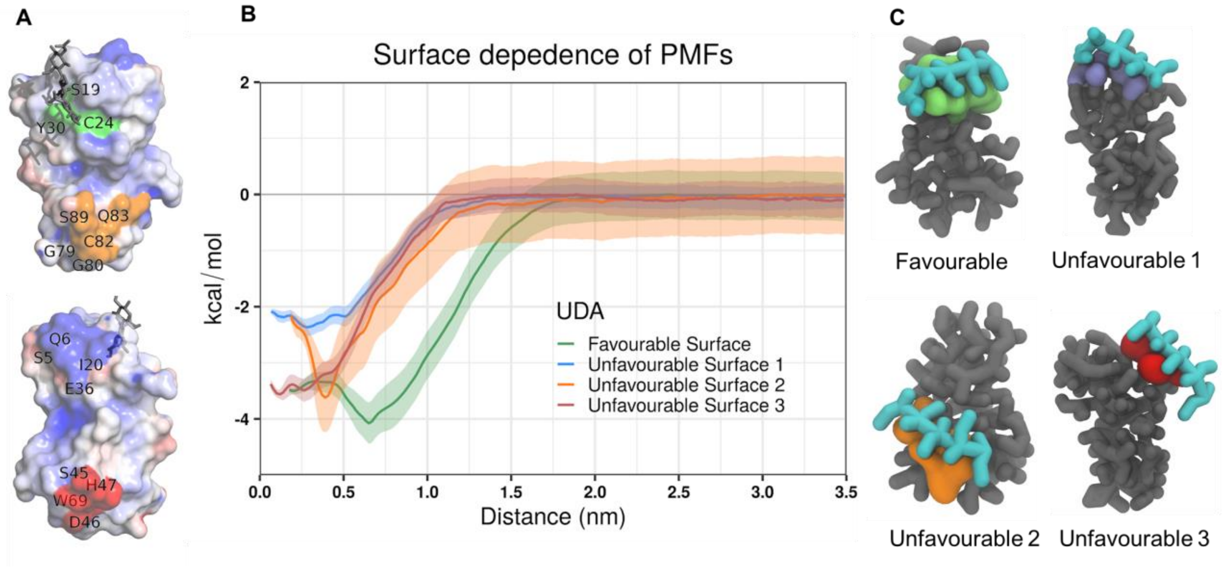

**Figure S11: Surface dependence of PMFs for (GlcNAc)5 binding to Urtica dioica agglutinin. (**A) ATM representation of different binding surfaces: “favourable” native (green), “unfavourable 1” (blue), “unfavourable 2” (orange), and “unfavourable 3” (red) surfaces. (B) PMFs for binding to the different protein surfaces. (C) Snapshots of the different protein surfaces bound to $(GlcNAc)_5$ in CG representation, which were used for the umbrella sampling simulations.

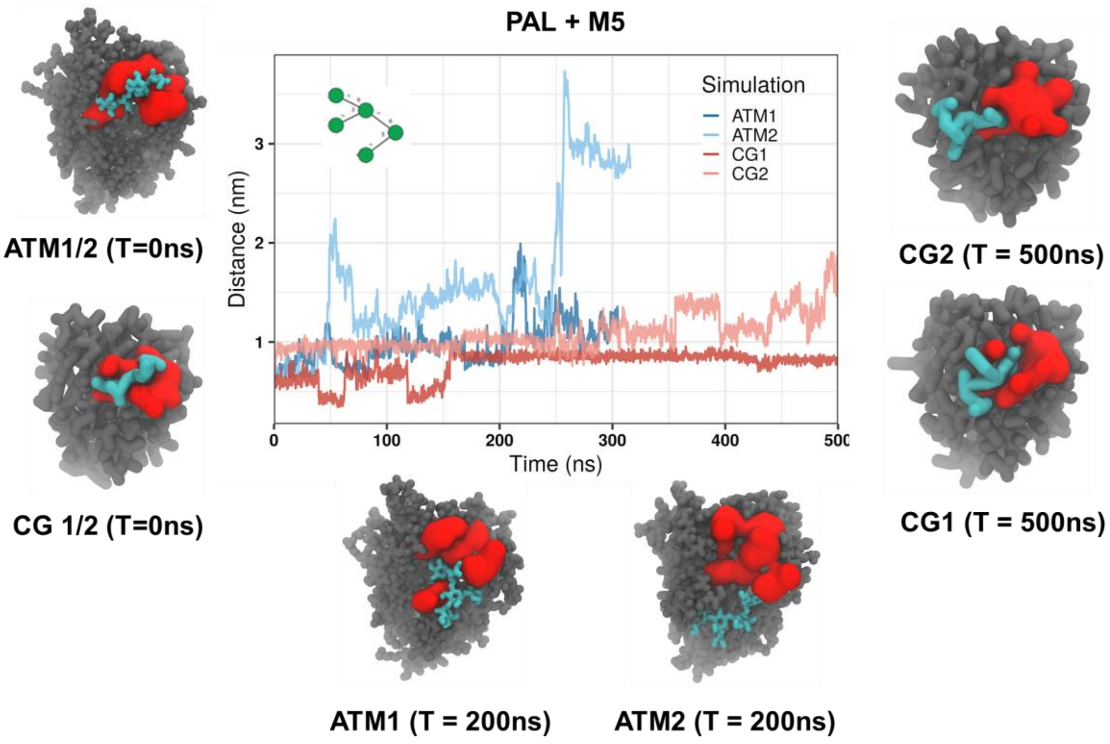

**Figure S12: Unrestrained simulations of Pterocarpus angolensis lectin (PAL) and M5 ligand (low affinity case).** Two replicates of each CG and ATM simulation were run without any umbrella restraining potential. Centre of mass distance between the ligand and the protein pocket throughout the simulations are plotted, and snapshots of different bound states during the simulations are shown alongside.

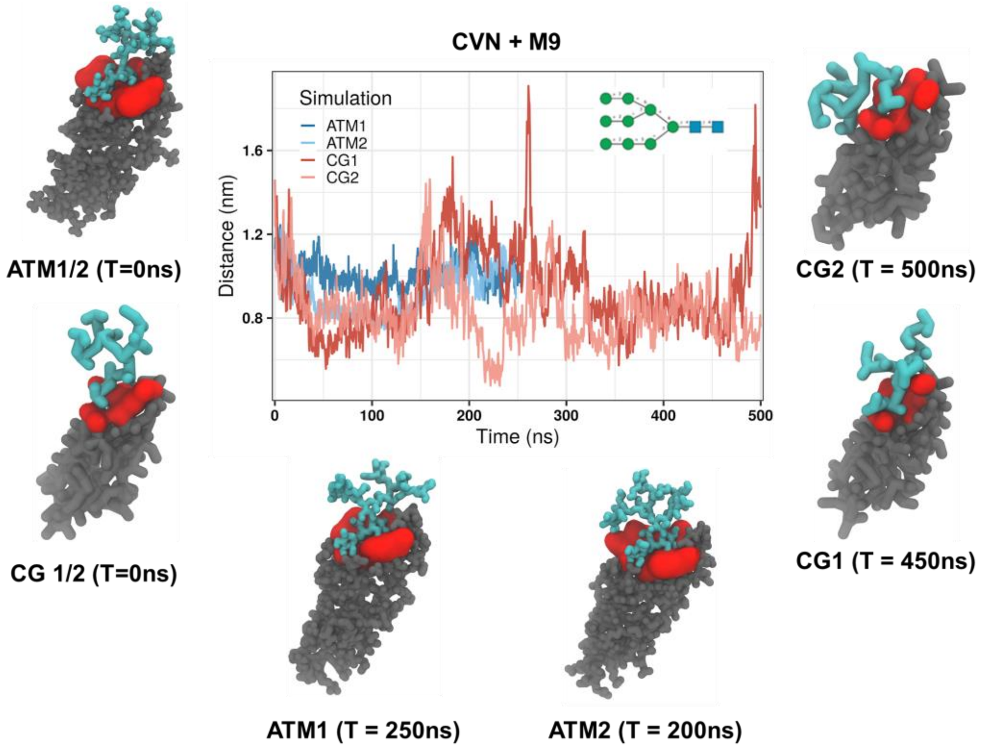

**Figure S13: Unrestrained simulations of Cyanovirin (CVN) and M9 ligand (high affinity case). T**wo replicates of each CG and ATM simulation were run without any umbrella restraining potential. Centre of mass distance between the ligand and the protein pocket throughout the simulations are plotted, and snapshots of different bound states during the simulations are shown alongside.

#### References

1. Shenoy, S. R., O'Keefe, B. R., Bolmstedt, A. J., Cartner, L. K. & Boyd, M. R. Selective interactions of the human immunodeficiency virus-inactivating protein cyanovirin-N with high-mannose oligosaccharides on gp120 and other glycoproteins. *J. Pharmacol. Exp. Ther.* **297,** 704–710 (2001).
2. Garcia-Pino, A., Buts, L., Wyns, L., Imberty, A. & Loris, R. How a plant lectin recognizes high mannose oligosaccharides. *Plant Physiol.* **144,** 1733–1741 (2007).
3. Haseley, S. R., Talaga, P., Kamerling, J. P. & Vliegenthart, J. F. G. Characterization of the carbohydrate binding specificity and kinetic parameters of lectins by using surface plasmon resonance. *Anal. Biochem.* **274,** 203–210 (1999).
4. Shinohara, Y. *et al.* Use of a biosensor based on surface plasmon resonance and biotinyl glycans for analysis of sugar binding specificities of lectins. *J. Biochem.* **117,** 1076–82 (1995).
5. Mandal, D. K., Kishore, N. & Brewer, C. F. Thermodynamics of Lectin-Carbohydrate Interactions. Titration Microcalorimetry Measurements of the Binding of N-Linked Carbohydrates and Ovalbumin to Concanavalin A. *Biochemistry* **33,** 1149–1156 (1994).
6. Baker, N. A., Sept, D., Joseph, S., Holst, M. J. & McCammon, J. A. Electrostatics of nanosystems: application to microtubules and the ribosome. *Proc. Natl. Acad. Sci. U. S. A.* **98,** 10037–41 (2001).
